# Compact engineered human mechanosensitive transactivation modules enable potent and versatile synthetic transcriptional control

**DOI:** 10.1101/2022.03.21.485228

**Authors:** Barun Mahata, Alan Cabrera, Daniel A. Brenner, Rosa Selenia Guerra-Resendez, Jing Li, Jacob Goell, Kaiyuan Wang, Yannie Guo, Mario Escobar, Abinand Krishna Parthasarathy, Hailey Szadowski, Guy Bedford, Daniel Reed, Isaac B. Hilton

**Affiliations:** Department of Bioengineering, Rice University, Houston, TX, USA; Systems, Synthetic, and Physical Biology Graduate Program, Rice University, Houston, TX, USA; Department of BioSciences, Rice University, Houston, TX, USA

## Abstract

Engineered transactivation domains (TADs) combined with programmable DNA binding platforms have revolutionized synthetic transcriptional control. Despite recent progress in programmable CRISPR/Cas-based transactivation (CRISPRa) technologies, the TADs used in these systems often contain poorly tolerated elements and/or are prohibitively large for many applications. Here we defined and optimized minimal TADs built from human mechanosensitive transcription factors (MTFs). We used these components to construct potent and compact multipartite transactivation modules (MSN, NMS, and eN3×9) and to build the CRISPR-<u>d</u>Cas9 recruited <u>e</u>nhanced <u>a</u>ctivation <u>m</u>odule (CRISPR-DREAM) platform. We found that CRISPR-DREAM was specific, robust across mammalian cell types, and efficiently stimulated transcription from diverse regulatory loci. We also showed that MSN and NMS were portable across Type I, II, and V CRISPR systems, TALEs, and ZF proteins. Further, as proofs of concepts, we used dCas9-NMS to efficiently reprogram human fibroblasts into iPSCs and demonstrated that MTF TADs are efficacious and well tolerated in therapeutically important primary human cell types. Finally, we leveraged the compact and potent features of these engineered TADs to build new dual and all-in-one CRISPRa AAV systems. Altogether, these compact human TADs, fusion modules, and new delivery architectures should be valuable for synthetic transcriptional control in biomedical applications.

## Introduction

Nuclease deactivated CRISPR-Cas (dCas) systems can be used to modulate transcription in cells and organisms^1–8^. For CRISPR-based activation (CRISPRa) approaches, transcriptional activators can be recruited to genomic regulatory elements using direct fusions to dCas proteins^9–13^, antibody-mediated recruitment^14^, or using engineered gRNA architectures^15, 16^. High levels of CRISPRa-driven transactivation have been achieved by shuffling^17^, reengineering^18^, or combining^9, 19, 20^ transactivation domains (TADs) and/or chromatin modifiers. However, many of the transactivation components used in these CRISPRa systems have coding sizes that are restrictive for applications such as viral vector-based delivery. Moreover, most of the transactivation modules that display high potencies harbor components derived from viral pathogens and can be poorly tolerated in clinically important cell types, embryos, and animal models, which could hamper biomedical or *in vivo* use^21–23^. Finally, there is an untapped repertoire of thousands of human transcription factors (TFs) and chromatin modifiers^24–27^ that has yet to be systematically tested and optimized as programmable transactivation components across endogenous target sites, DNA binding platforms, and recruitment architectures (e.g. direct protein fusions vs. aptamer-based). This diverse repertoire of human protein building blocks could be used to reduce the size of transactivation components, obviate the use of viral TFs, and possibly permit cell and/or pathway specific transactivation.

Mechanosensitive transcription factors (MTFs) modulate transcription in response to mechanical cues and/or external ligands^28, 29^. When stimulated, MTFs are shuttled into the nucleus where they can rapidly transactivate target genes by engaging key nuclear factors including RNA polymerase II (RNAP) and/or histone modifiers^30–33^. The dynamic shuttling of MTFs can depend upon both the nature and the intensity of stimulation. Mammalian cells encode several classes of MTFs, including serum regulated MTFs (e.g., YAP, TAZ, SRF, MRTF-A and B, and MYOCD)^29, 34^, cytokine regulated/JAK-STAT family MTFs (e.g., STAT proteins)^35^, and oxidative stress/antioxidant regulated MTFs (e.g., NRF2)^36^; each of which can potently activate transcription when appropriately stimulated. The robust, highly orchestrated, and relatively ubiquitous gene regulatory effects of these classes of human MTFs make them excellent potential sources of new non-viral TADs that could be leveraged as components of engineered CRISPRa systems and/or other synthetic gene activation platforms.

Here, we quantify the endogenous transactivation potency of dozens of different TADs derived from human MTFs in different combinations and across various dCas-based recruitment architectures. We use these data to design new multipartite transactivation modules, called MSN, NMS, and eN3×9 and we further apply the MSN and NMS effectors to build the CRISPR-<u>d</u>Cas9 recruited <u>e</u>nhanced <u>a</u>ctivation <u>m</u>odule (DREAM) platform. We demonstrate that CRISPR-DREAM potently stimulates transcription in primary human cells and cancer cell lines, as well as in murine and CHO cells. We also show that CRISPR-DREAM activates different classes of RNAs spanning diverse regulatory elements within the human genome. Further, we find that the MSN/NMS effectors are portable to smaller engineered dCas9 variants, natural orthologues of dCas9, dCas12a, Type I CRISPR/Cas systems, and TALE and ZF proteins. Moreover, we demonstrate that a dCas12a-NMS fusion enables superior multiplexing transactivation capabilities compared to existing systems.

We also show that dCas9-NMS efficiently reprograms human fibroblasts and we leverage the compact size of these new effectors to build potent dual and all-in-one CRISPRa AAVs. Finally, we demonstrate that MSN, NMS, and eN3×9 are better tolerated than viral-based TADs in primary human MSCs and T cells. Overall, the engineered transactivation modules that we have developed here are small, highly potent, devoid of viral sequences, versatile across programmable DNA binding systems, and enable robust multiplexed transactivation in human cells – important features that can be leveraged to test new biological hypotheses and engineer complex cellular functions.

## Results

### Select TADs from MTFs can activate transcription from diverse endogenous human loci when recruited by dCas9

We first isolated TADs from 7 different serum-responsive MTFs (YAP, YAP-S397A^37^, TAZ, SRF, MRTF-A, MRTF-B, and MYOCD) and analyzed their ability to activate transcription when recruited to human promoters using either N- or C-terminal fusion to *Streptococcus pyogenes* dCas9 (dCas9), SunTag-mediated recruitment^14^, or recruitment via a gRNA aptamer and fusion to the MCP protein^15^ (**Supplementary Fig. 1**). TADs derived from MRTF-A, MRTF-B, or MYOCD displayed consistent transactivation potential across recruitment architectures. We next compared the optimal recruitment strategies for MRTF-A and MRTF-B TADs because they were more potent than, or comparable to, the MYOCD TAD, yet slightly smaller. Our results demonstrated that TADs from MRTF-A and B functioned best when fused to the MCP protein and recruited via gRNA aptamers (**Supplementary Fig. 2**), and further that this strategy could be used with pools or single gRNAs, and to activate enhancer RNAs (eRNAs) and long noncoding RNAs (lncRNAs).

Although the NRF2-ECH homology domains 4 and 5 (Neh4 and Neh5, respectively) within the oxidative stress/antioxidant regulated NRF2 human MTF have been shown to activate gene expression in Gal4 systems^30^, we observed that neither Neh4 nor Neh5 were capable of potent human gene activation when recruited to promoters in any dCas9-based architecture (**Supplementary Fig. 3**). Therefore, we constructed an engineered TAD called eNRF2, consisting of Neh4 and Neh5 separated by an extended glycine-serine linker and found that the eNRF2 TAD stimulated high levels of transactivation in all dCas9-based recruitment configurations (**Supplementary Fig. 3**). Similar to the MRTF-A/B TADs, eNRF2 displayed optimal potency in the gRNA aptamer/MCP-based recruitment architecture and transactivated diverse human regulatory loci (**Supplementary Fig. 4**). We next tested whether TADs derived from one of 6 different cytokine regulated/JAK-STAT family MTFs (STAT1 – 6) could transactivate human genes but observed that single STAT TADs alone were incapable of potent transactivation regardless of dCas9-based recruitment context (**Supplementary Fig. 5**). Interestingly, all native TADs that displayed measurable efficacy at endogenous target sites harbored at least one 9aa segment that perfectly matched predictions generated by previously described software^38^ (**Supplementary Fig. 6**). Together, these data demonstrate that TADs from human MTFs can transactivate human loci when recruited via dCas9 and that these TADs are amenable to protein engineering.

### Combinations of TADs from MTFs can potently activate human genes when recruited by dCas9

STAT proteins typically activate gene expression in combination with co-factors^39^. Therefore, we tested if TADs from different STAT proteins might synergize with other MTF TADs. We built 24 different bipartite fusion proteins by linking each STAT TAD to the N- or C-terminus of either the MRTF-A or MRFT-B TAD and then assayed the relative transactivation potential of each bipartite fusion when recruited to the human *OCT4* promoter using gRNA aptamer/MCP-based recruitment (**Supplementary Fig. 7**). All of these 24 fusions markedly outperformed single TADs from MRTF-A/B or STAT alone, and one bipartite TAD configuration (MRFT-A/STAT1) was comparable to MCP fused to the dCas9-SAM derived bipartite p65-HSF1^15^ module. Similar results were obtained at other human promoters (**Supplementary Fig. 7d and e**).

We next investigated whether the eNRF2 TAD could further enhance the potency of the MRFT-A/STAT1 module by building tripartite fusions consisting of MRTF-A/STAT1/eNRF2 (MSN) or eNRF2/MRTF-A/STAT1 (NMS) TADs. Both MSN and NMS stimulated *OCT4* mRNA synthesis to levels comparable to the state-of-the-art CRISPRa platforms (**Supplementary Fig. 8a and b**) when recruited to the *OCT4* promoter using gRNA aptamers/MCP-based targeting. We further validated these results against dCas9 + MCP-p65-HSF1 at six other endogenous promoters (**Supplementary Fig. 8c and d**). Notably, switching the order of p65 and HSF1 did not improve the efficacy of the p65-HSF1 TAD module (**Supplementary Fig. 8e and f**). Surprisingly, the potency of the MSN tripartite effector was not further enhanced by the direct fusion of other TADs to the C-terminus of dCas9 (**Supplementary Fig. 8g**). Collectively, these data show that gRNA aptamer/MCP-based recruitment of the MSN or NMS modules – termed the CRISPR-<u>d</u>Cas9 recruited <u>e</u>nhanced <u>a</u>ctivation <u>m</u>odule (DREAM) platform – can efficiently stimulate transcription without viral components. Our results also demonstrate that natural and engineered human TADs can have non-obvious interactions when combinatorially recruited in bi- and tripartite fashions.

### CRISPR-DREAM displays potent activation of endogenous promoters, is specific, and is robust across diverse mammalian cell types

To assess the relative transactivation potential of CRISPR-DREAM, we first targeted the DREAM or SAM^15^ systems (**Fig. 1a, 1b**), to different human promoters in HEK293T cells. All components for both the DREAM and SAM systems were well-expressed in HEK293T cells (**Fig. 1c**). At all promoters targeted using pools of gRNAs (n = 15), DREAM was superior or comparable to the SAM system (**Fig. 1d and Supplementary Fig. 9**). Similar results were obtained using antibody staining of CD34 protein levels and flow cytometry analyses in single cells after DREAM or the SAM system (or dCas9-VPR^9^/dCas9 + MCP-VPR) was targeted to the CD34 promoter (**Supplementary Fig. 10**). Additionally, when human promoters were targeted using only single gRNAs (n = 11), DREAM remained superior or comparable to the SAM system in all experiments (**Fig. 1e and Supplementary Fig. 11**). Interestingly, this trend extended throughout ∼1kb upstream of the transcription start sites (TSSs) surrounding human genes (**Supplementary Fig. 12**). Collectively, these data demonstrate that, although the DREAM system is smaller than the SAM system, and is devoid of viral TADs, it displays superior or comparable transactivation potency in human cells.

**Fig. 1.**
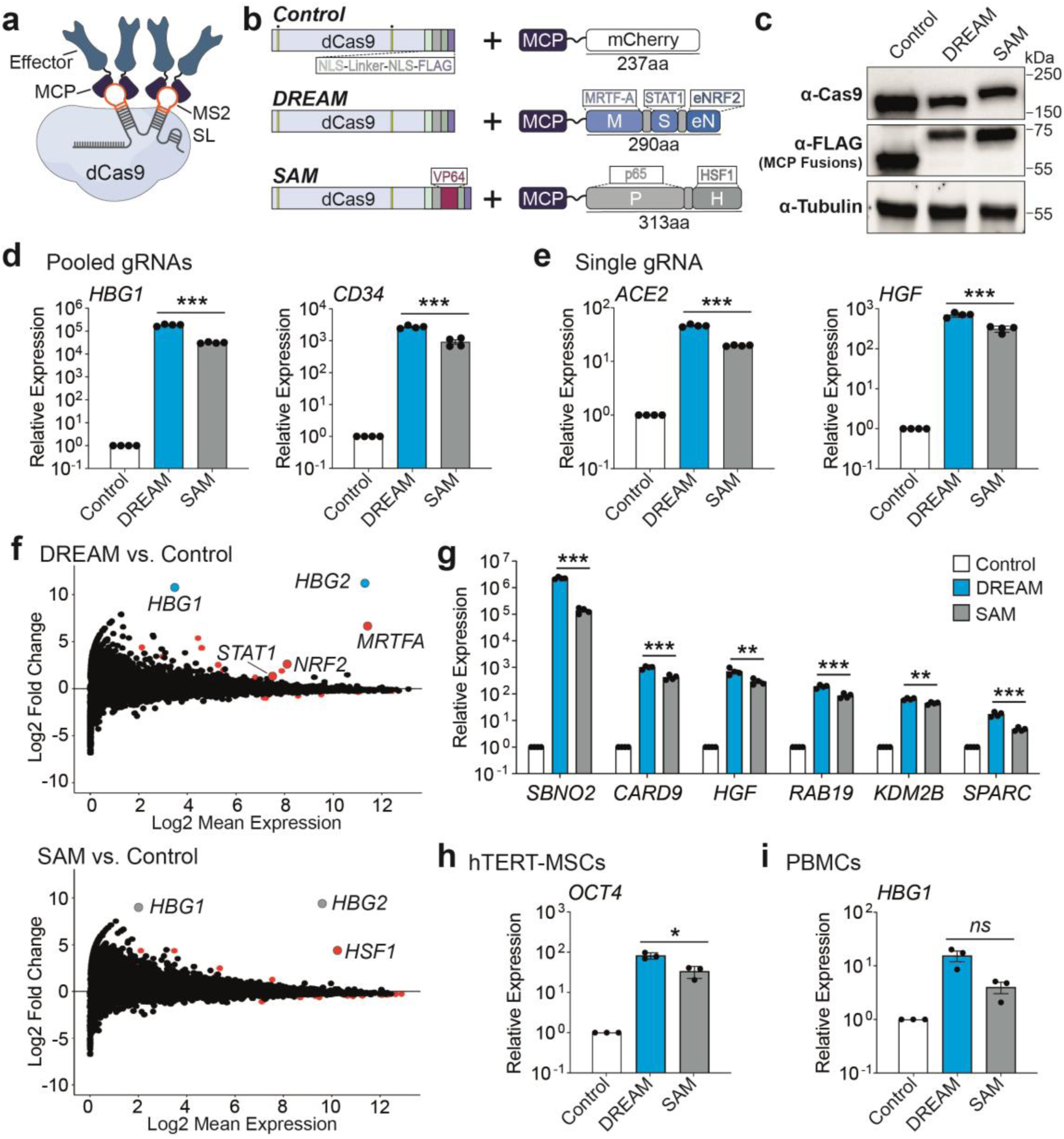
CRISPR-DREAM displays potent activation at human promoters, has high specificity, and is robust across cell types. **a**. Nuclease inactivated *Streptococcus pyogenes* dCas9 (dCas9), a gRNA containing two engineered MS2 stem-loops (MS2 SLs) and MS2 binding Cap Protein (MCP)-fused transcriptional effector proteins are schematically depicted. Nuclease-inactivating mutations (D10A and H840A) are indicated by yellow bars with dots above. **b**. dCas9 and MCP-fusion proteins, including an MCP-mCherry fusion (Control; top), the engineered tripartite MCP-MSN domain fusion (DREAM system; middle), and dCas9-VP64 and the MCP-p65-HSF1 fusion protein (SAM system; bottom) are schematically depicted. **c**. The expression levels of dCas9 and dCas9-VP64 (top), FLAG tagged MCP-mCherry, FLAG tagged MCP-MSN, FLAG tagged MCP-p65-HSF1 (middle), and β-Tubulin (loading control; bottom) are shown as detected by Western blotting in HEK293T cells 72 hours post-transfection. **d and e**. Relative expression levels of endogenous human genes 72 hours after Control, DREAM, or SAM systems were targeted to their respective promoters using pools of 4 or 3 gRNAs (*HBG1* and *CD34*, respectively; **panel d**), or using single gRNAs (*ACE2* and *HGF*, respectively; **panel e**) as measured by QPCR. **f.** Transcriptome wide RNA-seq data generated 72 hours after the DREAM (top) or SAM (bottom) systems were targeted to the *HBG1/HBG2* promoter using 4 pooled gRNAs. mRNAs identified as significantly differentially expressed (fold change >2 or <-2 and FDR <0.05) are shown as red dots in both MA plots. In the top MA plot (CRISPR-DREAM), mRNAs corresponding to *HBG1/HBG2* (target genes) are highlighted in light blue. mRNAs encoding components of the MSN tripartite fusion protein (*MRTF-A/STAT1/NRF2*; red), were also significantly differentially expressed (fold change >2 and FDR <0.05). In the bottom MA plot (SAM system), mRNAs corresponding to *HBG1/HBG2* (target genes) are highlighted in light gray. *HSF1* mRNA (a component of the p65-HSF1 bipartite fusion protein; red), was also significantly differentially expressed (fold change >2 and FDR <0.05). **g.** 6 endogenous genes were activated by DREAM or SAM using a pool of gRNAs (1 gRNA/gene) in HEK293T cells. **h and i.** *OCT4* (**panel h**) or *HBG1* (**panel i**) gene activation by DREAM or SAM systems when corresponding promoters were targeted by 4 gRNAs per promoter in hTERT-MSC or PMBC cells, respectively. All QPCR samples were processed 72 hours post-transfection and are the result of at least 3 biological replicates. See source data for more information. Error bars; SEM. *; *P* < 0.05, **; *P* < 0.01, ***; *P* < 0.001. *ns*; not significant.

To test the transcriptome-wide specificity of CRISPR-DREAM, we used 4 gRNAs to target the DREAM or the SAM system to the *HBG1*/*HBG2* locus in HEK293T cells and then performed RNA-seq (**Fig. 1f**). *HBG1*/*HBG2* gene activation was specific and potent for both the CRISPR-DREAM and SAM systems relative to dCas9 + MCP-mCherry control treated cells. However, DREAM activated substantially more H*BG1*/*HBG2* transcription than the SAM system or dCas9-VPR (**Fig. 1f and Supplementary Fig. 13**). DREAM maintained superior efficacy relative to the SAM system and dCas9-VPR at *HBG1* (and *SBNO2*) across time points (up to at least 12 days) and cell passages (**Supplementary Fig. 14**). We also found that the DREAM system was significantly (*P* < 0.05) more potent than the SAM system at all targeted genes when each system was combined with a pool of six gRNAs, each targeting a different gene (**Fig. 1g**). Additionally, we evaluated the efficacy of the DREAM system across a battery of different human cell types, including a diverse panel of cancer cell lines (**Fig. 1h and Supplementary Fig. 15**) as well as primary and/or karyotypically normal human cells (**Fig. 1i and Supplementary Fig. 16**). Finally, we tested the transactivation potency of the DREAM system in mammalian cell types widely used for disease modeling/biocompatibility applications and therapeutic production pipelines (NIH3T3 and CHO-K1 cells, respectively; **Supplementary Fig. 17**). Across all experiments the DREAM system displayed highly potent transactivation. Overall, our data demonstrate that CRISPR-DREAM is robust, broadly potent, specific, and functionally compatible with diverse human and mammalian cell types.

### CRISPR-DREAM efficiently catalyzes RNA synthesis from noncoding genomic regulatory elements

Since CRISPR-DREAM efficiently and robustly activated mRNAs when targeted to promoter regions, we next tested whether the DREAM system could also activate transcription from distal human regulatory elements (i.e., enhancers) and other non-coding transcripts (i.e., enhancer RNAs; eRNAs, long noncoding RNAs; lncRNAs, and microRNAs; miRNAs). We first targeted the DREAM or SAM systems to the *OCT4* distal enhancer (DE)^40^ and found that the DREAM system significantly (*P* < 0.05) upregulated *OCT4* expression relative to the SAM system when targeted to the DE (**Fig. 2a**). Similar results were observed when targeting the DREAM system to the DRR enhancer^41^ upstream of the *MYOD* gene (**Supplementary Fig. 18a**). We also targeted the DREAM system to the human HS2 enhancer^42, 43^ and observed that the DREAM system induced expression from the downstream *HBE*, *HBG*, and *HBD* genes (**Fig. 2b**). We further observed transactivation of the *SOCS1* gene when the DREAM system was targeted to either of two different intragenic *SOCS1* enhancers; one located ∼15kb, and the other ∼50kb downstream of the *SOCS1* TSS (**Fig. 2c**). Together these data demonstrate that CRISPR-DREAM can stimulate human gene expression when targeted to different classes of enhancers (those regulating a single-gene, multiple genes, or intragenic enhancers) embedded within native chromatin.

**Fig. 2.**
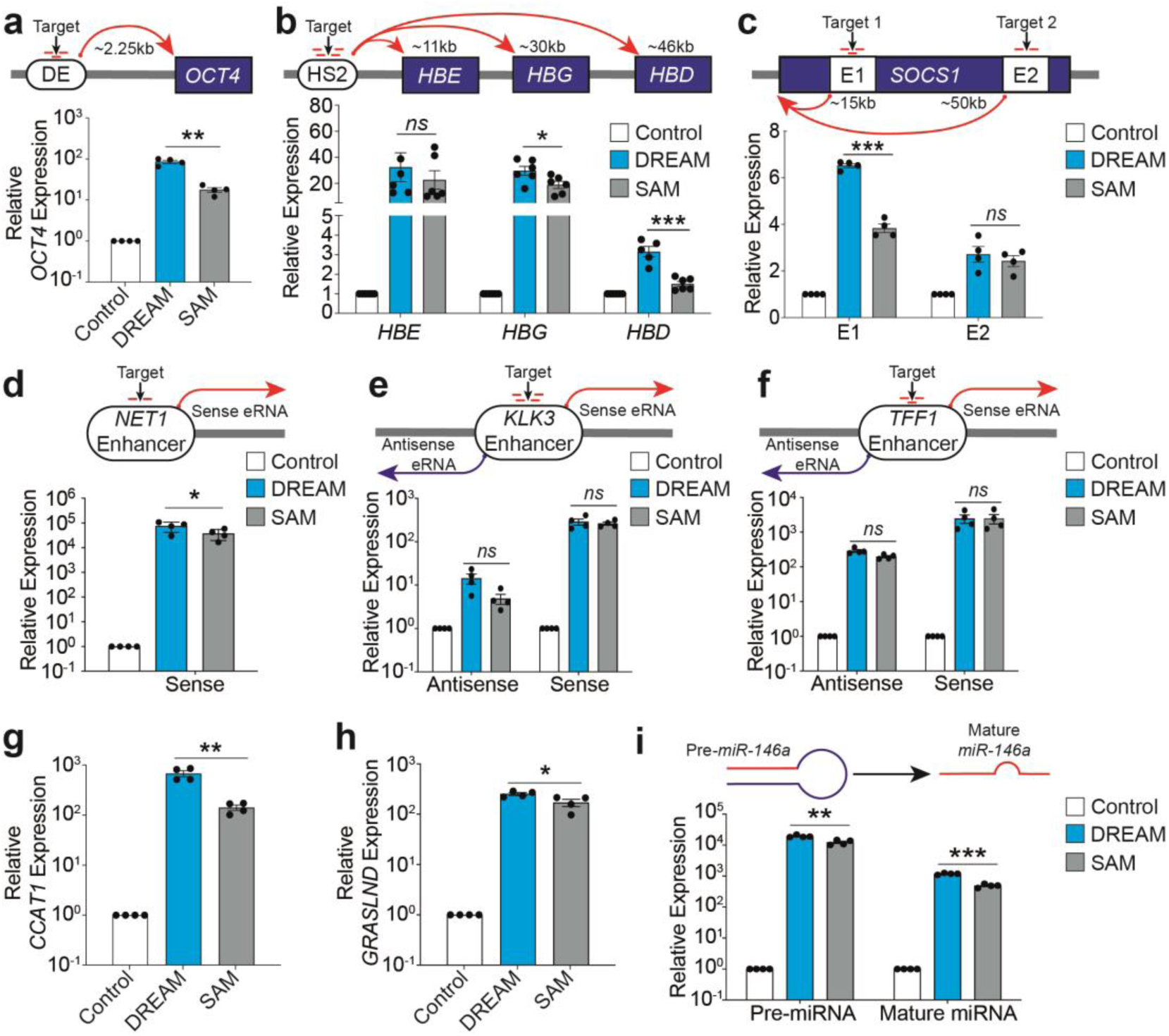
CRISPR-DREAM efficiently activates transcription from diverse human regulatory elements. **a-c.** CRISPR-DREAM and the SAM system activated downstream mRNA expression from *OCT4* (**panel a**), *HBE*, *HBG,* and *HBD* (**panel b**)*, and SOCS1* (**panel c**), when targeted to the *OCT4* distal enhancer (DE), HS2 enhancer, or one of two intragenic *SOCS1* enhancers, using pools of 3 (*OCT4* DE), 4 (*HS2*), 3 (*SOCS1* +15kb), or 2 (*SOCS1* + 50kb) gRNAs respectively. **d.** CRISPR-DREAM and the SAM system activated sense eRNA expression when targeted to the *NET1* enhancer using 2 gRNAs. **e and f.** CRISPR-DREAM and the SAM system bidirectionally activated eRNA expression when targeted to the *KLK3* (**panel e**) or *TFF1* (**panel f**) enhancers using pools of 4 or 3 gRNAs, respectively. **g and h.** CRISPR-DREAM and the SAM system activated the expression of long noncoding RNA when targeted to the *CCAT1* (**panel g**) or *GRASLND* (**panel h**) promoters using pools of 4 gRNAs, respectively. **i.** CRISPR-DREAM and the SAM system activated the expression of pre and mature *miR-146a* when targeted to the *miR-146a* promoter using a pool of 4 gRNAs. All samples were processed for QPCR 72 hours post-transfection. Data are the result of at least 4 biological replicates. See source data for more information. Error bars; SEM. *; *P* < 0.05, **; *P* < 0.01, ***; *P* < 0.001. *ns*; not significant.

We next tested whether CRISPR-DREAM could activate eRNAs when targeted to endogenous human enhancers. When targeted to the *NET1* enhancer, the DREAM system activated eRNA transcription (**Fig. 2d**), consistent with other reports^44^. Moreover, when the DREAM system was targeted to the bidirectionally transcribed *KLK3* and *TFF1* enhancers, we observed substantial upregulation of eRNAs in both the sense and antisense directions (**Fig. 2e and f**). Similar results were obtained when targeting the human *FKBP5* and *GREB1* enhancers (**Supplementary Fig. 18b and c**). CRISPR-DREAM also stimulated the production of endogenous lncRNAs when targeted to the *CCAT1, GRASLND*, *HOTAIR*, or *MALAT1* loci (**Fig. 2g and h, Supplementary Fig. 18d and e**). Finally, we found that the DREAM system activated *miRNA-146a* expression when targeted to the *miRNA-146a* promoter (**Fig. 2i**). Taken together, these data show that CRISPR-DREAM can robustly transactivate regulatory regions spanning diverse classes of the human transcriptome.

### Smaller, orthogonal CRISPR-DREAM platforms enable expanded genomic targeting beyond NGG PAM sites

To enhance the versatility of CRISPR-DREAM beyond SpdCas9 and to expand targeting to non-NGG PAM sites, we selected the two smallest naturally occurring orthogonal Cas9 proteins; SadCas9 (1,096aa) and CjdCas9 (1,027aa) for further analyses (**Fig. 3a, 3d**). We used SadCas9-specific gRNAs harboring MS2 loops^45^ to compare the potency between the SadCas9-DREAM and SAM systems in HEK293T cells. SadCas9-DREAM was significantly (*P* < 0.05) more potent than SadCas9-SAM when targeted to either the *HBG1* or *TTN* promoters (**Fig. 3b**). We also found that SadCas9-DREAM outperformed or was comparable to SadCas9-VPR when targeted to these loci (**Fig. 3c**) and that SadCas9-DREAM displayed high levels of transactivation in a second human cell line (**Supplementary Fig. 19a and b**).

**Fig. 3.**
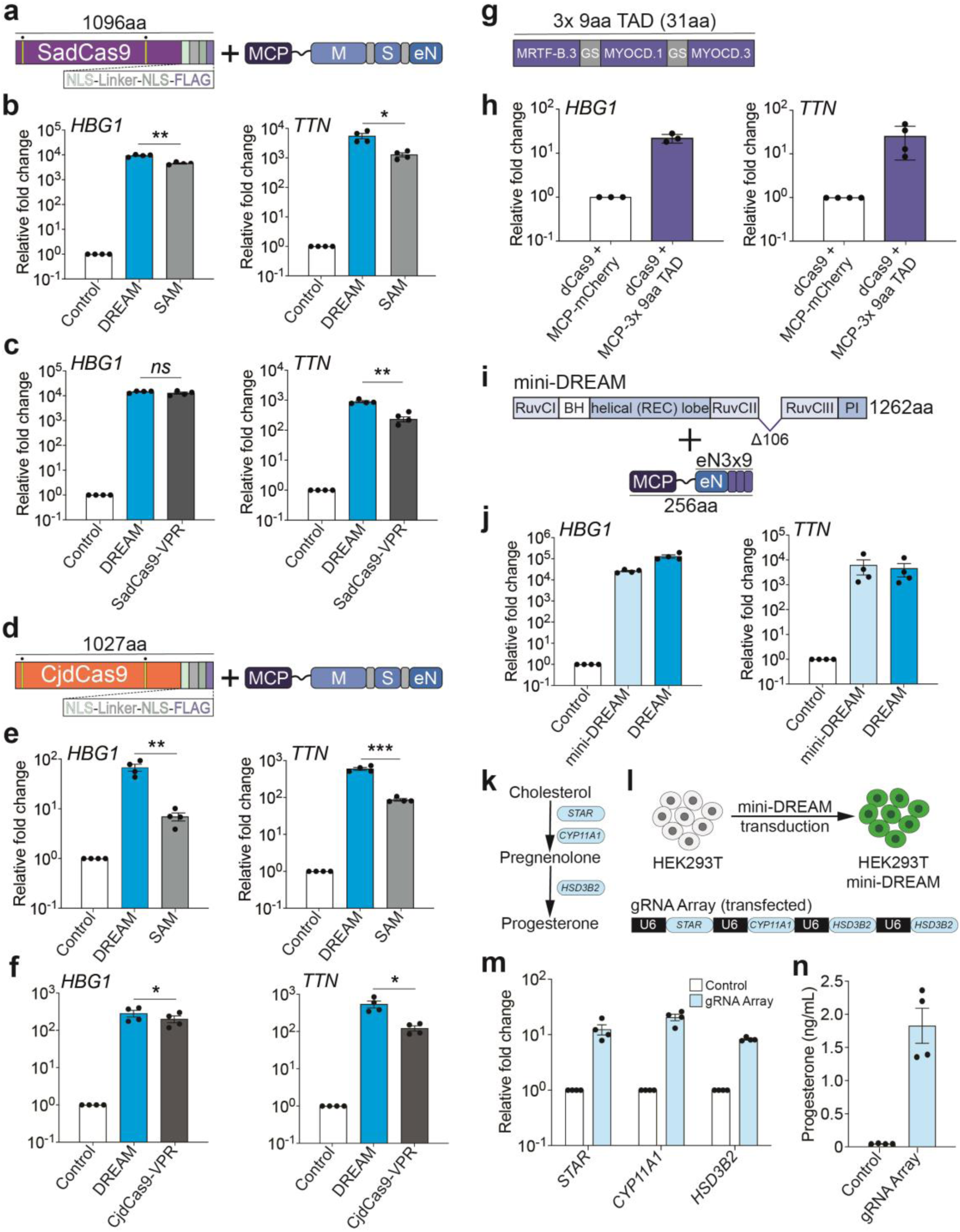
CRISPR-DREAM is portable to orthogonal dCas9 proteins and amenable to miniaturization. **a.** The SadCas9-DREAM system is schematically depicted, and nuclease-inactivating mutations (D10A and N580A) are indicated by yellow bars with dots above. **b.** *HBG1* (left) or *TTN* (right) gene activation using the SadCas9-DREAM or SadCas9-SAM systems, when targeted to each corresponding promoter using pools of 4 gRNAs, respectively. **c.** *HBG1* (left) or *TTN* (right) gene activation using the SadCas9-DREAM or SadCas9-VPR systems, when targeted to each corresponding promoter using pools of 4 MS2-modifed (SadCas9-DREAM) or standard gRNAs (SadCas9-VPR), respectively. **d.** The CjdCas9-DREAM system is schematically depicted, and nuclease-inactivating mutations (D8A and H559A) are indicated by yellow bars with dots above. **e.** *HBG1* (left) or *TTN* (right) gene activation using the CjdCas9-DREAM or CjdCas9-SAM systems, when targeted to each corresponding promoter using pools of 3 MS2-modified gRNAs, respectively. **f.** *HBG1* (left) or *TTN* (right) gene activation using the CjdCas9-DREAM or CjdCas9-VPR systems, when targeted to each corresponding promoter using pools of 3 MS2-modifed (SadCas9-DREAM) or standard gRNAs (CjdCas9-VPR), respectively. **g.** A 3x 9aa TAD derived from MYOCD and MRTF-B TADs is schematically depicted, GS; glycine-serine linker. **h.** *HBG1* (left) or *TTN* (right) gene activation when the 3x 9aa TAD was fused to MCP and recruited to each corresponding promoter using dCas9 and a pool of 4 MS2-modified gRNAs, respectively. **i.** The mini-DREAM system is schematically depicted. MCP-eN3×9 is a fusion protein consisting of MCP, eNRF2, and the 3x 9aa TAD derived from MYOCD and MRTF-B TADs. **j.** *HBG1* (left) or *TTN* (right) gene activation when either the mini-DREAM or CRISPR-DREAM system was targeted to each corresponding promoter using a pool of 4 MS2-modified gRNAs, respectively. **k.** A simplified biosynthetic pathway for progesterone production via cholesterol is schematically depicted. **l.** The workflow to build progesterone-producing HEK293T cell factories using the mini-DREAM platform and corresponding gRNA array is shown (see Methods for more details). **m and n.** *STAR*, *CYP11A1* and *HSD3B2* gene activation and secreted progesterone levels, respectively, after mini-DREAM-transduced HEK293T cells were transfected with the indicated gRNA array (targeting STAR, *CYP11A1* and *HSD3B2*) or a non-targeting gRNA control plasmid. All samples were processed for QPCR or ELISA 72 hours post-transfection. Data are the result of at least 3 biological replicates. See source data for more information. Error bars; SEM. *; *P* < 0.05, **; *P* < 0.01, ***; *P* < 0.001. *ns*; not significant.

CjdCas9-based transcriptional activation platforms have also recently been developed using viral TADs (e.g., miniCAFE)^46^; however, gRNA-based recruitment of transcriptional modulators using CjdCas9 has not been described. Therefore, we engineered the CjdCas9 gRNA scaffold to incorporate an MS2 loop within the tetraloop of the CjCas9 gRNA scaffold (**Supplementary Fig. 19c**). We used this MS2-modified CjdCas9 gRNA to generate CjdCas9-DREAM and compared the potency between CjdCas9-DREAM, CjdCas9-SAM, and CjdCas9-VPR at the *HBG1* or *TTN* promoters (**Fig. 3e and f**) in HEK293T cells. At all targeted sites, CjdCas9-DREAM outperformed or was comparable to the CjdCas9-SAM or CjdCas9-VPR systems. We also observed high levels of transactivation using CjdCas9-DREAM in a different human cell line (**Supplementary Fig. 19d and e**). These data demonstrate that DREAM is not only compatible with other orthogonal dCas9 targeting systems, but that it displays superior performance in terms of CRISPRa activity at most tested promoters.

### Generation and validation of a compact mini-DREAM system

We next sought to reduce the sizes of the CRISPR-DREAM components. We first investigated whether individual TADs could be minimized while still retaining the transactivation potency when recruited by dCas9. We focused on individual TADs from MTFs that displayed transactivation potential (i.e., MRTF-A, MRTF-B, and MYOCD proteins, **Supplementary Figs. 1 and 2**). As mentioned above, 9aa TADs have been shown to synthetically activate transcription previously using GAL4 systems^38, 47^. Therefore, we used predictive software^38^ to identify 9aa TADs in MRTF-A, MRTF-B, and MYOCD proteins, and recruited these TADs to human loci using dCas9 and MCP-MS2 fusions in single, bipartite, and tripartite formats (**Supplementary Note 1; Supplementary Fig. 20**). Interestingly, we observed that while single 9aa TADs did not activate endogenous gene expression, tripartite combinations of 9aa TADs were able to robustly activate endogenous genes, albeit to varying degrees (**Supplementary Fig. 20f**). We selected one tripartite 9aa combination (3x 9aa TAD; MRTF-B.3 + MYOCD.1 + MYOCD.3) for further analysis (**Fig. 3g**). This 3x 9aa TAD activated *HBG1*, *TTN*, and *CD34* gene expression when recruited to corresponding promoters using dCas9 (**Fig 3h; Supplementary Fig. 20g**). We also found that this 3x 9aa TAD combination could activate gene expression via a single gRNA, and moreover could transactivate other endogenous regulatory loci (**Supplementary Fig. 20h – j**). These results suggest that combinations of 9aa TADs can be used as minimal functional units to transactivate endogenous human loci when recruited via dCas9.

We next combined the 3x 9aa TAD with the engineered NRF2 TAD (eNRF2) in four different combinations to generate a small, yet potent transactivation module called eN3×9 (**Supplementary Fig. 21**). Notably, minimized Cas9 proteins that retain DNA binding activity have also been recently created^48, 49^. Therefore, we next evaluated the relative transactivation capabilities among a panel of minimized, HNH-deleted, dCas9 variants in tandem with MCP-MSN and found that an HNH-deleted variant without a linker between two RuvC domains was optimal, albeit with slight protein expression decreases (**Supplementary Fig. 22a and b**). We further validated this linker-less, HNH-deleted CRISPR-DREAM variant at multiple human promoters and other regulatory elements (**Supplementary Fig. 22c – h**) and then combined this minimized dCas9 with MCP-eN3×9 to generate the mini-DREAM system (**Fig. 3i**). The mini-DREAM system transactivated *HBG1*, *TTN*, and *IL1RN* gene expression when recruited to corresponding promoters (**Fig 3j; Supplementary Fig. 23a**). We also found that the mini-DREAM system could activate endogenous promoters via a single gRNA (**Supplementary Fig. 23b and c**) and could activate downstream gene expression when targeted to an upstream enhancer (**Supplementary Fig. 23d**).

To demonstrate the utility of the mini-DREAM platform, we used this system to create progesterone producing HEK293T cell factories. Specifically, we simultaneously targeted and activated three key genes in the progesterone production pathway (*STAR*, *CYP11A1*, and *HSD3B2*), which resulted in increased target gene expression and significant production of progesterone (**Fig. 3k – n**). We also evaluated whether the minimized components of the mini-DREAM system were functional when delivered within a single vector (**Supplementary Fig. 23e**) and found that this compact, single vector mini-DREAM system retained transactivation potential when targeted to human promoters using or a single gRNA (**Supplementary Fig. 23 f – i**). Notably, mini-DREAM and mini-DREAM compact also outperformed the miniCAFE platform at two different loci in HEK293T cells (**Supplementary Fig. 23j and k**). Overall, these data show that the components of the CRISPR DREAM system can be substantially minimized while retaining functionality.

### The MSN and NMS effector domains are robust across programmable DNA binding platforms

We next tested the potency of tripartite MSN and NMS effectors when fused to dCas9 in different architectures and observed that both effectors could activate gene expression when fused to the N- or C-terminus of dCas9 (**Supplementary Note 2; Supplementary Fig. 24**) or when recruited via the Sun-Tag^14^ architecture (**Supplementary Fig. 25**). Interestingly, in contrast to MCP-mediated recruitment (**Supplementary Fig. 8**), additional TADs were observed to improve performance in direct fusion architectures (**Supplementary Fig. 24a; Supplementary Note 2**). In the SunTag architecture, the NMS domain was superior to other benchmarked effector domains, such as VP64^14^, VPR^50^, and p65-HSF1^51^ (**Supplementary Fig. 25a – c**). To maximize the potential use of the MSN/NMS effector domains and explore their versatility, we next tested whether each was capable of gene activation when fused to TALE or ZF scaffolds (**Fig. 4a and b**). Both effectors strongly transactivated *IL1RN* using a single TALE fusion protein (**Supplementary Fig. 26**) or a pool of 4 TALE fusion proteins targeted to the *IL1RN* promoter (**Fig 4a**). Similarly, both effectors activated *ICAM1* expression using a single synthetic ZF fusion protein targeted to the *ICAM1* promoter (**Fig. 4b**). These data demonstrate that the MSN and NMS effectors are compatible with diverse programmable DNA binding scaffolds beyond Type II CRISPR/Cas systems.

**Fig. 4.**
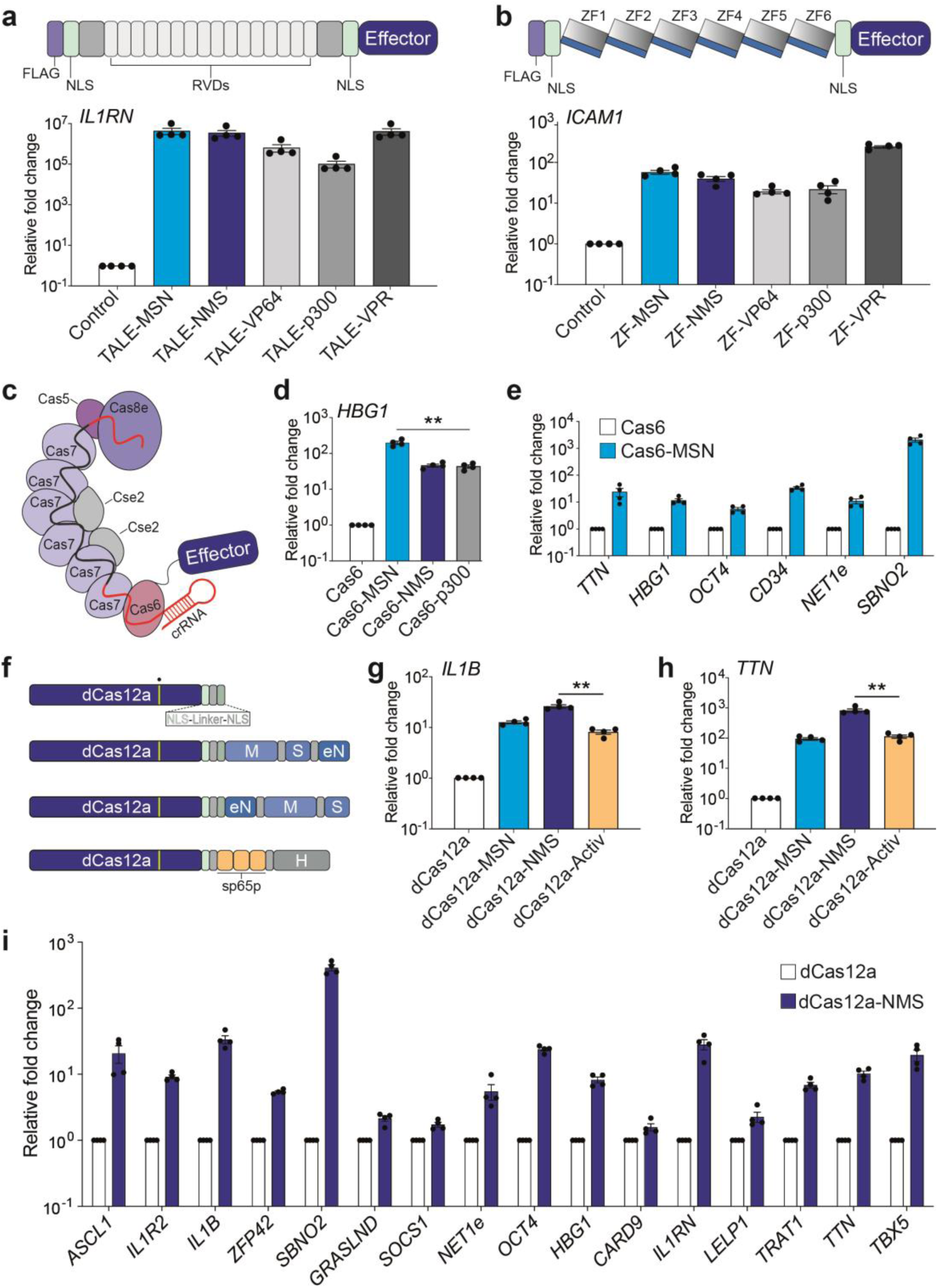
The MSN and NMS effector domains are portable to diverse DNA binding platforms and enable superior multiplexing when fused to dCas12a. **a**. Synthetic transcription activator-like effector (TALE) proteins harboring indicated effector domains were designed to target the human *IL1RN* promoter. Repeat variable di-residues, RVDs. Relative *IL1RN* expression (bottom) 72 hours after indicated TALE fusion protein encoding plasmids were transfected. **b**. Synthetic zinc finger (ZF) proteins harboring indicated effector domains were designed to target the human *ICAM1* promoter. Relative *ICAM1* expression (bottom) 72 hours after indicated ZF fusion protein encoding plasmids were transfected. **c**. The Type I CRISPR system derived from *E. Coli* K-12 (Eco-cascade) is schematically depicted along with an effector fused to the Cas6 protein subunit. **d**. *HBG1* gene activation when either the MSN, NMS, or p300 effector domains were fused to Cas6 and the respective engineered Eco-Cascade complexes were targeted to the *HBG1* promoter using a single crRNA. **e**. Multiplexed activation of 6 endogenous genes 72hours after co-transfection of Eco-cascade complexes when MSN was fused to Cas6 and targeted using a single crRNA array expression plasmid (1 crRNA/promoter). **f.** The dCas12a protein and indicated fusions are schematically depicted along with the G993A DNase-inactivating mutation indicated by a yellow bar with a dot above. **g and h**. *IL1B* (**panel g**) or *TTN* (**panel h**) gene activation using the indicated dCas12a fusion proteins when targeted to each corresponding promoter using a pool of 2 crRNAs (for *IL1B*) or a single array encoding 3 crRNAs (*TTN*), respectively. **i**. Multiplexed activation of 16 indicated endogenous genes 72hours after co-transfection of dCas12a-NMS and a single crRNA array expression plasmid encoding 20 crRNAs. All samples were processed for QPCR 72 hours post-transfection in HEK293T cells. See source data for more information. Data are the result of at least 4 biological replicates. Error bars; SEM. **; *P* < 0.01.

Transcriptional activators have recently been shown to modulate the expression of endogenous human loci when recruited by Type I CRISPR systems^52^. Therefore, to evaluate whether MSN and/or NMS were functional beyond Type II CRISPR systems, we fused each to the Cas6 component of the *E. coli* Type I CRISPR Cascade (Eco-Cascade) system (**Fig. 4c**). Our data showed that Cas6-MSN (or NMS) performed comparably to the Cas6-p300 system when targeted to a spectrum of human promoters (**Fig. 4d; Supplementary Fig. 27a – d**). We also observed that the Cas6-MSN (or NMS) systems could activate eRNAs from when targeted to the endogenous *NET1* enhancer (**Supplementary Fig. 27e**). One advantage of CRISPR Cascade is that the system can process its own crRNA arrays, which can enable multiplexed targeting to the human genome. Previous reports have leveraged this capability to simultaneously activate two human genes^52^. We found that when Cas6 was fused to MSN, the CRISPR Cascade system could simultaneously activate up to six human genes when corresponding crRNAs were co-delivered in an arrayed format (**Fig. 4e; Supplementary Fig. 27f**). We also found that these transactivation capabilities were extensible to another Type I CRISPR system; *Pae*-Cascade^53^ (**Supplementary Fig. 27g – i**). In sum, these data show that the MSN and NMS effectors are robust and directly compatible with programmable DNA binding platforms beyond Type II CRISPR systems without any additional engineering.

### The NMS effector enables superior multiplexed gene activation when fused to dCas12a

The CRISPR/Cas12a system has attracted significant attention because the platform is smaller than SpCas9, and because Cas12a can process its own crRNA arrays in human cells^54^. This feature has been leveraged for both multiplexed genome editing and multiplexed transcriptional control^18^. Therefore, we next investigated the potency of the tripartite MSN and NMS effectors when they were directly fused to dCas12a (**Fig. 4f**). We selected the AsdCas12a variant for this analysis because AsdCas12a (hereafter dCas12a) has been shown to activate human genes when fused to transcriptional effectors^18^. Our results demonstrated that both dCas12a-MSN and dCas12a-NMS were able to induce gene expression when targeted to different human promoters using pooled or single crRNAs (**Fig. 4g and h, Supplementary Fig. 28a – e**). dCas12a-NMS was generally superior to dCas12a-MSN and to the previously described dCas12a-Activ system^18^ at the loci tested here. These data demonstrate that the NMS and MSN effectors domains are potent transactivation modules when combined with the dCas12a targeting system in human cells.

We next tested the extent to which dCas12a-MSN/NMS could be used in conjunction with crRNA arrays for multiplexed endogenous gene activation. We cloned 8 previously described crRNAs^18^ (targeting the *ASCL1*, *IL1R2*, *IL1B* or *ZFP42* promoters) into a single plasmid in an array format and then transfected this vector into HEK293T cells with either dCas12a control, dCas12a-MSN, dCas12a-NMS, or the dCas12a-Activ system. Again, our data demonstrated that dCas12a-NMS was superior or comparable to dCas12a-Activ, even in multiplex settings (**Supplementary Fig. 28f**). Finally, to evaluate if dCas12-NMS could simultaneously activate multiple genes on a larger scale, we cloned 20 full-length (20bp) crRNAs targeting 16 different loci into a single array (**Supplementary Fig. 28g**). This array was designed to enable simultaneous targeting of several classes of human regulatory elements; including 13 different promoters, 2 different enhancers (one intrageneric; *SOCS1,* and one driving eRNA output; *NET1*), and one lncRNA (*GRASLND*). When this crRNA array was transfected into HEK293T cells along with dCas12a-NMS, RNA synthesis was robustly stimulated from all 16 loci (**Fig. 4i**). To our knowledge this is the most loci that have been targeted simultaneously using CRISPR systems, demonstrating the versatility and utility of the engineered NMS effector in combination with dCas12a.

### dCas9-NMS permits efficient reprogramming of human fibroblasts *in vitro*

CRISPRa systems using repeated portions of the alpha herpesvirus VP16 TAD (dCas9-VP192) have been used to efficiently reprogram human foreskin fibroblasts (HFFs) into induced pluripotent stem cells (iPSCs)^17^. To evaluate the functional capabilities of our engineered human transactivation modules, we fused the NMS domain directly to the C-terminus of dCas9 (dCas9-NMS) and tested its ability to reprogram HFFs. We used a direct dCas9 fusion architecture so that we could leverage gRNAs previously optimized for this reprogramming strategy and to better compare dCas9-NMS with the corresponding state of the art (dCas9-VP192)^17^. We used the NMS effector as opposed to MSN, as NMS displayed more potency than MSN when directly fused to dCas9 (**Supplementary Fig. 24a**). We targeted dCas9-NMS (or dCas9-VP192) to endogenous loci using the 15 gRNAs previously optimized to reprogram HFFs to pluripotency with the dCas9-VP192 system. Using this approach, we observed morphological changes beginning by 8 days post-nucleofection (**Fig. 5a**) and efficient reprogramming by 16 days post-nucleofection, although to a lesser extent than when using dCas9-VP192 (**Supplementary Fig. 29a**).

**Fig. 5.**
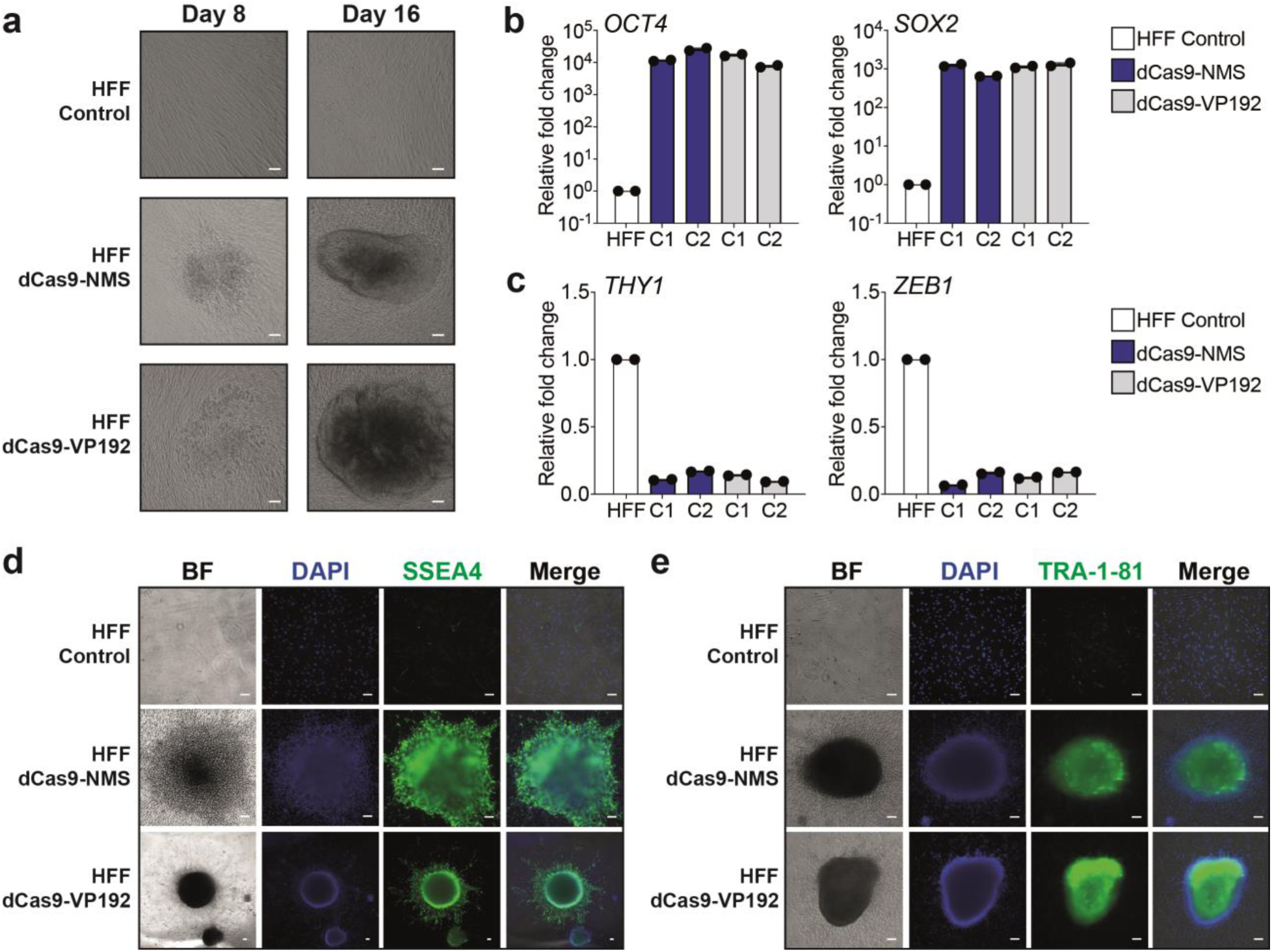
dCas9-NMS permits efficient *in vitro* reprogramming of human fibroblasts. **a**. Primary human foreskin fibroblasts (HFFs) were nucleofected with plasmids encoding 15 multiplexed gRNAs targeting the *OCT4*, *SOX2*, *KLF4*, c*-MYC*, and *LIN28A* promoter and EEA motifs (as in previous reports^17^), and either dCas9-NMS (middle row) or dCas9-VP192 (bottom row). HFF morphology was analyzed 8 and 16 days later (white scale bars, 100μm). **b.** Relative expression of pluripotency-associated genes *OCT4* (left) and *SOX2* (right) in representative iPSC colonies (C1 or C2) approximately 40 days after nucleofection of either dCas9-NMS (blue) or dCas9-VP192 (gray) and multiplexed gRNAs compared to untreated HFF controls. **c.** Relative expression of mesenchymal-associated genes *THY1* (left) and *ZEB1* (right) in representative iPSC colonies (C1 or C2) approximately 40 days after nucleofection of either dCas9-NMS (blue) or dCas9-VP192 (gray) and multiplexed gRNAs compared to untreated HFF controls. **d and e.** Immunofluorescence microscopy of HFFs approximately 40 days after nucleofection of either dCas9-NMS or dCas9-VP192 and multiplexed gRNAs compared to untreated HFF controls (white scale bars, 100μm). Cells were stained for the expression of pluripotency-associated cell surface markers SSEA4 (**panel d**, green) or TRA-1-81 (**panel e**, green). All cells were counterstained with DAPI for nuclear visualization.

We picked and expanded iPSC colonies and then measured the expression of pluripotency and mesenchymal genes ∼40 days post-nucleofection. We found that genes typically associated with pluripotency (*OCT4, SOX2, NANOG, LIN28A, REX1, CDH1,* and *FGF4*)^55, 56^ were highly expressed in colonies derived from HFFs nucleofected with the gRNA cocktail and dCas9-NMS or dCas-VP192 (**Fig. 5b; Supplementary Fig. 29b – f**). Conversely, we observed that genes typically associated with fibroblast/mesenchymal cell identity (*THY1, ZEB1, ZEB2, TWIST,* and *SNAIL2*)^55, 56^ were poorly expressed in colonies derived from HFFs nucleofected with the gRNA cocktail and dCas9-NMS or dCas-VP192 (**Fig. 5c; Supplementary Fig. 29g – i)**^57^ and found that all were highly expressed in iPSC colonies derived from HFFs nucleofected with the gRNA cocktail and either dCas9-NMS or dCas-VP192 (**Fig. 5d and e; Supplementary Fig. 29j**). These data show that engineered transactivation modules sourced from human MTFs can be used to efficiently reprogram complex cell phenotypes, including cell lineage.

### The MSN, NMS, and eN3×9 transactivation modules are well tolerated and effective in clinically useful primary human cell types

The recent development of CRISPRa tools has enabled new therapeutic opportunities^6, 58^. However, it has been shown that in some cases, CRISPRa tools harboring viral TADs can be poorly tolerated, and even toxic^12, 21–23^. This prompted us to test the relative expression and efficacy of the human MTF derived multipartite TADs MSN, NMS, and eN3×9 tools in comparison to the viral multipartite TAD VPR in therapeutically relevant human primary cells. We selected primary human umbilical cord MSCs and primary T cells for analysis. Lentiviral transduction was selected to ensure high levels of payload delivery. Interestingly, we observed that lentiviral titers were influenced by fused TAD, with MCP fused to eN3×9 consistently generating the highest titers (**Supplementary Fig. 30**). We next transduced MSCs using an MOI of ∼10.0 for all conditions and observed variable expression levels among MCP fusions proteins at 72 hours post-transduction using both microscopy and flow cytometry (**Fig. 6a**) despite using equal amounts of lentivirus. For instance, although MCP-eN3×9 and MCP-NMS displayed high levels of expression via microscopy, MCP-VPR and MCP-MSN were relatively poorly expressed. Similarly, we tested the expression levels of these MCP fusions in primary T cells using lentiviral transduction at a fixed MOI of ∼5.0 across conditions and observed that MCP-eN3×9 displayed the highest expression levels 72 hours post-transduction, while MCP-VPR showed the lowest expression (**Fig. 6b**).

**Fig. 6.**
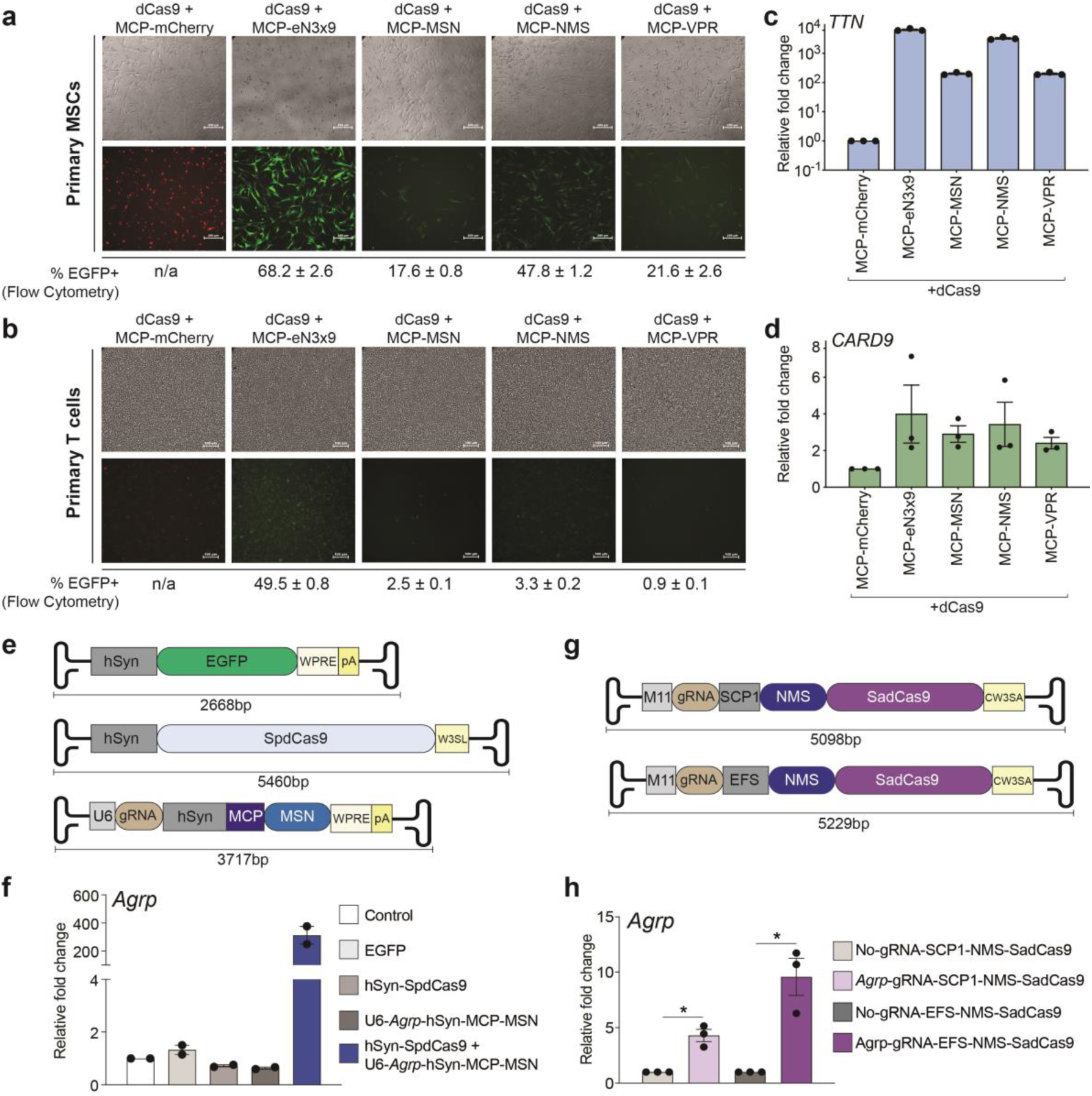
CRISPR-DREAM components are well tolerated in primary cells and compatible with viral delivery methods. **a and b.** Immunofluorescence microscopy showing mCherry/EGFP expression levels in MSCs (**panel a**) and human T cells (**panel b**) 72 hours after co-transduction of dCas9 in combination with either MCP-mCherry (control), MCP-eN3×9-T2A-EGFP, MCP-MSN-T2A-EGFP, MCP-NMS-T2A-EGFP, or MCP-VPR-T2A-EGFP respectively (white scale bars, 250μm for MSCs, 100μm for T cells). MCP-fusion vectors also contain a U6 driven gRNA expression cassette and either a *TTN* (MSCs) or *CARD9* (T cells). **c and d.** Relative expression of *TTN* (**panel c**) or *CARD9* (**panel d**) in MSCs and T cells, respectively, 3 days after lentiviral co-transduction using indicated components. **e**. AAV constructs used for dual-delivery of CRISPR-DREAM components are schematically depicted. The EFGP control vector is shown (top) along with the hSyn promoter driven SpdCas9 vector (middle), which consists of a modified WPRE/polyA sequence (W3SL). The U6 promoter driven gRNA expressing vector (bottom) is also shown and also encodes MCP fused to MSN, which is driven by the hSyn promoter. **f**. *Agrp* gene activation in mouse primary cortical neurons using the dual AAV8 transduced CRISPR-DREAM system described (in **panel e**) 5 days post-transduction. **g**. All-in-one (AIO) SadCas9-based AAV vectors are schematically depicted. AIO vectors consist of M11 promoter driven gRNA cassettes and either SCP1 (top) or EFS (bottom) promoter driven NMS-SadCas9. A modified WPRE/polyA sequence (CW3SA) was used in the AIO vectors. **h**. *Agrp* gene activation in mouse primary cortical neurons transduced with AIO AAV vectors (in **panel h**) 5 days post-transduction. Data are the result of at least 2 biological replicates. See source data for more information. Error bars; SEM. *; *P* < 0.05.

We next assessed the gene activation capabilities of these MCP-TAD fusions in primary MSCs and T cells. In MSCs, eN3×9 outperformed all other effectors, and VPR showed the lowest potency when targeted to the *TTN* promoter (**Fig. 6c**). In primary T cells each TAD activated *CARD9* expression to relatively similar and modest levels when targeted to the *CARD9* promoter (**Fig. 6d**). However, in primary T cells we observed that the human MTF derived multipartite TADs resulted in dramatically better T cell viability than the viral multipartite TAD VPR (**Supplementary Fig. 31**). Interestingly, these effects were less obvious in transformed cell lines (**Supplementary Fig. 32**). Collectively these data demonstrate that the human MTF derived multipartite MSN, NMS, and eN3×9 TADs are as or more potent than the VPR TAD, while also maintaining similar or superior expression levels in therapeutically relevant human primary cells. Notably, MSN, NMS, and eN3×9 are also much smaller than the VPR TAD, and in the case of primary T cells, are also much less cytotoxic.

### Dual and all-in-one AAV mediated delivery of CRISPR-DREAM and SadCas9-NMS systems efficiently activates gene expression in primary neurons

AAV mediated delivery has emerged as a powerful method to deliver therapeutic payloads *in vitro*^59^ and *in vivo*^60^. However, due to strict payload limitations, the delivery of CRISPRa tools using AAV has been limited to dual AAV systems and/or the use of viral TADs^61, 62^. To assess the transcriptional activation potential of the compact CRISPR-DREAM components in combination with AAV mediated delivery, we targeted the murine *Agrp* gene, which modulates food intake behavior and obesity^63, 64^, as a proof of concept. We first tested 15 individual gRNAs targeting a ∼1kb window upstream of the *Agrp* promoter in Neuro-2a cells to identify a top performing gRNA (**Supplementary Fig. 33a and b**). Based on these results, we constructed a dual AAV delivery system, wherein one AAV expressed dCas9, and the other AAV expressed the top performing *Agrp*-targeting gRNA along with MCP-MSN (**Fig. 6e**). Both recombinant AAVs (and an EGFP control AAV) used the AAV8 serotype capsid to ensure efficient neuronal transduction^65^ (**Supplementary Fig. 33e**). In dual AAV-transduced (dCas9 and gRNA/MCP-MSN, respectively) primary murine neurons, we observed high levels of *Agrp* activation **(Fig. 6f).**

Encouraged by this result using a dual AAV strategy, we next designed two different all-in-one (AIO) AAV approaches (**Fig. 6g**). These designs leveraged the M11 promoter to express a gRNA, and either the SCP1 or EFS promoter to drive the expression of NMS fused to the N-terminus of SadCas9. NMS was prioritized over MSN as it showed higher potency when fused to N-terminus of dCas9 (**Supplementary Fig. 24**). To further reduce packaging size, we also selected compact engineered WPRE and PolyA^66^ tail elements in these construct designs. After selecting a top performing *Agrp*-targeting SadCas9 gRNA in Neuro-2A cells **(Supplementary Fig. 33g and h)**, we made recombinant AAVs (using serotype AAV8) and delivered these AIO AAVs to primary murine neurons. In both cases, we observed significant (*P* value < 0.05) transcriptional upregulation of *Agrp*, with the EFS promoter harboring vector displaying superiority to the SCP promoter harboring vector (**Figure 6h**). These data demonstrate that the compact components of the CRISPR-DREAM retain high transactivation potency when delivered into primary cells using either dual or AIO AAV modalities.

## Discussion

Here we harnessed the programmability and versatility of different dCas9-based recruitment architectures (direct fusion, gRNA-aptamer, and SunTag-based) to optimize the transcriptional output of TADs derived from natural human TFs. We leveraged these insights to build superior and widely applicable transactivation modules that are portable across all modern synthetic DNA binding platforms, and that can activate the expression of diverse classes endogenous RNAs. We selected mechanosensitive TFs (MTFs) for biomolecular building blocks because they naturally display rapid and potent gene activation at target loci, can interact with diverse transcriptional co-factors across different human cell types, and because their corresponding TADs are relatively small^67–69^. We not only identified and validated the transactivation potential of TADs sourced from individual MTFs, but we also established the optimal TAD sequence compositions and combinations for use across different synthetic DNA binding platforms, including Type I, II and V CRISPR systems, TALE proteins, and ZF proteins.

Our study also revealed that for MTFs, tripartite fusions using TADs from MRTA-A (M), STAT1 (S), and NRF2 (N) in one of two different combinations (either MSN or NMS) consistently resulted in the most potent human gene activation across different DNA binding platforms. Interestingly, each of these components has been shown to interact with key transcriptional co-factors (**Supplementary Fig. 34a**). For example, individual TADs from MRTF-A, STAT1, NRF2 can directly interact with endogenous p300^32, 70^. Moreover, the Neh4 and Neh5 TADs from NRF2 can also cooperatively recruit endogenous CBP for transcriptional activity^30, 71^. MRTF-A and NRF2 can also interact with other histone modifiers and chromatin remodelers. For instance, MRTF-A can engage JMJD1A, SET1, and BRG1^72–74^, and NRF2 can also engage BRG1, as well as CDH6^75, 76^. Further, STAT1 and NRF2 can interact with components of the mediator complex^77, 78^. Therefore, we suspect that the potency of the engineered MSN and NMS effector proteins is likely related to their robust capacity to recruit powerful and ubiquitous endogenous transcriptional modulators, which is likely positively impacted by their direct tripartite fusion. In support of this hypothesis, we observed that the CRISPR/dCas9-recruited enhanced activation module (DREAM) system, (which harbored the MSN tripartite domain) significantly catalyzed both increased H3K4me3 and H3K27ac at targeted human promoters (**Supplementary Fig. 34b and c**).

Additionally, our study demonstrated that the superior transactivation capabilities of the CRISPR-DREAM system, again consisting of dCas9 and a gRNA-aptamer recruited MCP-MSN fusion, are not reliant upon the direct fusion(s) of any other proteins (viral or otherwise) to dCas9, in contrast to the SAM system which relies upon dCas9-VP64^15^. We used this advantage to combine the MCP-MSN module with HNH domain deleted dCas9 variants^48, 49^, which exhibited similar potencies to full-size dCas9 variants. To further reduce the size of CRISPR-DREAM, we built a minimal transactivation module (eN3×9; 96aa) by evaluating the potency of a suite of 9aa TADs from MTFs and by next combining the most potent variants with the small eNRF2 TAD. We then combined the minimized eN3×9 transactivation module with an HNH domain deleted dCas9 variant in two-vector (mini-DREAM) and single-vector (mini-DREAM compact) delivery architectures, which retained potent and functionally applicable transactivation capabilities.

We also integrated the MSN and NMS effectors with the Type I CRISPR/Cascade and Type II dCas12a platforms to enable superior multiplexed endogenous activation of human genes. This multiplexing capability holds tremendous promise for reshaping endogenous cellular pathways and/or engineering complex transcriptional networks. dCas9-based transcription factors harboring viral TADs have also been used for directed differentiation and cellular reprogramming^9, 17, 79, 80^. Here, we used established protocols based on dCas9-VP192 to show that we could reprogram human fibroblasts into iPSCs using dCas9 directly fused to the NMS transcriptional effector with similar gene expression profiles, times to conversion, and morphological characteristics compared to iPSCs derived using dCas9 fused to VP192^17^. However, dCas9-NMS resulted in slightly fewer iPSC colonies than dCas9-VP192, which we attribute to the reprogramming framework tested here being optimized for use with dCas9-VP192.

Additionally, we demonstrated that the MSN and NMS effectors were compatible with dual and all-in-one (AIO) AAV vectors. Our AIO AAV vector design, which combines the short SCP1 promoter, the short M11 gRNA promoter and the compact CW3SA modified WPRE/poly A tail elements, holds tremendous potential for future delivery architectures. Similarly, the potency of AIO AAV vectors encoding NMS-SadCas9 empower researchers with a new streamlined modality to induce endogenous gene expression *in vivo* that could be used within animal models or clinical settings. Finally, we found that the NMS, MSN, and eN3×9 TADs were well-expressed and potent in therapeutically important human cells. Although the tripartite VPR TAD contains the potent VP64 and RTA viral elements, in our primary cell experiments VPR showed the lowest expression levels and gene activation potencies. In contrast, the hypercompact eN3×9 TAD was well expressed in both MSCs and T cells. In MSCs eN3×9 was also extremely potent, however in T cells, gene activation efficacy was modest for all activators tested. Nevertheless, MSN, NMS, and eN3×9 TADs were substantially less toxic compared to the VPR TAD in T cells.

While studies using CRISPR systems in combination with viral TADs have observed toxic effects at the cellular and organismal levels, it should be noted that our experiments do not conclusively demonstrate that viral TADs themselves are toxic. One remaining limitation that affects all synthetic gene activation platforms is that some loci may remain refractory to engineered transactivation, regardless of effector deployed. This constraint likely stems from high basal expression levels at targeted sites^12, 15^ (**Supplementary Fig. 35**) and/or other contextual factors that require further interrogation. Focused analyses at specific target sites, within specific cell types/organisms^81^, and over longer time courses will likely be informative for optimized therapeutic use cases.

The new TADs developed here repeatedly displayed high potency despite having small coding sizes. The potency of CRISPR-DREAM was particularly evident when measured on a per cell basis (**Supplementary Fig. 10**). In addition, since CRISPR-DREAM does not rely upon any effector fused to dCas9, it could synergize with future dCas9-encoding model organisms. Further, the CRISPR-DREAM system (and associated derivatives) are the most potent tools ever developed in the context of arrayed multiplexed synthetic gene activation (**Figs. 3m, 4e, 4i, and Supplementary Fig. 28f**).

In summary, we have used the rational redesign of natural human TADs to build synthetic transactivation modules that enable consistent and potent performance across programmable DNA binding platforms, mammalian cell types, and genomic regulatory loci embedded within human chromatin. Although we used MTFs as sources of TADs here, our work establishes a framework that could be used with practically any natural or engineered TF and/or chromatin modifier in future efforts. The potency, small size, versatility, capacity for multiplexing, and the lack of components from pathogenic human viruses associated with the newly engineered MSN, NMS, and eN3×9 TADs and CRISPR-DREAM systems developed here could be valuable tools for fundamental and biomedical applications requiring potent and predictable activation of endogenous eukaryotic transcription.

## Methods

### Cell culture

All experiments were performed within 10 passages of cell stock thaws. HEK293T (ATCC, CRL-11268), HeLa (ATCC, CCL-2), A549 (ATCC, CCL-185), SK-BR-3 (ATCC, HTB-30), U2OS (ATCC, HTB-96), HCT116 (ATCC, CRL-247), K562 (ATCC, CRL-243), CHO-K1 (ATCC, CCL-61), ARPE-19 (ATCC, CRL-2302), HFF (ATCC, CRL-2429), Jurkat-T (ATCC, TIB-152), hTERT-MSC (ATCC, SCRC-4000), and Neuro-2a (ATCC, CCL-131) cells were purchased from American Type Cell Culture (ATCC, USA) and cultured in ATCC-recommended media supplemented with 10% FBS (Sigma-Aldrich) and 1% pen/strep (100units/ mL penicillin, 100µg/ mL streptomycin; Gibco) at 37° C and 5% CO_2_. NIH3T3 cells were a kind gift from Dr. Caleb Bashor’s lab and were cultured in DMEM supplemented with 10% FBS (Sigma-Aldrich) and 1% pen/strep (100units/ mL penicillin, 100µg/ mL streptomycin) at 37° C and 5% CO_2_.

### Plasmid transfection and nucleofection

HEK293T cell transfections were performed in 24-well plates using 375ng of dCas9 expression plasmid and 125ng of equimolar pooled or individual gRNAs/crRNAs. 1.25×10^5^ HEK293T cells were plated the day before transfection and then transfected using Lipofectamine 3000 (Invitrogen, USA) as per manufacturer’s instruction. For two component systems (dCas9 + MCP or dCas9 + scFv systems) 187.5ng of each plasmid was used. For multiplex gene activation experiments using DREAM platforms, 25ng of each gRNA encoding plasmid targeting each respective gene was used. Transfections in HeLa, A549, SK-BR-3, U2OS, HCT-116, HFF, NIH3T3, and CHO-K1 were performed in 12-well plates using Lipofectamine 3000 and 375ng dCas9 plasmid, 375ng of MCP-effector fusion proteins, and 250ng DNA of MS2-modifed gRNA encoding plasmid. For transfections using dCas12a fusion proteins where single genes were targeted, 375ng of dCas12a-effector fusion plasmids and 125ng of crRNA plasmids were transfected using lipofectamine 3000 per manufacturer’s instruction. For multiplex gene activation experiments using dCas12a, 375ng of dCas12a-effector fusion encoding plasmid and 250ng of multiplex crRNA expression plasmids were used. For experiments using *E. coli* and *P. aeruginosa* Type I CRISPR systems, we followed the same stoichiometries used in previous studies^52, 53^. For transfection of *ICAM1*-ZF effectors, 500ng of each *ICAM1* targeting ZF fusion was transfected. Transfections using *IL1RN*-TALE fusion proteins were performed using 500ng of either single TALE or a pool of 4 TALEs using 125ng of each TALE fusion. All ZF and TALE transfections were performed in HEK293T cells in 24-well format using Lipofectamine 3000 as per manufacturers instruction. For K562 cells, 1×10^6^ cells were nucleofected using the Lonza SF Cell Line 4D-Nucleofector Kit (Lonza V4XC-2012) and a Lonza 4D Nucleofector (Lonza, AAF1002X) using the FF-120 program. 2000ng of total plasmids were nucleofected in each condition using 1×10^6^ K562 cells and 667ng each of; dCas9 plasmid, MCP fusion plasmid, and pooled MS2-sgRNA expression plasmid was nucleofected per condition. Immediately after nucleofection, K562 cells were transferred to prewarmed media containing 6-well plates. hTERT-MSCs were electroporated with using the Neon transfection system (Thermo Fisher Scientific) using the 100µL kit. 5×10^5^ hTERT-MSCs were resuspended in 100µL resuspension buffer R and 10µg total DNA (3.75µg dCas9, 3.75µg MCP-fusion effector plasmid, and 2.5µg MS2-modifed gRNA encoding plasmid). Electroporation was performed using the settings recommended by the manufacturers for mesenchymal stem cells: Voltage: 990V, Pulse width: 40ms, Pulse number: 1. For fibroblast reprogramming experiments, we used the Neon transfection system using the amounts of endotoxin free DNA described previously^17^ and below. Dual AAV (500ng of each) and All-in-one (AIO) AAV (1µg) construct transfections were performed in Neuro-2a cells in 12-well format using Lipofectamine 3000 as per manufacturers instruction.

### PBMC isolation, culture, and nucleofection

De-identified white blood cell concentrates (buffy coats) were obtained from the Gulf Coast Regional Blood Center in Houston, Texas. PBMCs were isolated from buffy coats using Ficoll gradient separation and cryopreserved in liquid nitrogen until later use. 1×10^6^ PBMCs per well were stimulated for 48h in a CD3/CD28 (Tonbo Biosciences, 700037U100 and 70289U100, respectively)-coated 24-well plate containing RPMI media supplemented with 10% FBS (Sigma-Aldrich), 1% Pen/Strep (Gibco), 10ng/mL IL-15 (Tonbo Biosciences, 218157U002), and 10ng/mL IL-7 (Tonbo Biosciences, 218079U002). Stimulated PBMCs were electroporated using the Neon transfection system (Thermo Fisher Scientific) 100μL kit per manufacturer protocol. Briefly, PBMCs were centrifuged at 300g for 5min and resuspended in Neon Resuspension Buffer T to a final density of 1×10^7^ cells/mL. 100µL of the resuspended cells (1×10^6^ cells) were then mixed with 12µg total plasmid DNA (4.5µg of dCas9 fusion encoding plasmids, 4.5µg of MCP fusion encoding plasmids, and 3µg of four equimolar pooled MS2-modifed gRNA encoding plasmids) and electroporated with the following program specifications using a 100μL Neon Tip: pulse voltage 2,150v, pulse width 20ms, pulse number 1. Endotoxin free plasmids were used in all experiments. After electroporation, PBMCs were incubated in prewarmed 6-well plates containing RPMI media supplemented with 10% FBS (Sigma-Aldrich), 1% Pen/Strep (Gibco), 10ng/mL IL-15, and 10ng/mL IL-7. PBMCs were maintained at 37°C, 5% CO_2_ for 48h before RNA isolation and QPCR.

### Human primary T cell and primary umbilical cord MSC culture and lentiviral transduction

PBMCs were isolated from de-identified white blood cell concentrates (buffy coats) using Ficoll gradient separation. T cells were isolated using negative selection via the EasySep™ Human T Cell Isolation Kit (StemCell, 17951). T cells were frozen in Bambanker Cell Freezing Media (Bulldog Bio Inc, BB01) and stored in liquid nitrogen until use. Umbilical cord derived MSCs (ATCC, PCS-500-010) were cultured in MSC basal media (ATCC, PCS-500-030) supplemented with Mesenchymal Stem Cell Growth Kit (PCS-500-040) containing rhFGF basic (5ng/mL), rhFGF acidic (5ng/mL), rhEGF (5ng/mL), FBS (2%), and L-Alanyl-L-Glutamine (2.4 mM). MSC media was also supplemented with 1% Pen-strep (Gibco, 15140122). MSCs were maintained at 37°C, 5% CO_2_. Lentiviral transduction was performed in stimulated T cells as previously described^82^. Briefly, 1×10^6^ T cells per well were stimulated for 24h with Dynabeads™ Human T-Activator CD3/CD28 for T Cell Expansion and Activation (Thermo Fisher Scientific, 11161D) according to manufacturer’s instructions in a 24-well plate containing X-VIVO 15 media (Lonza, 04418Q) supplemented with 5% FBS (Sigma-Aldrich), 55 mM 2-Mercaptoethanol (Gibco, 21985023), 4 mM N-acetyl-L-cysteine (Thermo Fisher Scientific, 160280250), and 500 IU/ml of recombinant human IL-2 (Biolegend, 589104). Stimulated T cells were co-transduced via spinoculation at 931xg, 37°C for 2 hours in a plate coated with Retronectin (Takara Bio, T100B) with an MOI of ∼5.0 for each lentivirus (dCas9 lentivirus at MOI ∼5.0 and gRNA-MCP-fusion effector lentivirus). After spinoculation, T cells were maintained at 37°C, 5% CO2 for 48h before downstream experiments. MSCs were co-transduced with an MOI of ∼10.0 (dCas9 lentivirus at MOI ∼10.0 and gRNA-MCP-fusion effector lentivirus at MOI ∼10.0) for each lentivirus via reverse transduction by seeding 1.25×10^5^ cells into each well of a 12-well plate containing the virus in MSC media supplemented with 8 μg/mL polybrene. Media was changed after 16 hours. Further experimental analyses were performed 72 hours post-transduction.

### Mouse primary neuron culture and AAV8 transduction

Mouse C57 Cortex Neurons (Lonza, M-CX-300) were cultured in Primary Neuron Basal Medium (PNBM) supplemented with 2mM L-glutamine, GA-1000 and 2 % NSF. In brief, 4×10^5^ cells were seeded in poly-D-lysine and laminin coated 24 well plates and cultured for 7 days for neuronal differentiation. On day 8, cells from each well were transduced with 1×10^10^ AAV8 viral particles (2.5×10^4^/cell). 5 days post-transduction cells were harvested for RNA isolation and QPCR analysis.

### Plasmid cloning

Lenti-dCas9-VP64 (Addgene #61425), dCas9-VPR (Addgene #63798), dCas9-p300 (Addgene #83889), MCP-p65-HSF1 (Addgene #61423), scFv-VP64 (Addgene #60904), SpgRNA expression plasmid (Addgene #47108), MS2-modified gRNA expression plasmid (Addgene #61424), AsCas12a (Addgene #128136), *E. Coli* Type I Cascade system (Addgene #106270-106275) and *Pae* Type I Cascade System (Addgene #153942 and 153943), YAP-S5A (Addgene #33093) have been described previously. The eNRF2 TAD fusion was synthetically designed and ordered as a gBlock from IDT. To generate an isogenic C-terminal effector domain cloning backbone, the dCas9-p300 plasmid (Addgene #83889) was digested with BamHI and then a synthetic double-stranded ultramer (IDT) was incorporated using NEBuilder HiFi DNA Assembly (NEB, E2621) to generate a dCas9-NLS-linker-BamHI-NLS-FLAG expressing plasmid. This plasmid was further digested with AfeI and then a synthetic double-stranded ultramer (IDT) was incorporated using NEBuilder HiFi DNA Assembly to generate a FLAG-NLS-MCS-linker-dCas9 expressing Plasmid for N-terminal effector domain cloning. For fusion of effector domains to MCP, the MCP-p65-HSF1 plasmid (Addgene #61423) was digested with BamHI and NheI and respective effector domains were cloned using NEBuilder HiFi DNA Assembly. For SunTag components, the scFv-GCN4-linker-VP16-GB1-Rex NLS sequence was PCR amplified from pHRdSV40-scFv-GCN4-sfGFP-VP64-GB1-NLS (Addgene #60904) and cloned into a lentiviral backbone containing an EF1-alpha promoter. Then VP64 domain was removed and an AfeI restriction site was generated and used for cloning TADs using NEBuilder HiFi DNA Assembly. The pHRdSV40-dCas9-10xGCN4_v4-P2A-BFP (Addgene #60903) vector was used for dCas9-based scFv fusion protein recruitment to target loci. All MTF TADs were isolated using PCR amplified from a pooled cDNA library from HEK293T, HeLa, U2OS and Jurkat-T cells. TADs were cloned into the MCP, dCas9 C-terminus, dCas9 N-terminus, and scFv backbones described above using NEBuilder HiFi DNA Assembly. Bipartite N-terminal fusions between MCP-MRTF-A or MCP-MRTF-B TADs and STAT 1-6 TADs were generated by digesting the appropriate MCP-fusion plasmid (MCP-MRTF-A or MCP-MRTF-B) with BamHI and then subcloning PCR-amplified STAT 1-6 TADs using NEBuilder HiFi DNA Assembly. Bipartite C-terminal fusions between MCP-MRTF-A or MCP-MRTF-B TADs and STAT 1-6 TADs were generated by digesting the appropriate MCP-fusion plasmid (MCP-MRTF-A or MCP-MRTF-B) with NheI and then subcloning PCR-amplified STAT 1-6 TADs using NEBuilder HiFi DNA Assembly. Similarly, eNRF2 was fused to the N- or C-terminus of the bipartite MRTF-A-STAT1 TAD in the MCP-fusion backbone using either BamHI (N-terminal; MCP-eNRF2-MRTF-A-STAT1 TAD) or NheI (C-terminal; MCP-MRTF-A-STAT1-eNRF2 TAD) digestion and NEBuilder HiFi DNA Assembly to generate the MCP-NMS or MCP-MSN tripartite TAD fusions, respectively. SadCas9 (with D10A and N580A mutations derived using PCR) was PCR amplified and then cloned into the SpdCas9 expression plasmid backbone created in this study digested with BamHI and XbaI. This SadCas9 expression plasmid was digested with BamHI and then PCR-amplified VP64 or VPR TADs were cloned in using NEBuilder HiFi DNA Assembly. CjCas9 was PCR-amplified from pAAV-EFS-CjCas9-eGFP-HIF1a (Addgene #137929) as two overlapping fragments using primers to create D8A and H559A mutations. These two CjdCas9 PCR fragments were then cloned into the SpdCas9 expression plasmid digested with BamHI and XbaI using NEBuilder HiFi DNA Assembly. This CjdCas9 expression plasmid was digested with BamHI and the PCR-amplified VP64 or VPR TADs were cloned in using NEBuilder HiFi DNA Assembly. HNH domain deleted SpdCas9 plasmids were generated using different primer sets designed to amplify the N-terminal and C-terminal portions of dCas9 excluding the HNH domain and resulting in either: no linker, a glycine-serine linker, or an XTEN16 linker, between HNH-deleted SpdCas9 fragments. These different PCR-amplified regions were cloned into the SpdCas9 expression plasmid digested with BamHI and XbaI using NEBuilder HiFi DNA Assembly. MCP-mCherry, MCP-MSN and MCP-p65-HSF1 were digested with NheI and a single strand oligonucleotide encoding the FLAG sequence was cloned onto the C-terminus of each respective fusion protein using NEBuilder HiFi DNA Assembly to enable facile detection via Western blotting. 1x 9aa TADs were designed and annealed as double strand oligos and then cloned into the BamHI/NheI-digested MCP-p65-HSF1 backbone plasmid (Addgene #61423) using T4 ligase (NEB). Heterotypic 2x 9aa TADs were generated by digesting MCP-1x 9aa TAD plasmids with either BamHI or NheI and then cloning single strand DNA encoding 1x 9aa TADs to the N- or C-termini using NEBuilder HiFi DNA Assembly. Heterotypic MCP-3x 9aa TADs were generated similarly by digesting MCP-2x 9aa TAD containing plasmids either with BamHI or NheI and then single strand DNA encoding 1x 9aa TADs were cloned to the N- or C-termini using NEBuilder HiFi DNA Assembly. Selected fusions between 3x 9aa TADs and eNRF2 were generated using gBlock (IDT) fragments and cloned into the BamHI/NheI-digested MCP-p65-HSF1 backbone plasmid (Addgene #61423) using NEBuilder HiFi DNA Assembly. To generate mini-DREAM compact single plasmid system, SpdCas9-HNH (no linker) deleted plasmid was digested with BamHI and then PCR amplified P2A self-cleaving sequence and MCP-eNRF2-3x 9aa TAD (eN3×9) was cloned using NEBuilder HiFi DNA Assembly. For dCas12a fusion proteins, SiT-Cas12a-Activ (Addgene #128136) was used. First, we generated a nuclease dead (E993A) SiT-Cas12a backbone using PCR amplification and we used this plasmid for subsequent C-terminal effector cloning using BamHI digestion and NEBuilder HiFi DNA Assembly. For *E. coli* Type I CRISPR systems, the Cas6-p300 plasmid (Addgene #106275) was digested with BamHI and then MSN and NMS domains were cloned in using NEBuilder HiFi DNA Assembly. *Pae* Type I Cascade plasmids encoding Csy1-Csy2 (Addgene #153942) and Csy3-VPR-Csy4 (Addgene #153943) were obtained from Addgene. The Csy3-VPR-Csy4 plasmid was digested with MluI (NEB) and BamHI (to remove the VPR TAD) and then the nucleoplasmin NLS followed by a linker sequence was added using NEBuilder HiFi DNA Assembly. Next, this Csy3-Csy4 plasmid was digested with AscI and either the MSN or NMS TADs were cloned onto the N-terminus of Csy3 NEBuilder HiFi DNA Assembly. ZF fusion proteins were generated by cloning PCR-amplified MSN, NMS, or VPR domains into the BsiWI and AscI digested *ICAM1* targeting ZF-p300 plasmid^10^ using NEBuilder HiFi DNA Assembly. Similarly, TALE fusion proteins were created by cloning PCR-amplified MSN, NMS, or VPR domains into the BsiWI and AscI digested *IL1RN* targeting TALE plasmid backbone^10^ using NEBuilder HiFi DNA Assembly. pCXLE-dCas9VP192-T2A-EGFP-shP53 (Addgene #69535), GG-EBNA-OSK2M2L1-PP (Addgene #102898) and GG-EBNA-EEA-5guides-PGK-Puro (Addgene #102898) used for reprogramming experiments have been described previously^17, 83^. The PCR-amplified NMS domain was cloned into the sequentially digested (XhoI then SgrDI; to remove the VP192 domain) pCXLE-dCas9VP192-T2A-EGFP-shP53 backbone using NEBuilder HiFi DNA Assembly. TADs were directly fused to the C-terminus of dCas9 by digesting the dCas9-NLS-linker-BamHI-NLS-FLAG plasmid with BamHI and then cloning in PCR-amplified TADs using NEBuilder HiFi DNA Assembly. TADs were directly fused to the N-terminus to dCas9 by digesting the FLAG-NLS-MCS-linker-dCas9 plasmid with AgeI (NEB) and then cloning in PCR-amplified TADs using NEBuilder HiFi DNA Assembly. For constructs harboring both N- and C-terminal fusions, respective plasmids with TADs fused to the C-terminus of dCas9 were digested with AgeI and then PCR-amplified TADs were cloned onto the N-terminus of dCas9 using NEBuilder HiFi DNA Assembly. hSyn-AAV-EGFP (Addgene #50465) plasmid was used to generate different AAV based DNA constructs. For SpdCas9 cloning both EFGP and WPRE were removed using XbaI and XhoI and SpdCas9 and the modified smaller WPRE along with SV40 polyA signal (W3SL) were then cloned into this backbone using NEBuilder HiFi DNA Assembly. For expression of MS2-gRNA and hSyn-MCP-MSN from a single plasmid, both components were PCR amplified and cloned into an EGFP-removed hSyn-AAV-EGFP backbone using NEBuilder HiFi DNA Assembly. For All-In-One AAV backbone the M11 promoter^84^ was used to drive SaCas9 gRNA expression. The SCP1^85^ and the EFS promoters were used to drive the expression of NMS-SadCas9. The efficient, smaller synthetic WPRE and polyadenylation signal CW3SA^66^ was utilized to maximize expression this size-limited context. Following cloning and sequence verification, 3 SaCas9 specific gRNAs targeting mouse *Agrp* gene were cloned into the all-in-one (AIO) vectors using Bbs1 restriction digestion. Following identification of the most efficacious gRNA (by transfecting into Neuro-2a cells), the SCP1 and EFS promoter driven SadCas9 based AIO plasmids were sequence verified by Plasmidsaurus. Sequence verified SpdCas9 and SCP1 and EFS promoter driven SadCas9 based AIO plasmids were sent to Charles River Laboratories for AAV8 production. Titers of different AAVs are included in source data.

### gRNA design and construction

All protospacer sequences for SpCas9 systems were designed using the Custom Alt-R® CRISPR-Cas9 guide RNA design tool (IDT). All gRNA protospacers were then phosphorylated, annealed, and cloned into chimeric U6 promoter containing sgRNA cloning plasmid (Addgene #47108) and/or an MS2 loop containing plasmid backbone (Addgene #61424) digested with Bbs1 and treated with alkaline phosphatase (Thermo) using T4 DNA ligase (NEB). The SaCas9 gRNA expression plasmid (pIBH072) was a kind gift from Charles Gersbach and was digested with BbsI or Bpil (NEB or Thermo, respectively) and treated with alkaline phosphatase and then annealed protospacer sequences were cloned in using T4 DNA ligase (NEB). gRNAs were cloned into the pU6-Cj-sgRNA expression plasmid (Addgene #89753) by digesting the vector backbone with BsmBI or Esp3I (NEB or Thermo, respectively), and then treating the digested plasmid with alkaline phosphatase, annealing phosphorylated gRNAs, and then cloning annealed gRNAs into the backbone using T4 DNA ligase. MS2-stem loop containing plasmids for SaCas9 and CjCas9 were designed as gBlocks (IDT) with an MS2-stem loop incorporated into the tetraloop region for both respective gRNA tracr sequences. crRNA expression plasmids for the Type I *Eco* Cascade system were generated by annealing synthetic DNA ultramers (IDT) containing direct repeats (DRs) and cloning these ultramers into the BbsI and SacI-digested SpCas9 sgRNA cloning plasmid (Addgene #47108) using NEBuilder HiFi DNA Assembly. crRNA expression plasmids for *Pae* Type I Cascade system were generated by annealing and then PCR-extending overlapping oligos (that also harbored a BsmBI or Esp3I cut site for facile crRNA array incorporation) into the sequentially BbsI (or Bpil) and SacI-digested SpCas9 sgRNA cloning plasmid (Addgene #47108) using NEBuilder HiFi DNA Assembly. crRNA expression plasmids for Cas12a systems were generated by annealing and then PCR-extending overlapping oligos (that also harbored a BsmBI or Esp3I cut site for facile crRNA array incorporation) into the sequentially BbsI (or Bpil) and SacI-digested SpCas9 sgRNA cloning plasmid (Addgene #47108) using NEBuilder HiFi DNA Assembly. All gRNA spacer sequences used in this study for SpdCas9, SadCas9, and CjdCas9 are listed in **Supplementary Tables 1, 2, and 3,** respectively.

### crRNA array cloning

crRNA arrays for AsCas12a and Type I CRISPR systems were designed in fragments as overlapping ssDNA oligos (IDT) and 2-4 oligo pairs were annealed. Oligos were designed with an Esp3I cut site at 3’ of the array for subsequent cloning steps. Equimolar amounts of oligos were mixed, phosphorylated, and annealed similar to the standardized gRNA/crRNA assembly protocol above. Phosphorylated and annealed arrays were then cloned into the respective Esp3I-digested and alkaline phosphatase treated crRNA cloning backbone (described above) using T4 DNA ligase (NEB). crRNA arrays were verified by Sanger sequencing. Correctly assembled 4-8 crRNA array expressing plasmids were then digested again with Esp3I and alkaline phosphatase treated to enable incorporation of subsequent arrays up to 20 crRNAs. All crRNA spacer sequences used in this study for AsdCas12a, *Eco*-Cascade, and *Pae*-Cascade are listed in **Supplementary Tables 4, 5, and 6,** respectively.

### Lentiviral packaging

All lentiviral transfer and packaging plasmids were purified using the Endofree Plasmid Maxi Kit (Qiagen, 12362). Lentivirus was packaged as previously described^82^ with minor modifications. Briefly, HEK293T cells were seeded into 225mm flasks and maintained in DMEM. OptiMem was used for transfection and Sodium butyrate was added to a final concentration of 4mM. Lentivirus was then concentrated 100X using the Lenti-X concentrator (Takara Bio,631232). Biological titration of lentivirus by QPCR was carried out as previously described^86^, with the following modifications. Volumes of 10, 5, 1, 0.1, 0.01, and 0 µl of concentrated lentiviral particles were reverse transduced into 5×10^4^ HEK293T cells with 8μg/mL polybrene (Millipore-Sigma, TR1003G) in 24 well format with media exchanged after 14 hours of transduction. gDNA was extracted 96 hours post-transduction using the DNeasy Blood & Tissue Kit (Qiagen, 69506). qPCR was performed using 67.5ng of gDNA for each condition in 10µl reactions using Luna Universal qPCR Master Mix (NEB, M3003E).

### Western blotting

Cells were lysed in RIPA buffer (Thermo Scientific, 89900) with 1X protease inhibitor cocktail (Thermo Scientific, 78442), lysates were cleared by centrifugation and protein quantitation was performed using the BCA method (Pierce, 23225).15-30µg of lysate were separated using precast 7.5% or 10% SDS-PAGE (Bio-Rad) and then transferred onto PVDF membranes using the Transblot-turbo system (Bio-Rad). Membranes were blocked using 5% BSA in 1X TBST and incubated overnight with primary antibody (anti-Cas9; 1:1000 dilution, Diagenode #C15200216, Anti-FLAG; 1:2000 dilution, Sigma-Aldrich #F1804, anti-β-Tubulin; 1:1000 dilution, Bio-Rad #12004166). Then membranes were washed with 1X TBST 3 times (10mins each wash) and incubated with respective HRP-tagged secondary antibodies (1:2000 dilution) for 1hr. Next membranes were washed with 1X TBST 3 times (10mins each wash). Membranes were then incubated with ECL solution (BioRad # 1705061) and imaged using a Chemidoc-MP system (BioRad). The β-tubulin antibody was tagged with Rhodamine (Bio-Rad #12004166) and was imaged using Rhodamine channel in Chemidoc-MP as per manufacturer’s instruction.

### Quantitative reverse-transcriptase PCR (QPCR)

RNA (including pre-miRNA) was isolated using the RNeasy Plus mini kit (Qiagen #74136). 500-2000ng of RNA (quantified using Nanodrop 3000C; Thermo Fisher) was used as a template for cDNA synthesis (Bio-Rad #1725038). cDNA was diluted 10X and 4.5µL of diluted cDNA was used for each QPCR reaction in 10µL reaction volume. Real-Time quantitative PCR was performed using SYBR Green mastermix (Bio-Rad #1725275) in the CFX96 Real-Time PCR system with a C1000 Thermal Cycler (Bio-Rad). Results are represented as fold change above control after normalization to *GAPDH* in all experiments using human cells. For murine cells, 18s rRNA was used for normalization. For CHO-K1 cells, *Gnb1* was used for normalization. Undetectable samples were assigned a Ct value of 45 cycles. All QPCR primers and cycling conditions are listed in **Supplementary Table 7.**

### Mature miRNA isolation and QPCR for miRNAs

Mature miRNA (miRNA) was isolated using the miRNA isolation kit (Qiagen #217084). 500ng of isolated miRNA was polyadenylated using poly A polymerase (Quantabio #95107) in 10µL reactions per sample and then used for cDNA synthesis using qScript Reverse Transcriptase and oligo-dT primers attached to unique adapter sequences to allow specific amplification of mature miRNA using QPCR in a total 20µL reaction (Quantabio #95107). cDNA was diluted and 10ng of miRNA cDNA was used for QPCR in a 25µL reaction volume. PerfeCTa SYBR Green SuperMix (Quantabio #95053), *miR-146a* specific forward primer, and PerfeCTa universal reverse primer was used to perform QPCR. U6 snRNA was used for normalization. All QPCR primers and cycling conditions are listed in **Supplementary Table 7.**

### Immunofluorescence microscopy

Human foreskin fibroblasts (HFFs; CRL-2429, ATCC) and HFF-derived iPSCs were grown in Geltrex (Gibco, A1413302) coated 12-well plates and were fixed with 3.7% formaldehyde and then blocked with 3% BSA in 1X PBS for 1hr at Room Temperature prior to imaging. Primary antibodies for SSEA-4 (CST #43782), TRA1-60 (CST #61220) and TRA1-81 (CST #83321) were diluted (1:200) in 1% BSA in 1X PBS and incubated overnight at 4°C. The next day, cells were washed with 1X PBS, incubated with appropriate Alexaflour-488 conjugated secondary antibodies (1:500) for 1hr at Room Temperature and then washed again 3 times with 1X PBS. Cells were then incubated with DAPI (Invitrogen #D1306) containing PBS (100nM final concentration) for 10m, washed 3 times with 1X PBS, and then imaged using a Nikon ECLIPSE Ti2 fluorescent microscope.

### CD34 surface expression analysis

Surface staining of CD34 in HEK293T cells was performed using CD34-PE antibody (Invitrogen, #MA1-10205). In brief, 72 hours post-transfection, cells from 24 well plates were detached using TrypLE select (Gibco, #12563011). Single cell suspensions were washed with complete media and then with 0.22 um filtered 1X FACS Buffer (1% BSA in 1X PBS). Next, cells were incubated with CD34-PE antibody or IgG-PE isotype antibody (Invitrogen, #12-4714-42) in 1X FACS Buffer for 30 mins. Stained cell fluorescence intensity was measured using a Sony SA3800 spectral analyzer. To assess the CD34 expression in EGFP positive MCP-effector transfected cells, single cells were gated based on EGFP expression and assessed for CD34 expression. Data were analyzed using FlowJo software (V10). First cell debris and doublets were excluded with the FSC-A/SSC-A dot plot, followed by FSC-H/FSC-A dot plot. Events were excluded with intensity <1X 10^-2^, using FSC-A/PE-A dot plots. The CD34 surface marker-positive cells were gated using unstained or isotype controls. For cells transfected with an EGFP expressing effector construct, similar gating strategy was used to select singles cells as above and then events were excluded with intensity <1X 10^-2^, using PE-A/EGFP-A dot plots, then EGFP expressing cells were analyzed for CD34 surface marker expression using unstained or isotype controls.

### CUT&RUN

CUT&RUN was performed using the Epicypher CUTANA ChIC/CUT&RUN Kit (Epicypher, #14-1048). Briefly, transfected cells were detached and harvested using TrypLE select (Gibco, #12563011), washed once with 1X PBS and then dissolved in 300ul of wash buffer. Next each of 3 100ul aliquots (∼1/3 of each 24 well) of cells were processed for H3K4me3 antibody (Epicypher, #13-0041), H3K27ac antibody (Epicypher, #13-0045) or input DNA, respectively. Cells were first immobilized on concanavalin A beads, and then incubated with respective antibody (0.5ug/sample) overnight at 4°C in antibody dilution buffer (cell permeabilization buffer + EDTA). On the following day, cells were washed twice with cell permeabilization buffer. After washing the beads, pAG-MNase was added to the immobilized cells and then incubated for 2 hours at 4°C to digest and release DNA. For CUT&RUN-qPCR assays, purified DNA from both H3K4me3 antibody and H3K27ac antibody incubated samples was then assayed using QPCR. Relative enrichment of H3K4me3 and H3K27ac is expressed as fold change above control cells transfected with dCas9+MCP-mCherry plasmid and after normalization to purified input DNA. QPCR primers used for CUT&RUN are shown in **Supplementary Table 8**.

### Generation of mini-DREAM component expressing HEK293T cell line

HEK293T cells were co-transduced with HNH-deleted dCas9 and MCP-eN3×9-T2A-EGFP lentiviruses (each with an MOI of ∼5.0) using 8μg/mL polybrene (Millipore-Sigma, TR1003G) in 24 well format. Media was exchanged after 14 hours post-transduction. Mini-DREAM HEK293T cells were then transfected with gRNA/gRNA array for further experimentation.

### Progesterone ELISA

Secreted progesterone was measured using the Progesterone Competitive ELISA Kit (Invitrogen #EIAP4C21). In brief, 72 hours post-transfection of control gRNA or the indicated MS2-gRNA array into a mini-DREAM expressing HEK293T cell line. 50µl of cell culture supernatant was directly used for ELISA as per manufacturer’s instruction, along with all recommended progesterone standards. Standard curves were generated using polynomial function and progesterone concentration was determined and expressed in ng/mL.

### Fibroblast reprogramming

HFFs were cultured in 1X DMEM supplemented with 1X Glutamax (Gibco, 35050061) for two passages before transfection with respective components. Cells were grown in 15cm dishes (Corning), and detached using TrypLE select (Gibco, #12563011). Single cell suspensions were washed with complete media and then with 1X PBS. For each 1 x 10^6^ cells, a total of 6µg of endotoxin free plasmids (Macherey-Nagel, 740424; 2µg CRISPR activator plasmid, 2µg of pluripotency factor targeting gRNA plasmid, and 2µg of EEA-motif targeting gRNA expression plasmids) were nucleofected using a 100µL Neon transfection tip in R buffer using the following settings: 1650V, 10ms, and 3 pulses. Nucleofected fibroblasts were then immediately transferred to Geltrex (Gibco) coated 10cm cell culture dishes in prewarmed media. The next day media was exchanged. 4 days later, media was replaced with iPSC induction media^17^. Induction media was then exchanged every other day for 18 days. After 18 days iPSC colonies were counted, and colonies picked using sterile forceps and then transferred to Geltrex coated 12-well plates. iPSC colonies were maintained in complete E8 media and passaged as necessary using ReLeSR passaging reagent (Stem Cell Technology, #05872). RNA was isolated from iPSC clones using the RNeasy Plus mini kit (Qiagen #74136) and colonies were immunostained using indicated antibodies and counterstained with DAPI (Invitrogen) for nuclear visualization.

### RNA sequencing (RNA-seq)

RNA-seq was performed in duplicate for each experimental condition. 72 hours post-transfection RNA was isolated using the RNeasy Plus mini kit (Qiagen). RNA integrity was first assessed using a Bioanalyzer 2200 (Agilent) and then RNA-seq libraries were constructed using the TruSeq Stranded Total RNA Gold (Illumina, RS-122-2303). The qualities of RNA-seq libraries were verified using the Tape Station D1000 assay (Tape Station 2200, Agilent Technologies) and the concentration of RNA-seq libraries were checked again using real time PCR (QuantStudio 6 Flex Real time PCR System, Applied Biosystem). Libraries were normalized and pooled prior to sequencing. Sequencing was performed using an Illumina Hiseq 3000 with paired end 75 base pair reads. Reads were aligned to the human genome (hg38) Gencode Release 36 reference using STAR aligner (v2.7.3a). Transcript levels were quantified to the reference genome using a Bayesian approach. Normalization was done using counts per million (CPM) method. Differential expression was done using DESeq2 (v3.5) with default parameters. Genes were considered significantly differentially expressed based upon a fold change >2 or <-2 and an FDR <0.05.

### 9aa TAD prediction

9aa TADs were predicted using previously described software (http://www.at.embnet.org/toolbox/9aatad/.)^38^ using the “moderately stringent pattern” criteria and all “refinement criteria” and only TADs with 100% matches were then selected for evaluation in MCP fusion proteins.

### Toxicity assays

Cellular toxicity assays in primary T cells were performed 72 hours post-transduction using the Annexin V:PE Apoptosis Detection Kit (BD Biosciences, 559763). In brief, cells were stained with 7-AAD and Annexin V:PE according to the manufacturer’s protocol. Stained cell fluorescence was measured using a Sony SA3800 spectral analyzer. EGFP positive single cells were gated and assessed for 7-AAD and Annexin V: PE fluorescence. All conditions were measured in biological triplicate and measured in technical duplicate. The toxicity of treatment groups was compared to the negative control (dCas9 alone), camptothecin (5mM), and 65^0^C heat shock were used as positive controls of apoptosis and membrane permeability respectively. Cellular toxicity assays in HEK293T and U2OS cells were performed using Hoechst and 7-AAD staining followed by microscopy. In brief, 48 hours post-transfection of different CRISPRa tools, media was aspirated from each 24 well and 150µl of staining solution was gently added to cover the cells in each condition. The staining solution contained Hoechst 33342 (Thermo Scientific, #62249) diluted 1:6000 and 5 µl of 7-AAD (BD Biosciences, #51-68981E) in sterile 1X PBS. 24 well plates were then incubated for 30 minutes at RT while protected from light. After incubation, automated images were taken of cell using a Nikon Ti2-E inverted microscope equipped with an Andor Zyla 4.2 sCMOS camera and 488nm and 561nm lasers using the 10 X PLAN APO λD objective.

### Data analysis

All data used for statistical analysis had a minimum of 3 biological replicates. Data are presented as mean ± SEM Gene expression analyses were conducted using Student’s t-tests (Two-tailed pair or multiple unpaired). Results were considered statistically significant when the *P*-value was <0.05. All bar graphs, error bars, and statistics were generated using GraphPad Prism v 9.0.

## Data Availability

Selected CRISPR/Cas constructs and fusions are available through Addgene, and all reagents are available from the authors upon request. All RNA sequencing data are available through the NCBI.

## Acknowledgement

The authors thank Jacopo de Rossi and Dhiraj Jain for their assistance with 9aa TADs prediction and cloning. The authors thank Arijita Sarkar for her critical help with NGS data analysis and the selection of miRNA target loci in HEK293T cells. The authors thank Harshavardhan Deshmukh and Suchir Mishra for their assistance with flow cytometry and protein sequence analyses, respectively. The authors also thank Adrian Picker-Oliver and Charles A. Gersbach for providing *E. coli* Type I CRISPR Cascade plasmids. The authors thank all members of the Hilton lab for helpful discussions and insights.

## Author Contributions

B.M. and I.B.H. conceived the project and designed experiments. B.M performed most experiments and analyzed the data with the assistance of all authors. B.M. and I.B.H. wrote the manuscript with input from all authors.

