## Supplemental Information for "Compact engineered human mechanosensitive transactivation modules enable potent and versatile synthetic transcriptional control"

#### Supplementary Note 1

Although single 9aa TADs derived from MRTF-A, MRTF-B or MYOCD were unable to substantially induce transcription (**Supplementary Fig. 17d**), we hypothesized that the combinatorial fusion of multiple 9aa TADs could do so. To test this hypothesis, we generated heterotypic bipartite 9aa TAD fusions derived from full-length TADs separated by a 1x glycine-serine linker. Our data showed that fusion of two distinct 9aa TADs from MYOCD.1 and MYOCD.3 respectively, modestly induced transcription (**Supplementary Fig. 17e**). We used this bipartite fusion for further testing. We fused different 9aa TADs derived from MRTF-A or MRTF-B to the N-terminus or C-terminus of the 2x 9aa TAD MYOCD.1-GS-MYOCD.3, and interestingly observed that all heterotypic 3x 9aa TADs strongly induced transcription, with a 3x 9aa TAD configuration of MRTF-B.3-GS-MYOCD.1-GS-MYOCD.3 displaying the highest transcriptional induction from our *OCT4* testbed locus (**Supplementary Fig. 17f**). To assess the effectiveness of this 3x 9aa TAD more generally, we targeted endogenous protein coding genes using pooled gRNAs (*HBG1*, *TTN and CD34*) and single gRNAs (*SBNO2*). We also targeted this 3x 9aa TAD, to the *GRASLND* lncRNA and the *OCT4-DE* enhancer using pools of gRNAs and observed substantial transactivation at all tested loci (**Fig. 3g and Supplementary Fig. 17g-17j**).

**Supplementary Note 2**

Direct fusion of effectors to dCas9 is attractive because it enables less complex delivery. However, many effectors display optimal performance when recruited by dCas9 using other architectures (e.g., SunTag/MCP; **Supplementary Figs. 2a and b, 4a and b and 8b**). That is, some effectors appear to lose efficacy when directly fused to dCas9. Therefore, we sought to test the relative efficacy of MSN and NMS in direct dCas9 fusion formats relative to similar directly fused architectures such as dCas9-VPR. Although, direct fusion of MSN or NMS to the C- or N-terminus of dCas9 enabled robust activation of target genes, efficacy with any of these formats remained lower than that of dCas9-VPR (**Supplementary Fig. 24a**). To test if we could further boost the potency of MSN/NMS direct fusions to dCas9, we added the VP64 domain in various locations and, interestingly observed highly potent transcriptional activation comparable to that of dCas9-VPR at the *OCT4* testbed locus. Moreover, the size of the effector components in the best performing fusion protein; NMS-dCas9-VP64 (1,020bp) is ~35% smaller than the size of the VPR effector (1,569bp) in dCas9-VPR (**Supplementary Fig. 24d**). We also noted that the relative expression of NMS-dCas9-VP64 was qualitatively higher than dCas9-VPR in HEK293T cells using Western blotting (**Supplementary Fig. 24b**). NMS-dCas9-VP64 performed comparably to dCas9-VPR at a battery of endogenous loci in HEK293T cells using pool of gRNAs and single gRNAs (**Supplementary Fig. 24e and f,** respectively).

#### Supplementary Figure 1

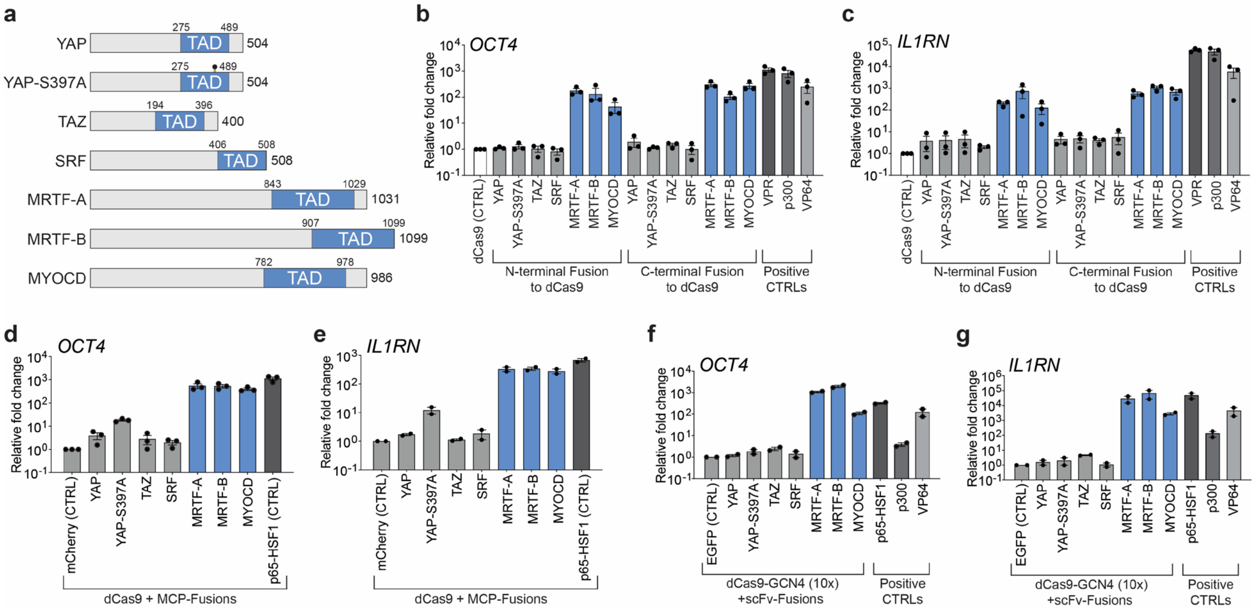


**Supplementary Fig. 1. Transactivation potency of serum responsive MTF TADs when recruited to human promoters via dCas9. a.** Schematics showing Hippo and SRF-MRTF family proteins; YAP, the hyperactive YAP mutant (YAP S397A), TAZ, SRF, MRTF-A, MRTF-B and MYOCD proteins. TADs for each respective MTF, along with amino acid (aa) coordinates, are shown in light blue. **b and c.** *OCT4* and *IL1RN* mRNA levels after the indicated TADs were fused to the N- or C-terminus of dCas9 targeted to each respective promoter using 4 pooled gRNAs. **d and e.** *OCT4* and *IL1RN* mRNA levels after the indicated TADs were fused to the MCP protein and recruited via 4 pooled MS2 modified gRNAs and dCas9. MCP fused to the bipartite p65-HSF was used as a positive control. **f and g.** *OCT4* and *IL1RN* mRNA levels after the indicated TADs were fused to scFv and recruited via dCas9 harboring 10xGCN4 C-terminal fusion protein (the SunTag system) along with 4 pooled standard gRNAs targeting each respective promoter. All samples were processed for QPCR analysis 72 hours post-transfection in HEK293T cells and are the result of at least 2 biological replicates. Error bars; SEM.

#### Supplementary Figure 2

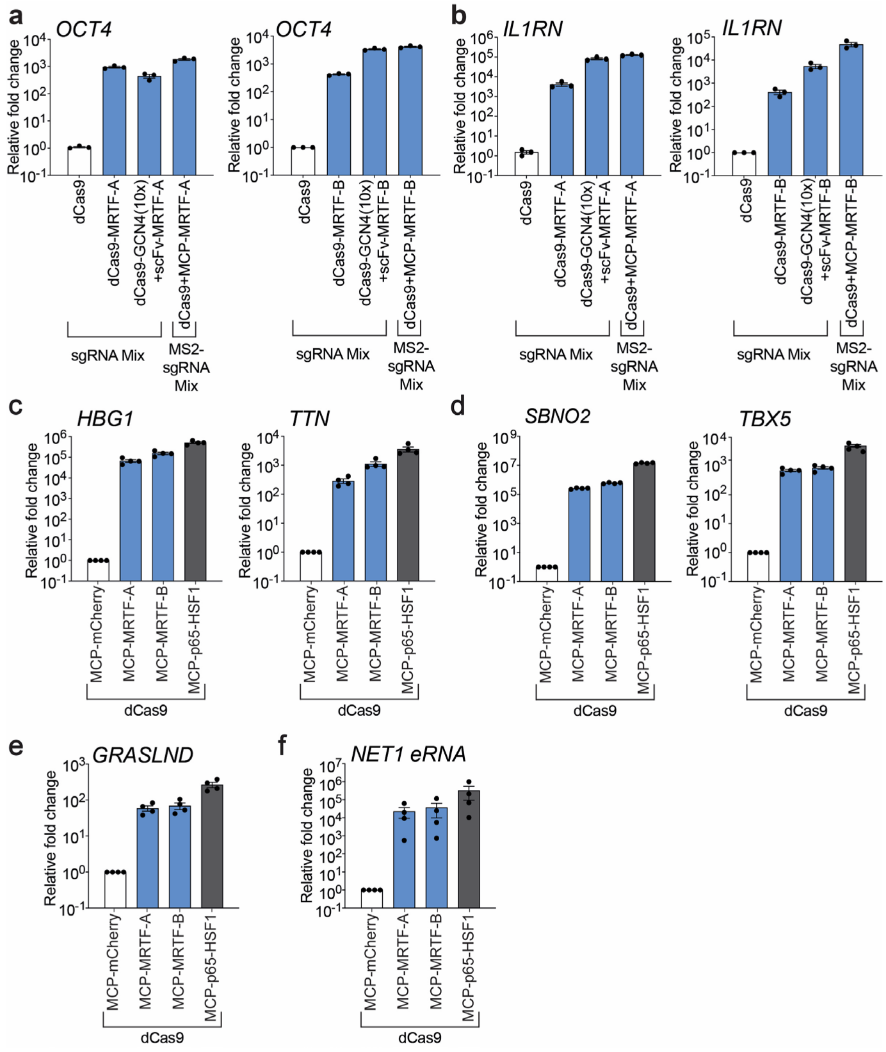


**Supplementary Fig. 2. Comparison and versatility of gene activation potential between TADs from MRTF-A and MRTF-B. a and b.** *OCT4* and *IL1RN* mRNA levels after the indicated TADs from MRTA-A (left panels) or MRTF-B (right panels) were recruited using the specified dCas9-based recruitment architecture (direct fusion, SunTag-based, or MCP-based) and 4 corresponding pooled gRNAs. **c and d.** mRNA levels for indicated loci after TADs from MRTF-A or MRTF-B were recruited via dCas9 and targeted to promoters using pools of MS2 modified gRNAs (**panel c**; *HBG1* and *TTN* promoters) or a single gRNA (**panel d**; *SBNO2* and *TBX5* promoters). **e and f**. *GRASLND* long noncoding RNA (**left**) or *NET1* eRNA (**right**) levels after TADs from MRTF-A or MRTF-B were recruited via dCas9 and targeted to each respective locus using 4 and 2 pooled MS2 modified gRNAs, respectively. All samples were processed for QPCR analysis 72 hours post-transfection in HEK293T cells and are the result of at least 2 biological replicates. Error bars; SEM.

#### Supplementary Figure 3

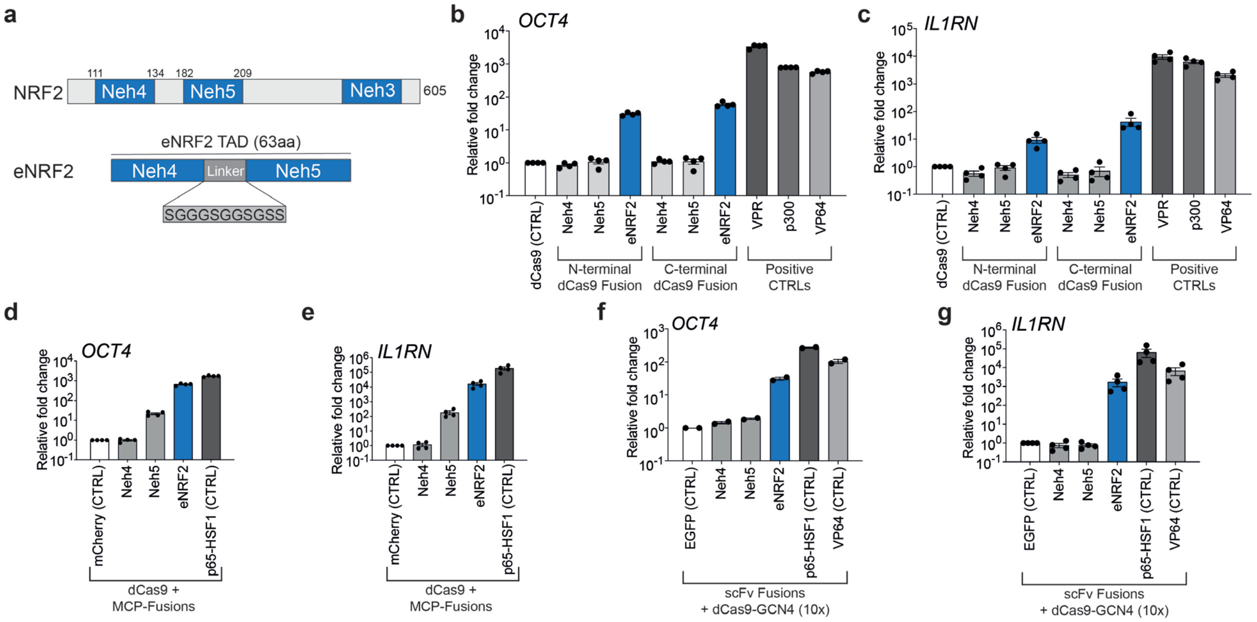


**Supplementary Fig. 3. Transactivation potency of oxidative stress responsive MTF TADs when recruited to human promoters via dCas9. a.** TADs from the NRF2 protein are depicted schematically. eNRF2 is a fusion between Neh4 and Neh5 TADs separated by an 11 amino acid (aa) extended glycine-serine linker. The length of NRF2 and aa coordinates of individual TADs (Neh4 and Neh5) are shown. **b and c.** *OCT4* and *IL1RN* mRNA levels after the indicated TADs were fused to the N- or C-terminus of dCas9 and targeted to each respective promoter using 4 pooled standard gRNAs. **d and e.** *OCT4* and *IL1RN* mRNA levels after the indicated TADs were fused to the MCP protein and recruited via 4 pooled MS2 stem-loop modified gRNAs and dCas9. MCP fused to the bipartite p65-HSF was used as a positive control. **f and g.** *OCT4* and *IL1RN* mRNA levels after the indicated TADs were fused to scFv and recruited via dCas9 harboring 10xGCN4 C-terminal fusion protein (the SunTag system) along with 4 pooled standard gRNAs targeting each respective promoter. All samples were processed for QPCR analysis 72 hours post-transfection in HEK293T cells and are the result of at least 2 biological replicates. Error bars; SEM.

#### Supplementary Figure 4

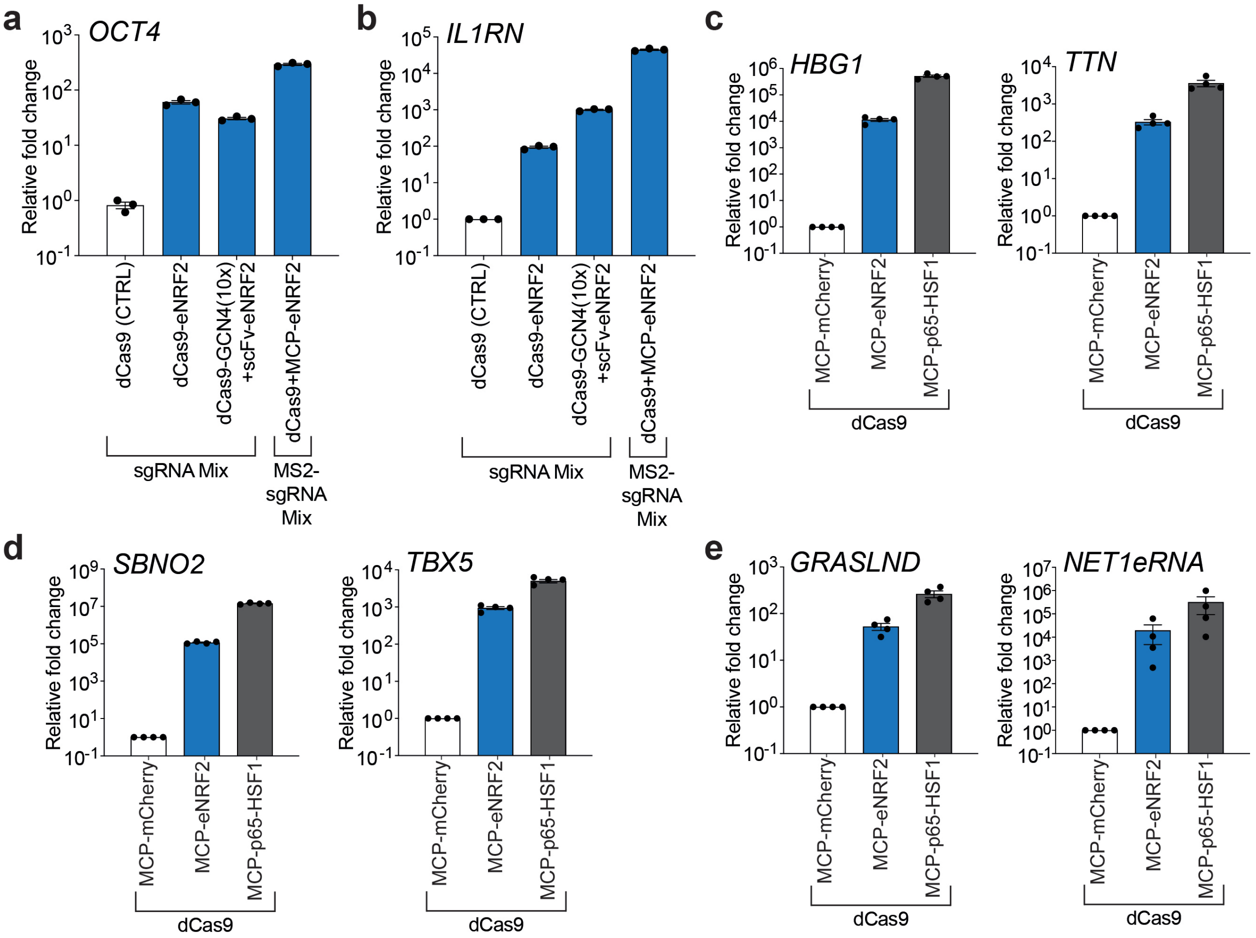


**Supplementary Fig. 4. Comparison and versatility of gene activation potential of the eNRF2 TAD. a and b.** *OCT4* and *IL1RN* mRNA levels after the eNRF2 TAD was recruited using the specified dCas9-based recruitment architecture (direct fusion, SunTag-based, or MCP-based) and 4 corresponding pooled gRNAs. **c and d.** mRNA levels for indicated loci after the eNRF2 TAD was recruited via dCas9 and targeted to promoters using pools of MS2 modified gRNAs (**panel c**; *HBG1* and *TTN* promoters) or a single gRNA (**panel d**; *SBNO2* and *TBX5* promoters). **e**. *GRASLND* long noncoding RNA (left) or *NET1* eRNA (right) levels after the eNRF2 TAD was recruited via dCas9 and targeted to each indicated locus using 4 and 2 pooled MS2 modified gRNAs, respectively. All samples were processed for QPCR analysis 72 hours post-transfection in HEK293T cells and are the result of at least 3 biological replicates. Error bars; SEM.

#### Supplementary Figure 5

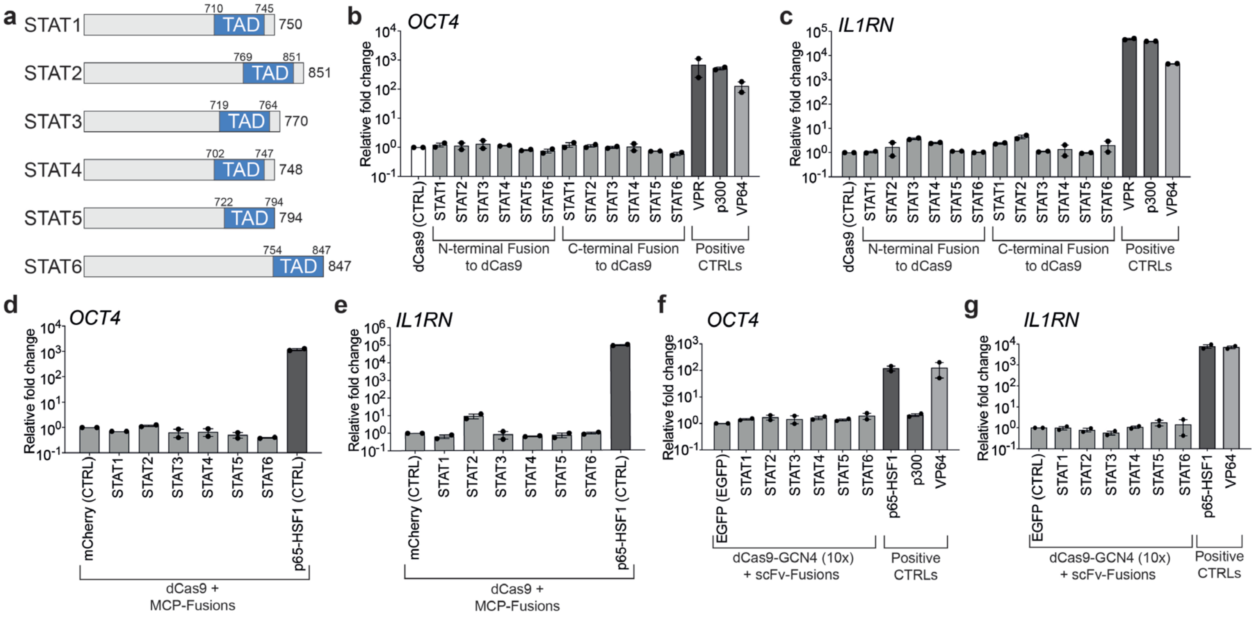


**Supplementary Fig. 5. Transactivation potency of cytokine responsive MTF TADs when recruited to human promoters via dCas9. a.** Schematics showing STAT family proteins STAT1, STAT2, STAT3, STAT4, STAT5 and STAT6. TADs for each respective protein are shown in light blue along with amino acid (aa) coordinates. **b and c.** *OCT4* and *IL1RN* mRNA levels after the indicated TADs were fused to the N- or C-terminus of dCas9 targeted to each respective promoter using 4 pooled standard gRNAs. **d and e.** *OCT4* and *IL1RN* mRNA levels after the indicated TADs were fused to the MCP protein and recruited via 4 pooled MS2 modified gRNAs and dCas9. MCP fused to the bipartite p65-HSF was used as a positive control. **f and g.** *OCT4* and *IL1RN* mRNA levels after the indicated TADs were fused to scFv and recruited via dCas9 harboring a 10xGCN4 C-terminal fusion protein (the SunTag system) along with 4 pooled standard gRNAs targeting each respective promoter. All samples were processed for QPCR analysis 72 hours post-transfection in HEK293T cells and are the result of at least 2 biological replicates. Error bars; SEM.

#### Supplementary Figure 6

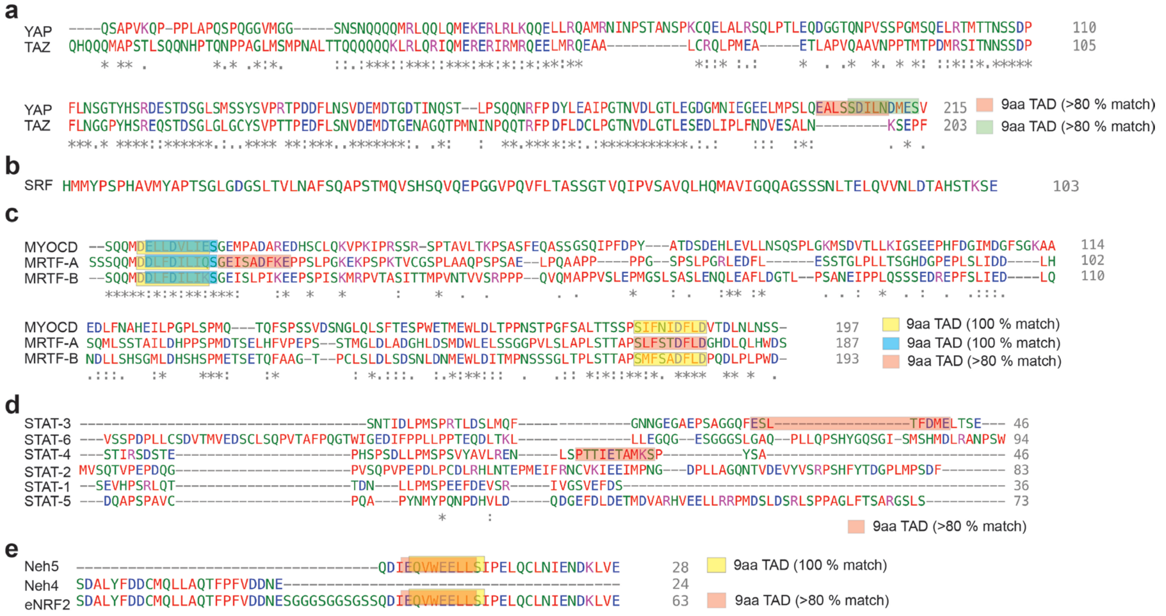


f

| TAD Domain | 9AA TAD (100% match) | 9AA TAD (>80% match) |
|---|---|---|
| YAP | 0 | 2 |
| TAZ | 0 | 0 |
| SRF | 0 | 0 |
| **MRTF-A** | **2** | **2** |
| **MRTF-B** | **3** | **0** |
| **MYOCD** | **3** | **0** |
| **Neh5** | **1** | **1** |
| Neh4 | 0 | 0 |
| **eNRF2** | **1** | **1** |
| STAT-1 | 0 | 0 |
| STAT-2 | 0 | 0 |
| STAT-3 | 0 | 1 |
| STAT-4 | 0 | 1 |
| STAT-5 | 0 | 0 |
| STAT-6 | 0 | 0 |

**Supplementary Fig. 6. Amino acid sequence analysis shows that potent mechanosensitive TAD domains contain single or multiple 9aa TAD segments with 100% matches to predictive software. a.** Clustal omega sequence alignment of TAD domains of two paralogous transcriptional activators YAP and TAZ. 9aa TADs with >80% match to algorithmic predictions (https://www.med.muni.cz/9aaTAD/) are either highlighted with light pink or light green. **b**. Amino acid sequence of the SRF TAD domain is shown. **c**. Clustal omega sequence alignment of TAD domains of the MYOCD, MRTF-A, and MRTF-B transcriptional activators are shown. 9aa TADs with 100% match to algorithmic predictions are either highlighted with light yellow or light blue and 9aa TADs with >80% to algorithmic predictions match are either highlighted with light pink or light green. **d**. Clustal omega sequence alignment of TAD domains from STAT-1 through STAT-6 are shown. 9aa TADs with >80% match to algorithmic predictions are highlighted with light pink. **e**. Amino acid sequence of Neh5, Neh4 and eNRF2 TADs are shown. 9aa TADs with 100% match to algorithmic predictions are highlighted with light yellow and 9aa TADs with >80% match to algorithmic predictions are highlighted with light pink. **f**. Table showing overall number of predicted 9aa TADs from all TAD domains used in this study.

#### Supplementary Figure 7

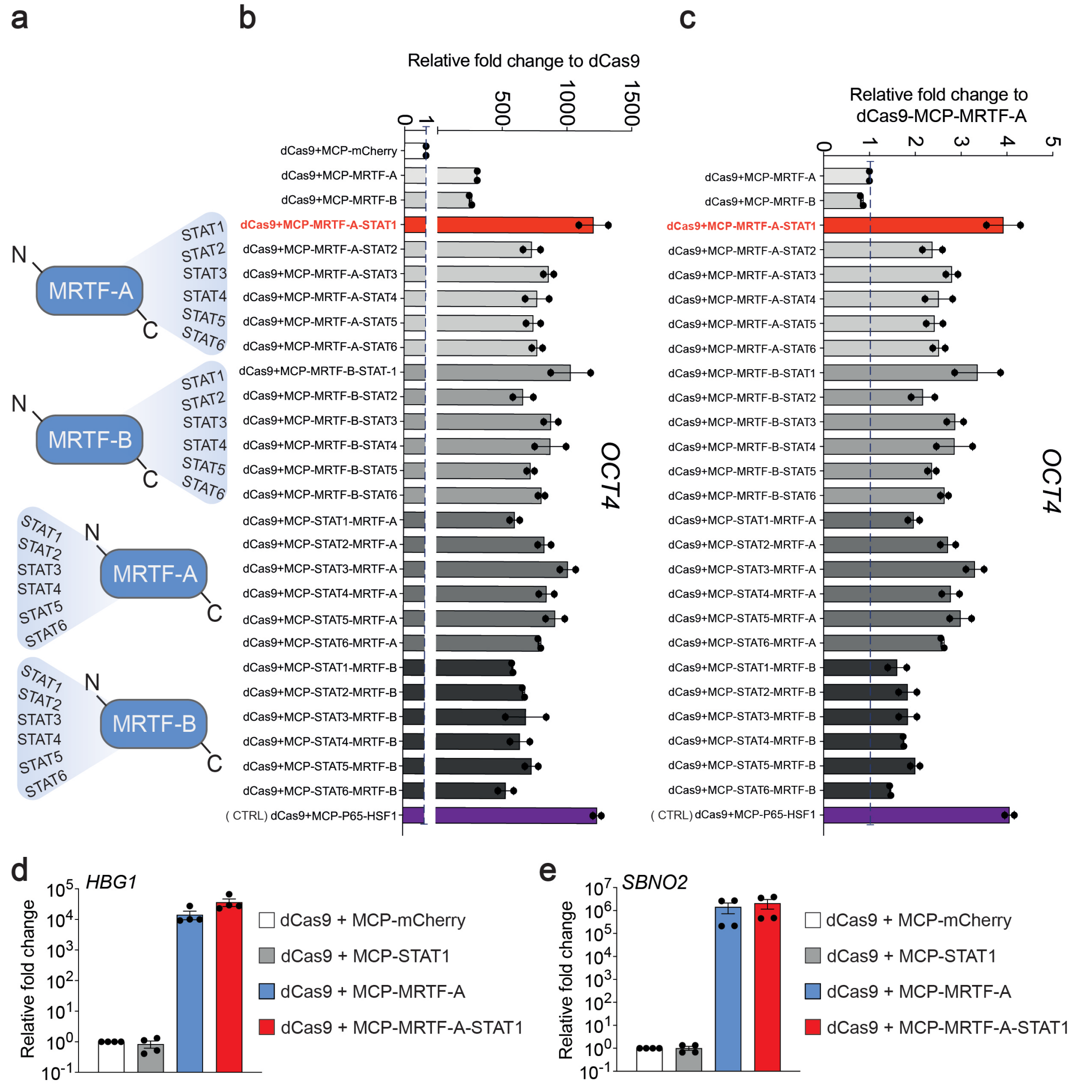


**Supplementary Fig. 7. Bipartite fusions between MRTF and STAT TADs enhance transactivation potential. a.** Schematic diagram showing the fusion of STAT1 through STAT6 TADs to the C- or N-terminus of MRTF-A or MRTF-B TADs. **b.** *OCT4* mRNA activation when targeted by indicated bipartite MRTF-STAT TAD fusions relative to dCas9 + MCP-mCherry. The dotted line indicates basal *OCT4* expression in dCas9 + MCP-mCherry transfected HEK293T cells. **c.** *OCT4* mRNA activation potential of indicated bipartite MRTF-STAT TAD fusions relative to dCas9 + MCP-MRTF-A. The dotted line indicates *OCT4* expression in dCas9 + MCP-MRTF-A transfected HEK293T cells. **d.** *HBG1* gene activation when dCas9 and indicated MCP fusions were targeted to the *HBG1* promoter using a pool of 4 MS2-modified gRNAs. **e.** *SBNO2* gene activation when dCas9 and indicated MCP fusions were targeted to the *SBNO2* promoter using a single MS2-modified gRNAs. All samples were processed for QPCR analysis 72 hours post-transfection and are the result of at least 2 biological replicates. Error bars; SEM.

#### Supplementary Figure 8

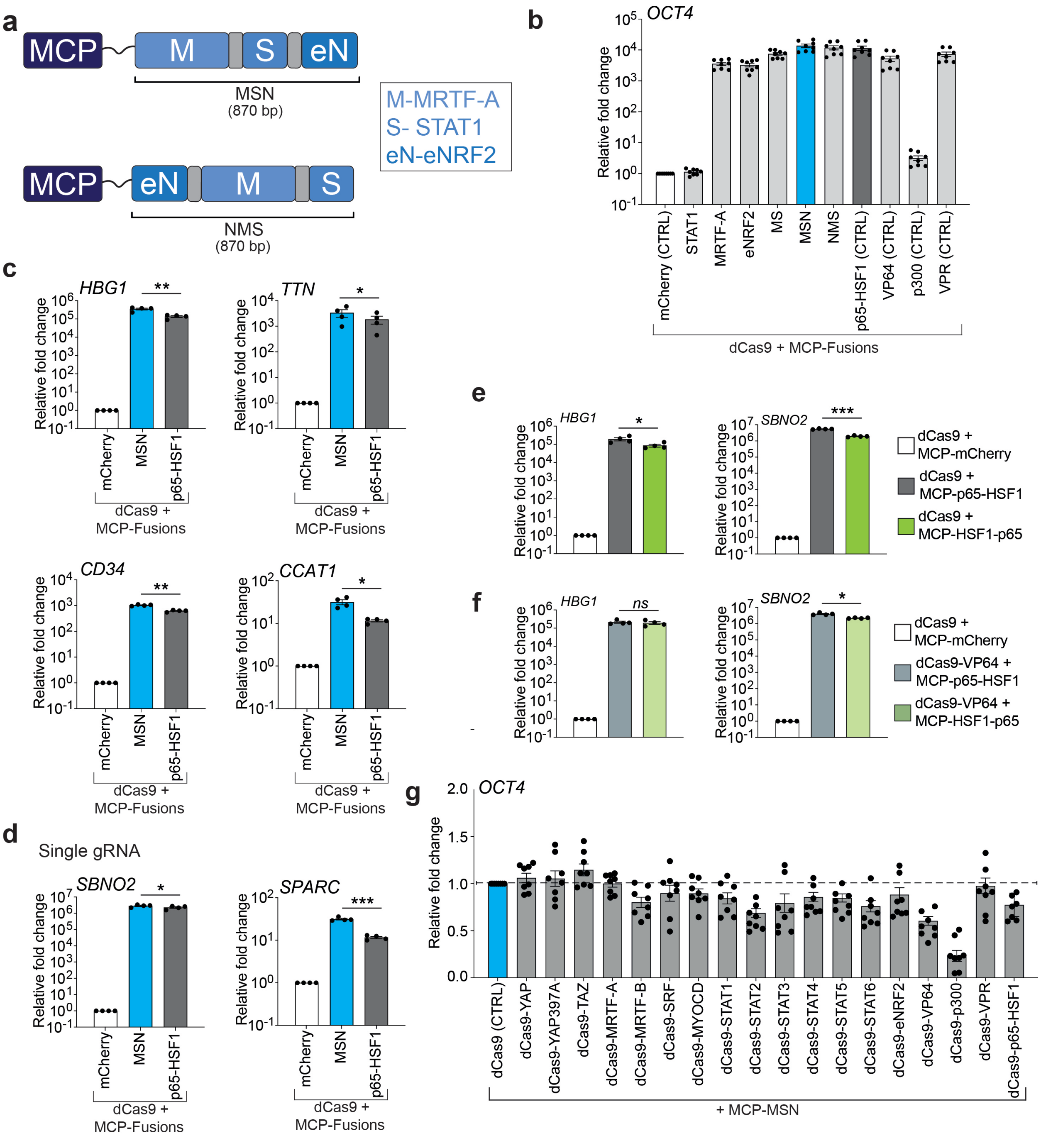


**Supplementary Fig. 8. Fusion of eNRF2 to the C- or N-terminus of MRTF-A-STAT1 (MS) fusions further enhances transactivation potency. a.** Schematic diagram showing tripartite MSN or NMS fusion proteins (base pair, bp sizes below) in conjunction with the MS2 binding protein; MCP. The eNRF2 TAD was either fused to the C-terminus or N-terminus of the bipartite MRTF-A-STAT1 TAD. **b.** *OCT4* mRNA levels after targeting with dCas9, 4 MS2-modified *OCT4* gRNAs, and the indicated MS2-recruited tripartite

TADs. **c.** Relative expression of *HBG1***,** *TTN***,** *CD34*, or *CCAT1* after dCas9 + MCP-mCherry control, dCas9 + MCP-MSN, or dCas9 + MCP-p65-HSF1 systems were targeted to their respective promoters using pools of gRNAs in HEK293T cells. **d.** Relative expression of *SBNO2* or *SPARC* after dCas9 + MCP-mCherry control, dCas9 + MCP-MSN, or dCas9 + MCP-p65-HSF1 systems were targeted to their respective promoters using a single gRNA in HEK293T cells. **e.** Relative expression of *HBG1* or *SBNO2* mRNA levels after dCas9 + MCP-mCherry control, dCas9 + MCP-p65-HSF1, or dCas9 + MCP-HSF1-p65 systems were targeted to their respective promoters using pools of gRNAs or a single gRNA, respectively in HEK293T cells. **f.** Relative expression of *HBG1* or *SBNO2* mRNA levels after dCas9 + MCP-mCherry control, dCas9-VP64 + MCP-p65-HSF1, or dCas9-VP64 + MCP-HSF1-p65 systems were targeted to their respective promoters using pools of gRNAs or a single gRNA, respectively in HEK293T cells. **g.** *OCT4* mRNA levels after targeting with dCas9 or indicated direct dCas9 C-terminal TAD fusions, 4 MS2-modified gRNAs, and the MS2-recruited tripartite MCP-MSN TAD. *OCT4* activation levels are presented relative to the activation achieved by dCas9+MCP-MSN. All samples were processed for QPCR analysis 72 hours post-transfection and are the result of at least 3 biological replicates. See source data for more information. Error bars; SEM. *; $P < 0.05$, **; $P < 0.01$, ***; $P < 0.001$. *ns*; not significant.

#### Supplementary Figure 9

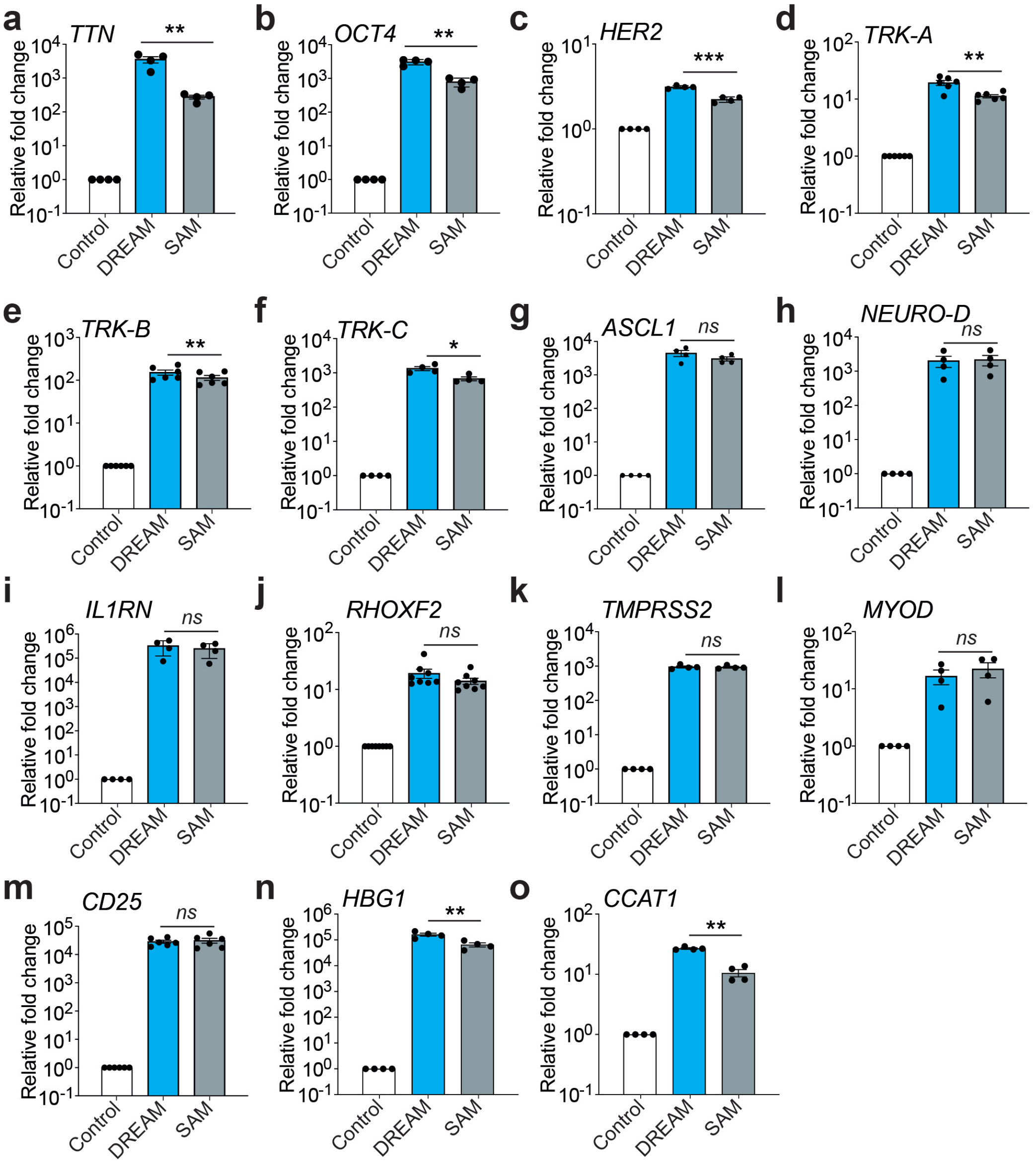


**Supplementary Fig. 9. CRISPR-DREAM mediated activation of coding genes in HEK293T cells using pooled gRNAs. a – m.** Relative expression levels of 13 different endogenous human genes after Control, DREAM, or SAM systems were targeted to their respective promoters using pools of gRNAs in HEK293T cells. **n and o.** Relative expression of *HBG1* or *CCAT1* after equimolar amounts of control (187.5ng dCas9 + 187.05ng MCP-mCherry), DREAM (187.5ng dCas9 + 187.5ng MCP-MSN), or SAM (189.77ng dCas9-VP64 + 188.64ng MCP-p65-HSF1) systems were targeted to their respective promoters using pools of gRNAs, respectively in HEK293T cells. All samples were processed for QPCR analysis 72 hours post-transfection and are the result of at least 3 biological replicates. Error bars; SEM. *; $P < 0.05$, **; $P < 0.01$, ***; $P < 0.001$. *ns*; not significant.

#### Supplementary Figure 10

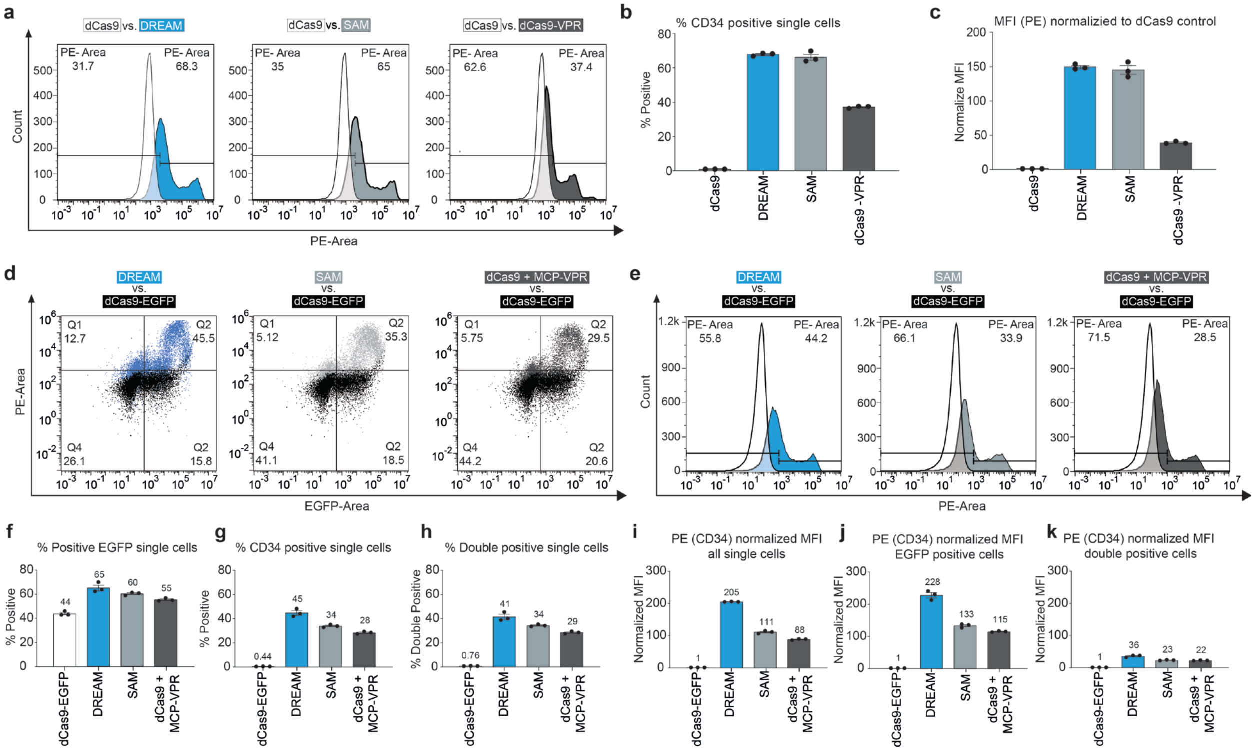


**Supplementary Fig. 10. Comparisons of CRISPR-DREAM, SAM, and dCas9-VPR induced CD34 expression in single HEK293T cells. a.** Histograms illustrating CD34-PE fluorescence after transfection of 4 *CD34*-targeting gRNAs and either DREAM (left in blue; gRNAs are MS2-modified), SAM (middle in light gray; gRNAs are MS2-modfied), dCas9-VPR (right in dark gray) or dCas9 (control in white). **b and c.** CD34 positivity and normalized mean fluorescence intensity (MFI) compared to dCas9 control, respectively following transfection of dCas9 (control), DREAM (dCas9 + MCP-MSN-STOP), SAM (dCas9 + MCP-p65-HSF1-STOP), or dCas9-VPR as in **panel a**. **d.** Histograms showing CD34 (via PE) and MCP-fusion (via EGFP) signal intensities after transfection of 4 MS2-modified *CD34*-targeting gRNAs and either DREAM (left in blue), SAM (middle in light gray), or dCas9 + MCP-VPR (right in dark gray) compared with dCas9-EGFP control (shown in black). **e.** Histograms illustrating CD34-PE fluorescence after transfection of 4 MS2-modified *CD34*-targeting gRNAs and either DREAM (left in blue), SAM (middle in light gray), dCas9 + MCP-VPR (right in dark gray) or dCas9-EGFP (control in white). **f and g.** Percent EGFP positive (**panel f**) and percent CD34 positive (**panel g**) single cell populations following transfection of 4 MS2-modified *CD34*-targeting gRNAs and either dCas9-EGFP (control), DREAM, SAM, or dCas9 + MCP-VPR. **h.** Percent CD34 and EGFP double positive HEK293T cells following transfection of 4 MS2-modified *CD34*-targeting gRNAs and either dCas9-EGFP (control), DREAM, SAM, or dCas9 + MCP-VPR. **i – k.** CD34 (via PE) MFI in single (**panel i**), EGFP positive (**panel j**), and double positive (**panel k**) cell populations following transfection of 4 MS2-modified *CD34*-targeting gRNAs dCas9-EGFP (control), DREAM, SAM, or dCas9 + MCP-VPR. All experiments were performed in triplicate and all histograms are representative of all data. See source data for more information.

**Supplementary Figure 11**

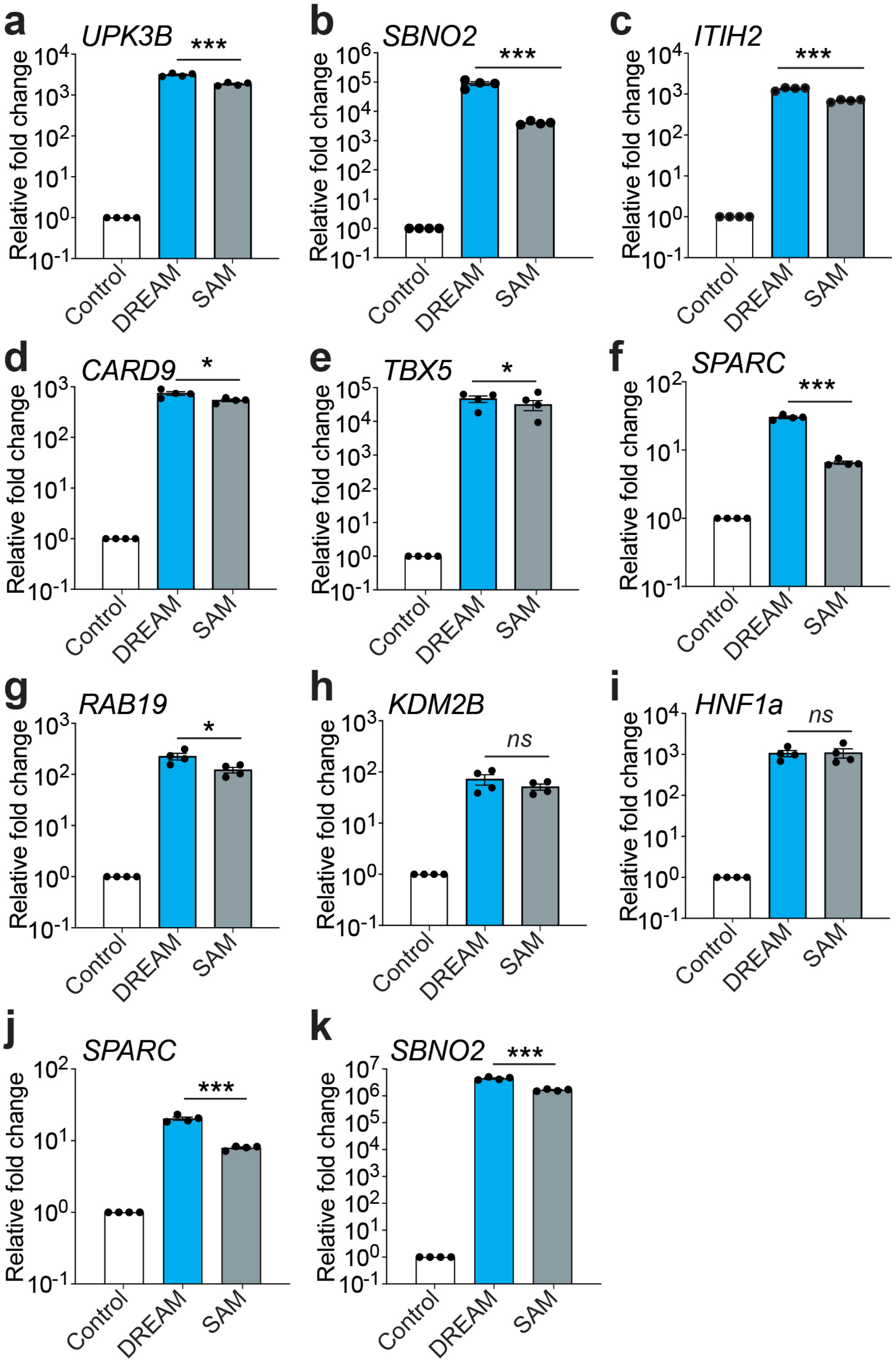


**Supplementary Fig. 11. CRISPR-DREAM mediated activation of coding genes in HEK293T cells using single gRNAs. a – i.** Relative expression levels of 9 different endogenous human genes 72 hours after Control, DREAM, or SAM systems were targeted to their respective promoters using a single gRNA in HEK293T cells**. j and k.** Relative expression of *SPARC* or *SBNO2* after equimolar amounts of control (187.5ng dCas9 + 187.05ng MCP-mCherry), DREAM (187.5ng dCas9 + 187.5ng MCP-MSN), or SAM (189.77ng dCas9-VP64 + 188.64ng MCP-p65-HSF1) systems were targeted to their respective promoters using a single gRNA, respectively in HEK293T cells. All samples were processed for QPCR analysis 72 hours post-transfection and are the result of at least 3 biological replicates. See source data for more information. Error bars; SEM. *; $P < 0.05$, ***; $P < 0.001$. *ns*; not significant.

#### Supplementary Figure 12

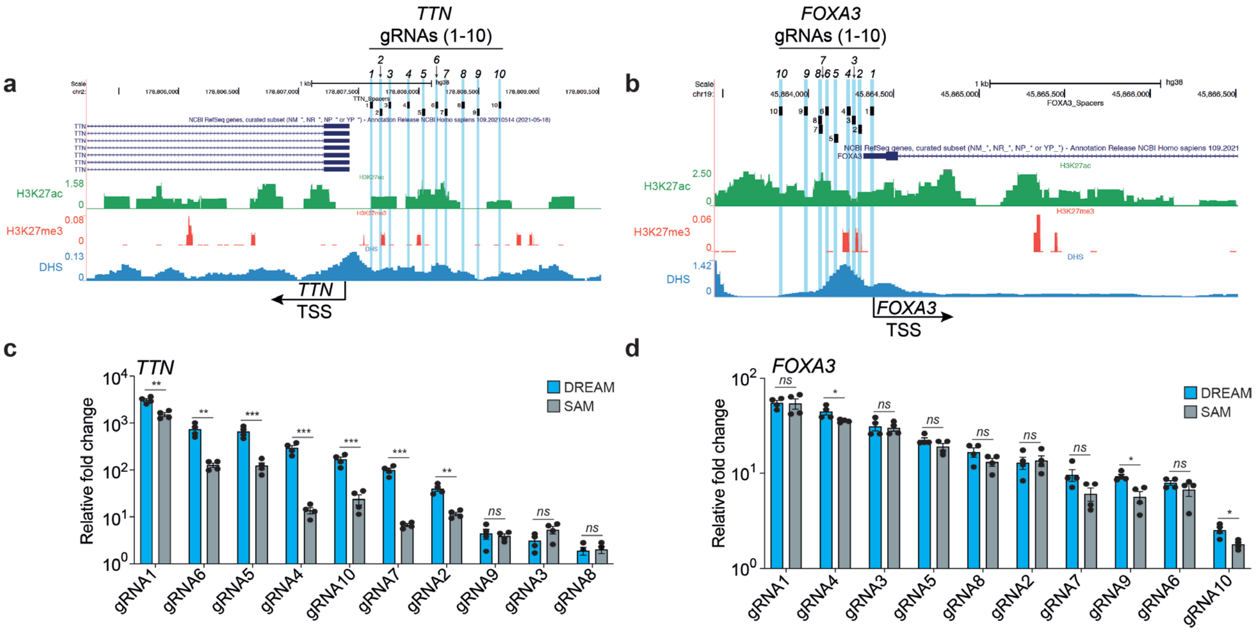


**Supplementary Fig. 12. Comparison of gene activation potency between DREAM and SAM systems spanning ~1kb regions upstream from human TSSs. a and b.** The genomic regions (hg38) encompassing the human *TTN* (**panel a**) and *FOXA3* (**panel b**) genes on chromosomes 2 and 19, respectively, are shown. Genes and isoforms are shown in dark blue; gRNA target regions are indicated by black lines, numbers, and light blue highlighting. H3K27ac (from GSE174866), H3K27me3 (from DRX013192), and DNase Hypersensitivity Sites (DHSs; from GSE32970) are shown in green, red, and blue, respectively. Transcription Start Sites (TSSs) for each gene indicated by black arrows. **c and d**. Comparison of transactivation potency between DREAM and SAM systems when targeted to indicated sites within the *TTN* or *FOXA3* promoters, respectively. 10 gRNAs were designed to tile across ~1kb upstream of respective promoter regions. Non-transfected HEK293T cells were used as control. All samples were processed for QPCR analysis 72 hours post-transfection and are the result of at least 4 biological replicates. Error bars; SEM. *; $P < 0.05$, **; $P < 0.01$, ***; $P < 0.001$. *ns*; not significant.

#### Supplementary Figure 13

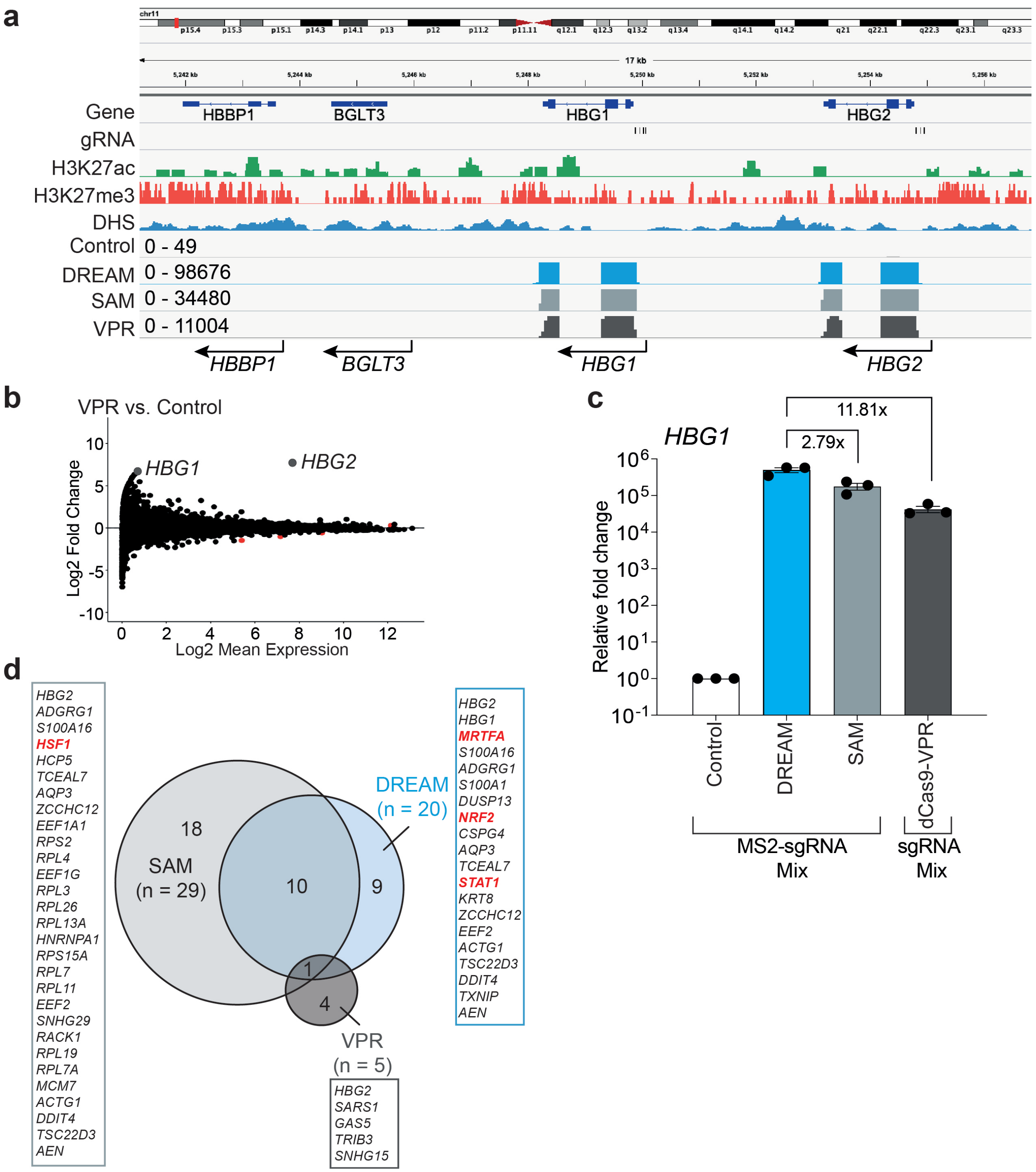


**Supplementary Fig. 13. CRISPR-DREAM mediated activation of *HBG1/HBG2* is specific, robust, and potent. a.** The genomic region encompassing the human *HBG1* and *HBG2* genes, along with two nearby genes *BGLT3* and *HBBP1* on chromosome 11 (hg38) is shown. Genes are shown in dark blue; gRNA target regions are indicated by black lines. H3K27ac (from GSE174866), H3K27me3 (from DRX013192), and DNase

Hypersensitivity Sites (DHSs; from GSE32970) are shown in green, red, and blue, respectively. **b.** An MA plot generated from DESeq2 analysis 72 hours after HEK293T cells were transiently co-transfected with dCas9-VPR and four *HBG1/HBG2* promoter targeting gRNAs. mRNAs corresponding to *HBG1* and *HBG2* isoforms (statistically significant differentially expressed with a fold change (FC) > 2 or < -2 and a false discovery rate (FDR) < 0.05) are shown in deep gray. Red dots indicate other statistically significant differentially expressed genes (FC > 2 or < -2, and FDR < 0.05). **c.** QPCR analysis showing *HBG1* expression after targeting with DREAM, SAM, or dCas9-VPR in HEK293T cells. dCas9 + MCP-mCherry was used as a control. All samples were processed for QPCR analysis 72 hours post-transfection in HEK293T cells and are the result of at least 3 biological replicates. Error bars; SEM. **d.** Venn diagram showing all statistically significant differentially regulated genes (FC > 2 or < -2, and FDR< 0.05) in HEK293T cells after the *HBG1/HBG2* promoters were targeted as above using the DREAM, SAM, or dCas9-VPR systems. Genes in bold red font are components of human TADs used in respective DREAM or SAM systems.

### Supplementary Figure 14

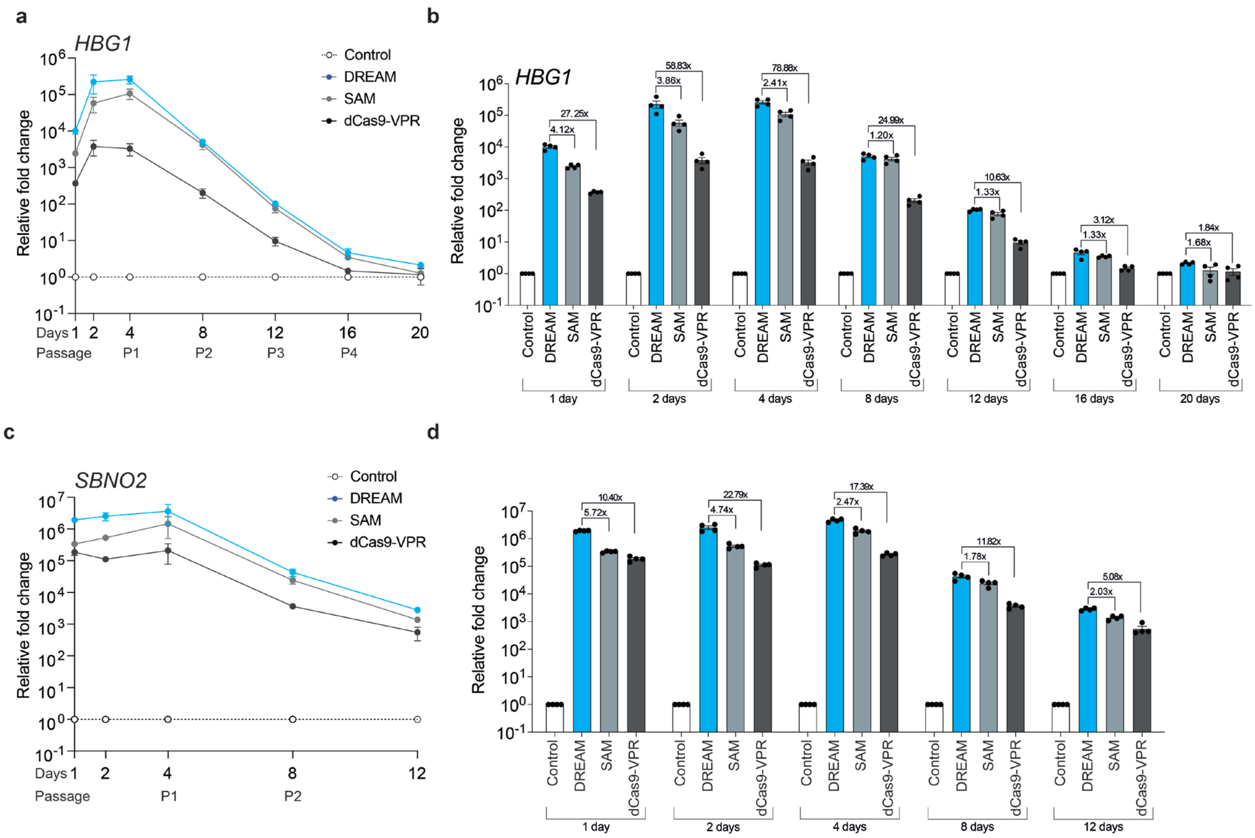


**Supplementary Fig. 14. Comparisons of functional durability between the DREAM, SAM, and dCas9-VPR systems in HEK293T cells. a and b.** *HBG1* mRNA expression is shown over 20 days post-transfection of 4 *HBG1* gRNAs and either DREAM (MS2-modified gRNAs), SAM (MS2-modified gRNAs), or dCas9-VPR compared with the dCas9 + MCP-mCherry control (MS2-modified gRNAs). ~15% of cells were passaged into a new 24 well plate at days 4, 8, 12, and 16 post-transfection. **c and d.** *SBNO2* mRNA expression is shown over 12 days post-transfection of 1 *SBNO2* gRNA and either DREAM (MS2-modified gRNA), SAM (MS2-modified gRNA), or dCas9-VPR compared with the dCas9 + MCP-mCherry control (MS2-modified gRNA). ~15% of cells were passaged into a new 24 well plate at days 4 and 8 post-transfection. All data are the result of at least 4 biological replicates.

#### Supplementary Figure 15

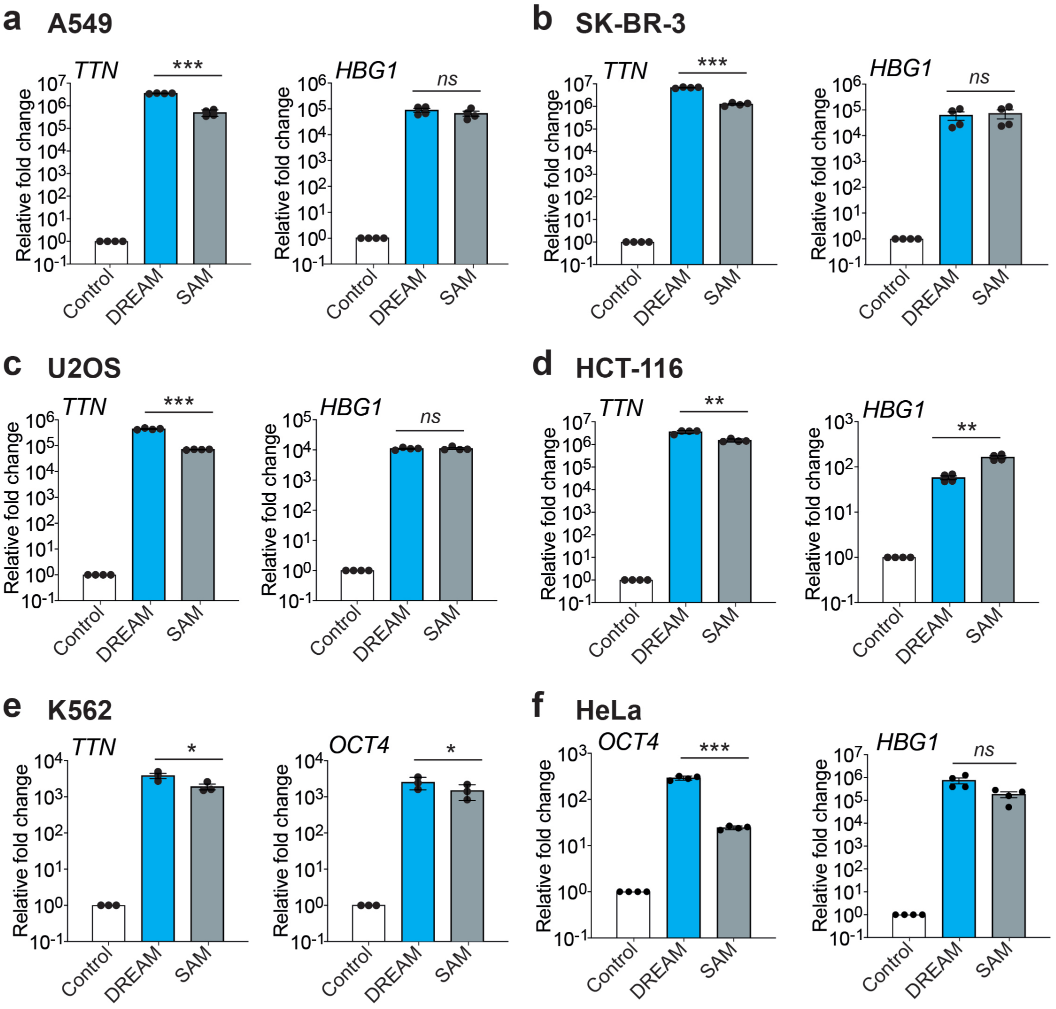


**Supplementary Fig. 15. CRISPR-DREAM is robust across diverse human cancer cell lines. a – f.** Transactivation potencies of DREAM and SAM systems are shown when targeted to indicated human promoters in a lung adenocarcinoma cell line (A549; **panel a**), a breast cancer cell line (SK-BR-3; **panel b**), a bone osteosarcoma epithelial cell line (U2OS **panel c**), a colorectal carcinoma cell line (HCT-116; **panel d**), a myelogenous leukemia cell line (K562; **panel e**), and a cervical cancer cell line (HeLa; **panel f**). All samples were processed for QPCR analysis 72 hours post-transfection and are the result of at least 3 biological replicates. See source data for more information. Error bars; SEM. *; $P < 0.05$, **; $P < 0.01$, ***; $P < 0.001$. *ns*; not significant.

**Supplementary Figure 16**

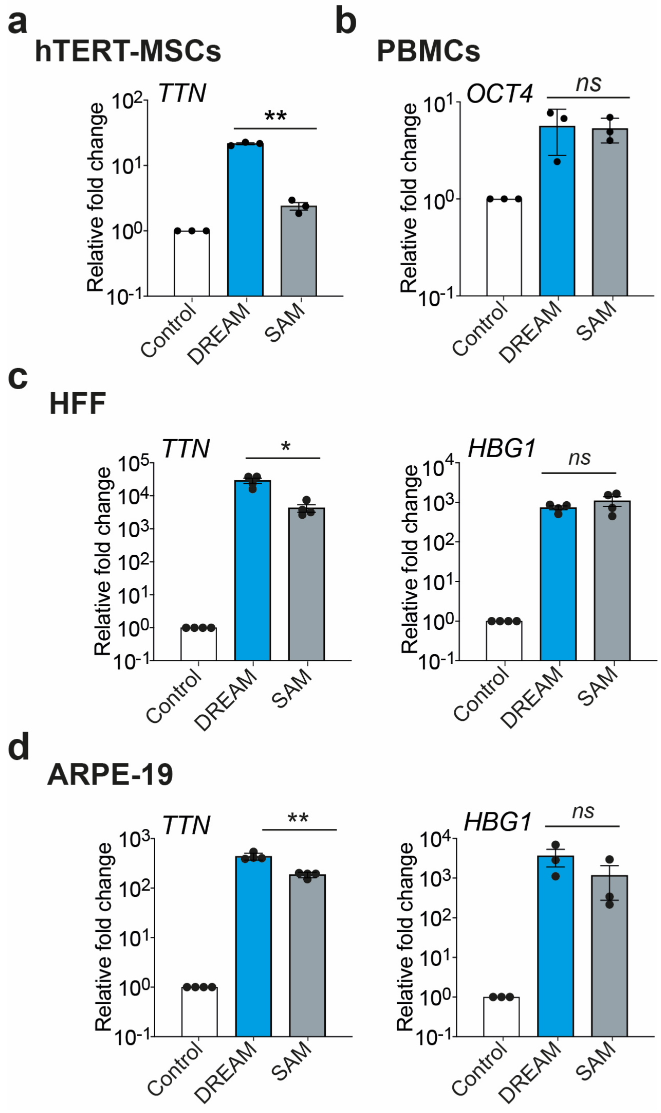


**Supplementary Fig. 16. CRISPR-DREAM is robust across diverse karyotypically normal human cells. a. a – c.** Transactivation potencies of DREAM and SAM systems are shown when targeted to indicated human promoters in hTERT-MSCs (**panel a**), PBMCs (**panel b**), Human Foreskin Fibroblasts (HFF; **panel c**), or retinal pigmented epithelial cells (ARPE-19; **panel d**). All samples were processed for QPCR analysis 72 hours post-transfection and are the result of at least 3 biological replicates. See source data for more information. Error bars; SEM. *; $P < 0.05$, **; $P < 0.01$. *ns*; not significant.

**Supplementary Figure 17**

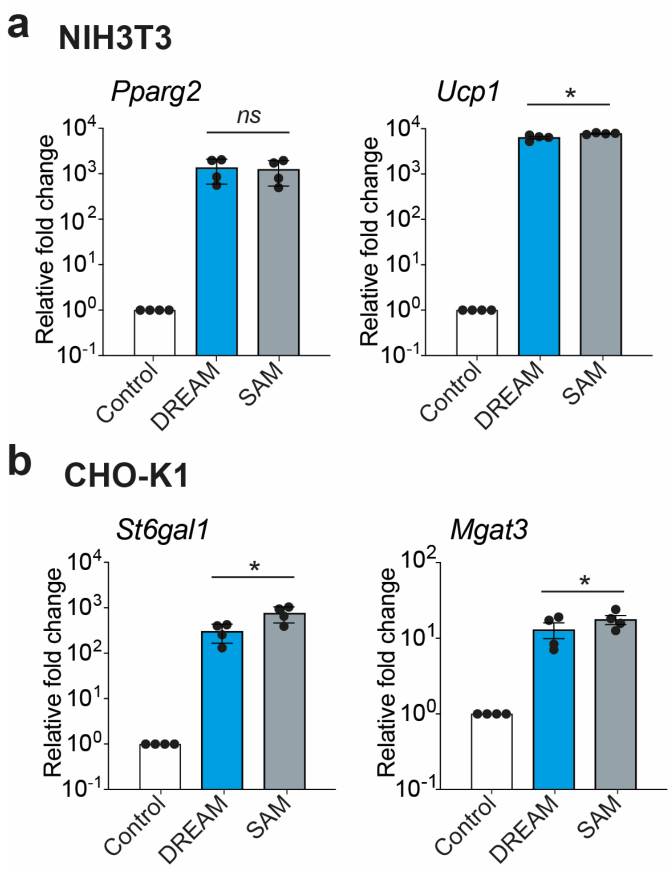


**Supplementary Fig. 17. CRISPR-DREAM is robust in rodent cells. a and b.** Transactivation potencies of DREAM and SAM systems are shown when targeted to indicated human promoters in murine NIH3T3 cells (**panel a**) or Chinese hamster ovary (CHO-K1) cells (**panel b**). All samples were processed for QPCR analysis 72 hours post-transfection and are the result of at least 3 biological replicates. See source data for more information. Error bars; SEM. *; $P < 0.05$. *ns*; not significant.

#### Supplementary Figure 18

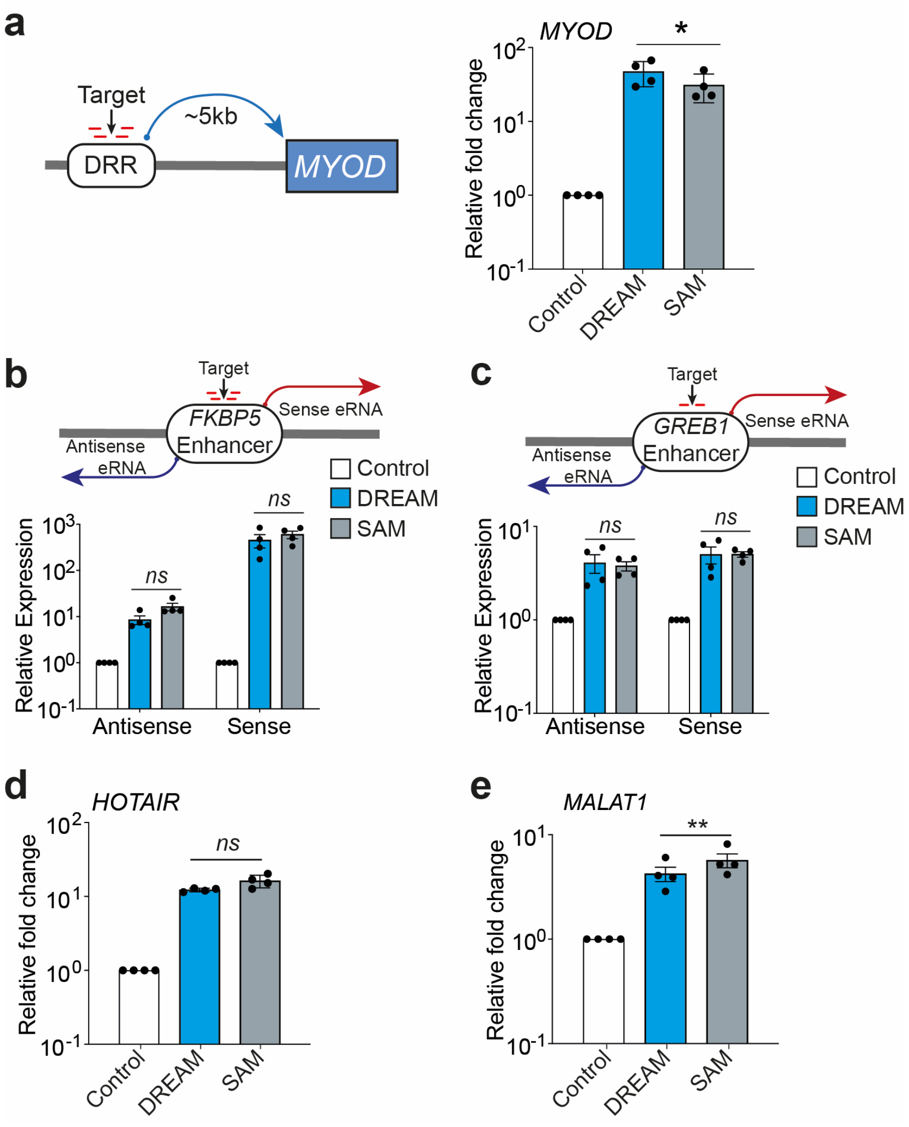


**Supplementary Fig. 18. CRISPR-DREAM activates gene expression when targeted to enhancers, activates eRNAs, and activates lncRNAs in human cells. a.** *MYOD* mRNA levels after DREAM or SAM systems were targeted to the *MYOD* distal regulatory region (DRR) using 4 MS2-modified gRNAs. **b and c**. eRNA levels induced after DREAM or SAM systems were targeted to the *FKBP5* (**panel b**) or *GREB1* (**panel c**) enhancer in HEK293T cells. **d and e**. lncRNA levels induced after DREAM or SAM systems were targeted to the *HOTAIR* (**panel d**) or *MALAT1* (**panel e**) lncRNA promoters in HEK293T cells. All samples were processed for QPCR analysis 72 hours post-transfection and are the result of at least 4 biological replicates. See source data for more information. Error bars; SEM. *; $P < 0.05$, **; $P < 0.01$. *ns*; not significant.

#### Supplementary Figure 19

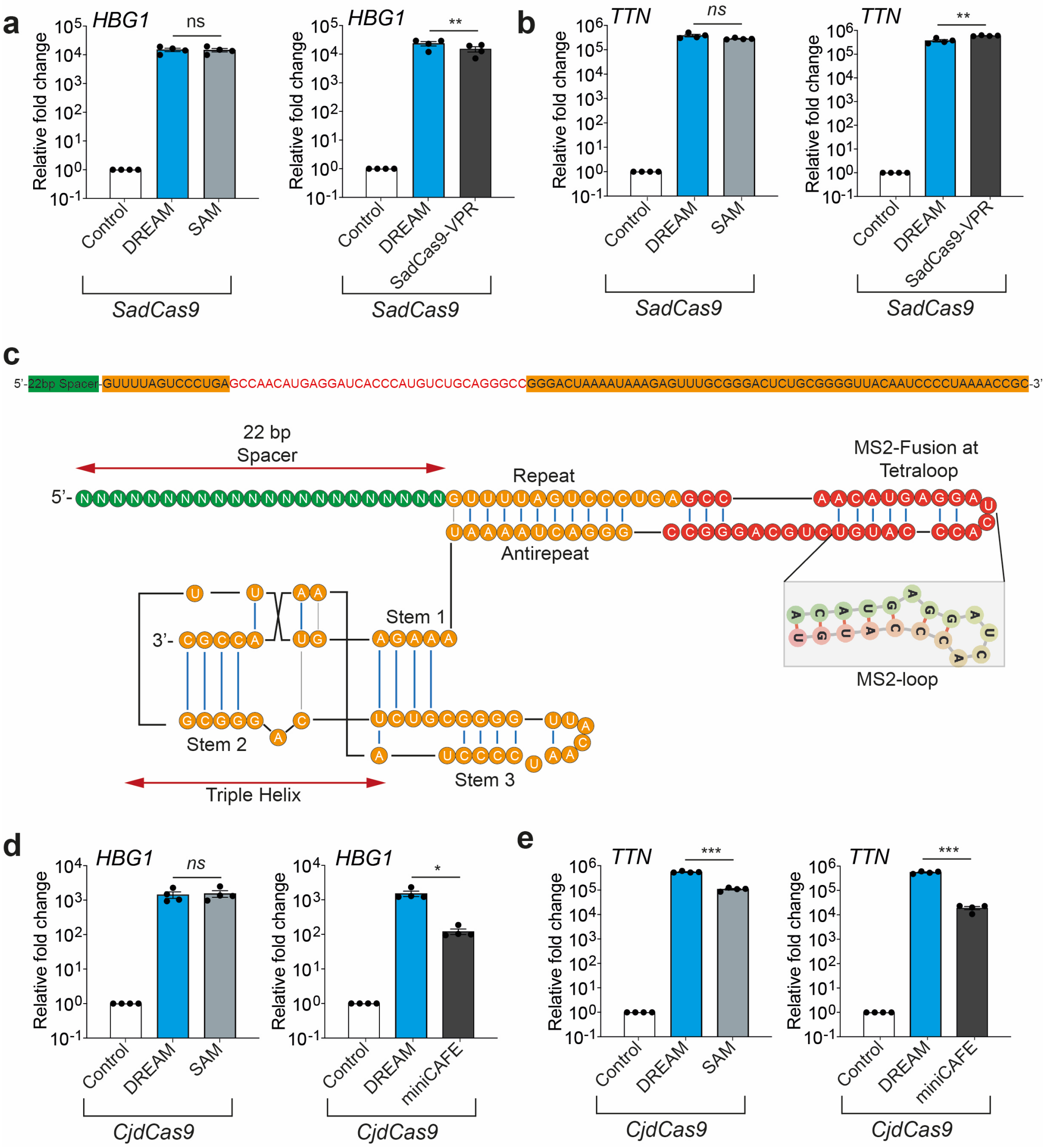


**Supplementary Fig. 19. Orthogonal CRISPR-DREAM systems are potent in HeLa cells. a and b.** SadCas9-DREAM mediated transactivation when targeted to the promoters of 2 different endogenous genes (*HBG1*; **panel a**, and *TTN;* **panel b**) in comparison to SadCas9-SAM or SadCas9-VPR systems. **c.** RNA sequence of the MS2 loop containing gRNA designed for CjdCas9 and schematics of CjCas9 gRNA with MS2 loop (magnified in inset) incorporated within the tetraloop region. **d and e.** CjdCas9-DREAM mediated transactivation when targeted to the promoters of 2 different endogenous genes (*HBG1*; **panel d**, and *TTN;* **panel e**) in comparison to CjdCas9-SAM or MiniCAFE systems. All samples were process for QPCR analysis 72 hours post-transfection and are the result of at least 4 biological replicates. See source data for more information. Error bars; SEM. *; $P < 0.05$, **; $P < 0.01$, ***; $P < 0.001$. *ns*; not significant.

#### Supplementary Figure 20

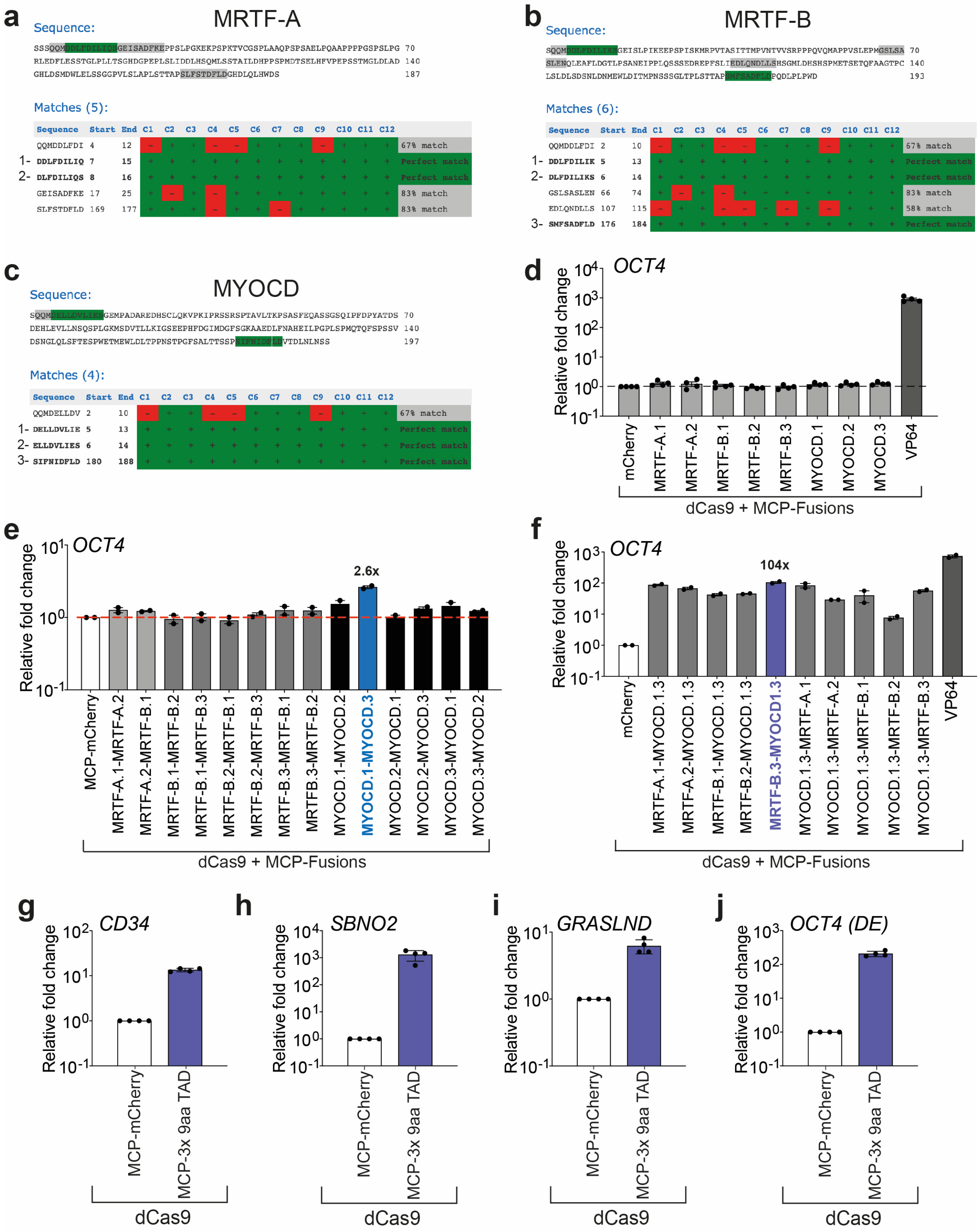

**Supplementary Fig. 20. Prediction, construction, and validation of transactivation potential among different 9aa TADs. a – c.** different 9aa TADs from MRTF-A (**panel a**), MRTF-B (**panel b**) or MYOCD (**panel c**) were predicted using the Nine Amino Acids

Transactivation Domain 9aaTAD Prediction Tool (https://www.med.muni.cz/9aaTAD/). We selected 9aa TADs that showed 100% matches to database predictions. **d.** *OCT4* gene activation when indicated 9aa TADs were fused to MCP and then recruited to *OCT4* promoter using dCas9 and a pool of 4 MS2-modified gRNAs. **e.** *OCT4* gene activation when indicated bipartite TADs, built using heterotypic 1x 9aa TADs, were fused to MCP and recruited to the *OCT4* promoter using dCas9 and a pool of 4 MS2-modified gRNAs. The blue bar (MCP-MYOCD.1-MYOCD.3) showed the highest gene activation among all 2x 9aa TADs and was selected for generating heterotypic 3x 9aa TADs. **f.** *OCT4* gene activation when indicated tripartite TADs, built using heterotypic 2x 9aa TADs, were fused to MCP and recruited to the *OCT4* promoter using dCas9 and a pool of 4 MS2-modified gRNAs. The purple bar (MCP-MRTF-B.3-MYOCD.1-MYOCD.3) showed the highest gene activation among all 3x 9aa TADs and was selected for further analysis. **g – h.** Relative gene activation when the selected 3x 9aa TAD (MCP-MRTF-B.3-MYOCD.1-MYOCD.3) was fused to MCP and recruited to either the *CD34* (**panel g**) or *SBNO2* (**panel h**) promoter using dCas9 and a pool of 4 MS2-modified gRNAs or a single MS2-modified gRNA, respectively. **i-j.** Levels of *GRASLND* lncRNA (**panel i**) or *OCT4* mRNA (**panel j**) after the selected 3x 9aa was recruited via dCas9 and targeted to either the *GRASLND* promoter or OCT distal enhancer (*OCT4*-DE), respectively, using pools of 4 MS2-modifed gRNAs. All samples were processed for QPCR analysis 72 hours post-transfection in HEK293T cells and are the result of at least 2 biological replicates. Error bars; SEM.

**Supplementary Figure 21**

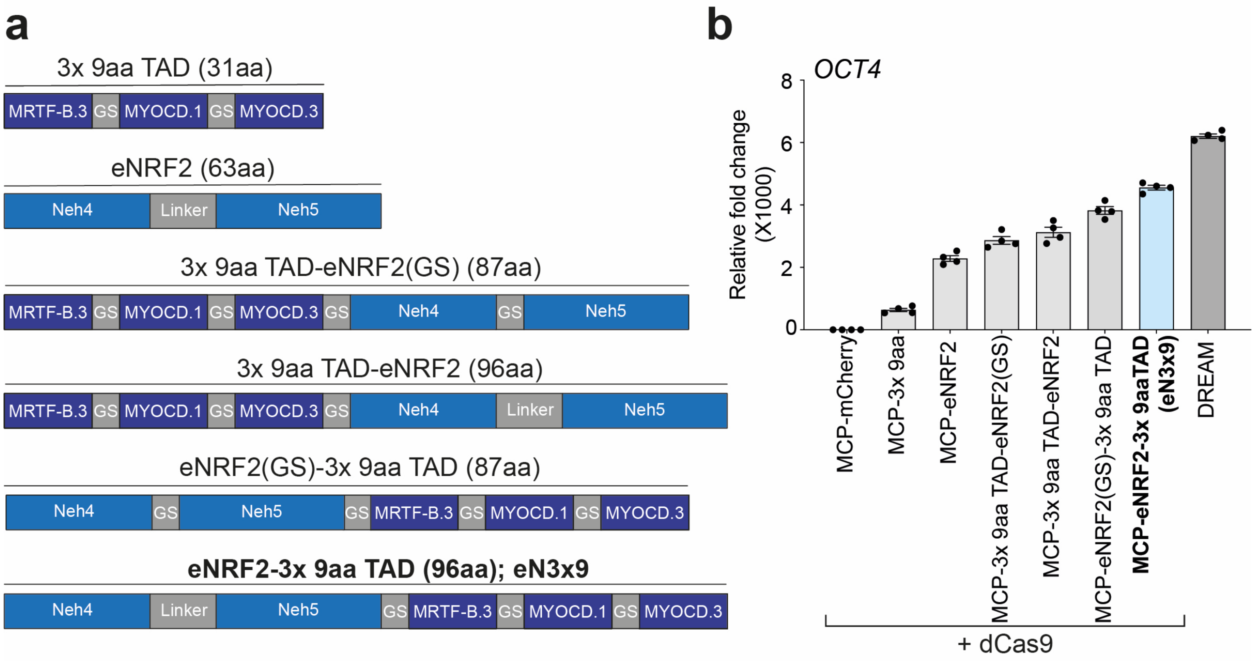


**Supplementary Fig. 21. Construction and validation of the eN3x9 TAD domain for the mini-DREAM system. a.** The 3x 9aa TADs derived from MRTF-B or MYOCD are depicted schematically. eNRF2 and different combinations of fusion architectures between indicated 3x 9aa TADs with eNRF2 fusions are also shown. 3x 9aa TADs were either cloned to the C- or N-terminal of eNRF2 and separated by either single glycine-serine linker (GS) or a 11 amino acid extended glycine-serine linker (Linker; see **Supplementary Fig. 3a**). Respective aa sizes of each fusion proteins are also shown. **b.** transcriptional activation of *OCT4* after 3x 9aa TAD, eNRF2, or indicated TAD fusions were recruited to the *OCT4* promoter via dCas9 and 4 pooled of MS2 modified gRNAs. The CRISPR-DREAM system is shown for comparison. All samples were processed for QPCR analysis 72 hours post-transfection in HEK293T cells and are the result of at least 4 biological replicates. Error bars; SEM.

#### Supplementary Figure 22

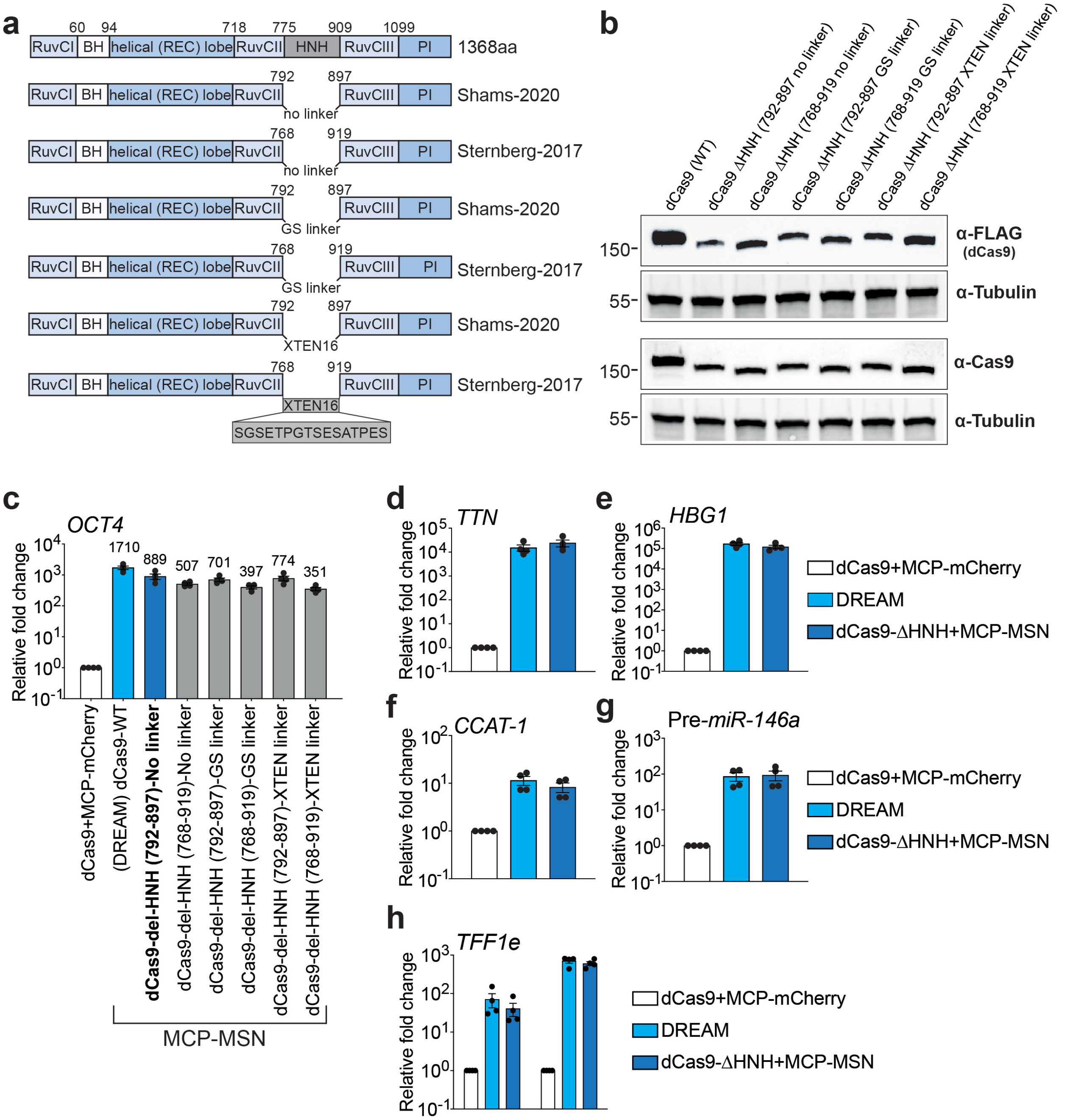


**Supplementary Fig. 22. Design, construction, and validation of HNH domain-deleted dCas9 variants for the mini-DREAM system. a.** Different HNH domain-deleted SpdCas9 variants, along with wildtype dCas9, are schematically depicted. Two different HNH domain-deleted SpCas9 variants (amino acid, aa 792-897, or aa 768-919 deleted, respectively) were selected for analysis and reconstructed using either no linker, a single glycine-serine linker, or an XTEN16 linker separating dCas9 protein segments. **b.** All HHN domain-deleted dCas9 variants were expressed in HEK293T cells and Western blotting was performed 72 hours post-transfection in HEK293T cells using either anti-FLAG or anti-Cas9 antibodies (Tubulin was used as a loading control). **c.** *OCT4* transactivation when MCP-MSN was recruited to the *OCT4* promoter using different indicated HNH domain-deleted dCas9 variants and a pool of 4 MS2-modified gRNAs. **d – h.** Comparison of transactivation potential between CRISPR-DREAM and selected HNH domain-deleted dCas9 (with no linker) + MCP-MSN at indicated endogenous loci. All samples were processed for QPCR analysis 72 hours post-transfection in HEK293T cells and are the result of at least 4 biological replicates. Error bars; SEM.

#### Supplementary Figure 23

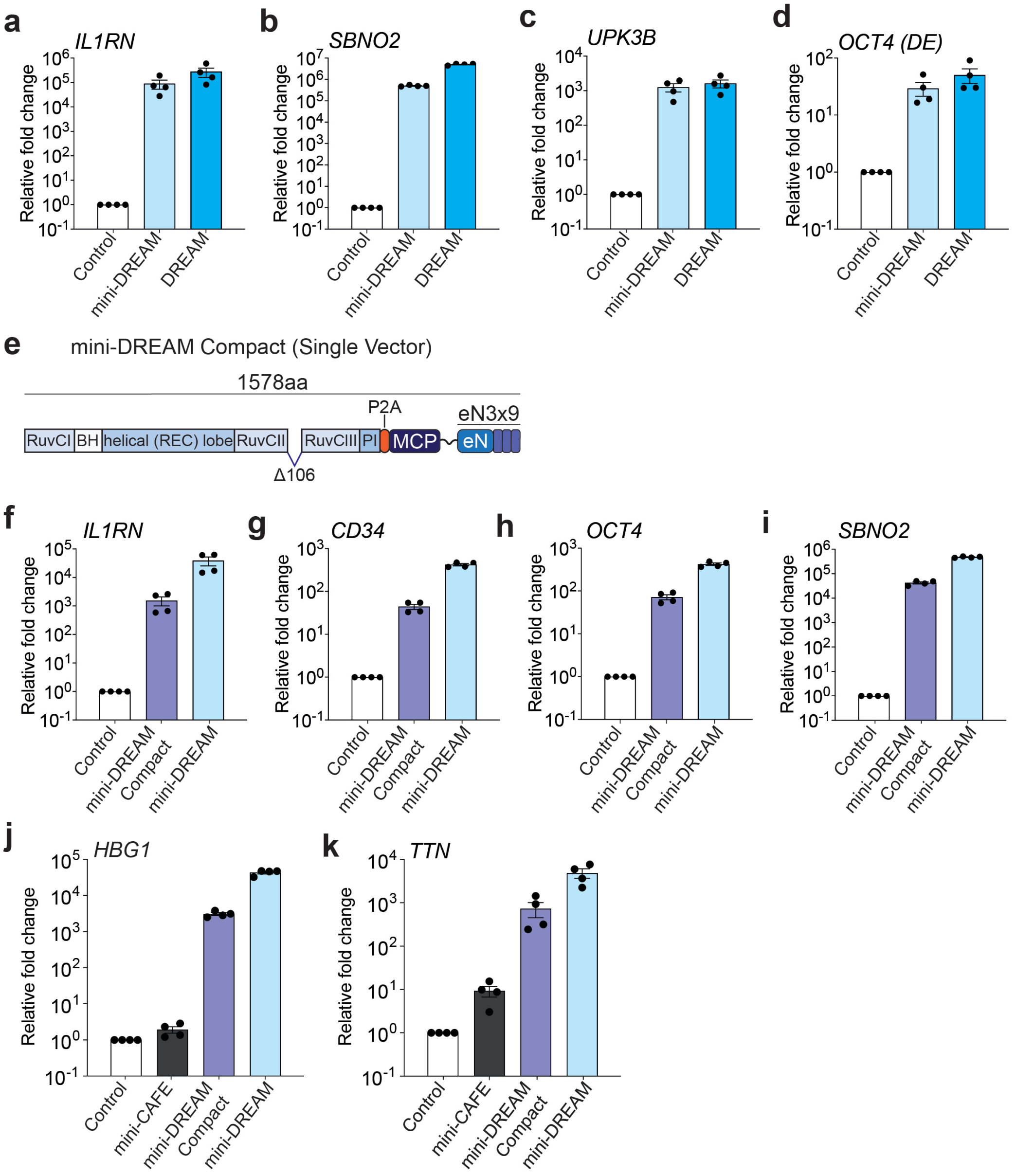


**Supplementary Fig. 23. mini-DREAM and mini-DREAM Compact systems display robust transactivation potencies in HEK293T cells. a – d.** Transactivation potencies of mini-DREAM and CRISPR-DREAM systems are shown when targeted to the *IL1RN* promoter (**panel a**) using pooled MS2-modified gRNAs, the *SBNO2* (**panel b**) or *UPK3B* promoters (**panel c**) using a single MS2-modified gRNA, respectively, and the *OCT4* distal enhancer (DE; **panel d**) using pooled MS2-modified gRNAs. **e.** The mini-DREAM Compact system is schematically depicted, P2A; self-cleaving peptide. **f – i.** Transactivation potencies of mini-DREAM and mini-DREAM Compact systems are shown when targeted to the *IL1RN* (**panel f**)*, CD34* (**panel g**) or *OCT4* (**panel h**) respectively, using pooled MS2-modified gRNAs, or to the *SBNO2* promoter using a single MS2-

modified gRNA (**panel i**). **j – k.** *HBG1* (left) or *TTN* (right) gene activation when either the CjdCas9 based mini-CAFE, SpdCas9 based mini-DREAM Compact or mini-DREAM system was targeted to each corresponding promoter using a pool of 4 MS2-modified gRNAs, respectively. All samples were processed for QPCR 72 hours post-transfection. Data are the result of at least 3 biological replicates. Error bars; SEM. *; $P < 0.05$, **; $P < 0.01$, ***; $P < 0.001$. *ns*; not significant.

#### Supplementary Figure 24

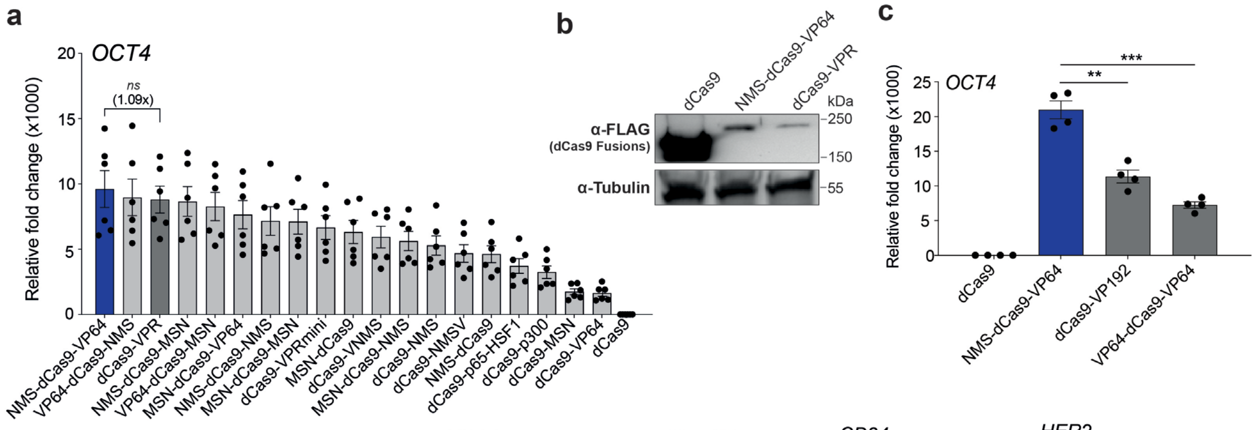


**d**

| dCas9 Fusion | N-Term | C-term | Total |
|---|---|---|---|
| NMS-dCas9-VP64 | 870 bp | 150 bp | 1020 bp |
| VP64-dCas9-NMS | 150 bp | 870 bp | 1020 bp |
| dCas9-VPR | | 1569 bp | 1569 bp |
| NMS-dCas9-MSN | 870 bp | 870 bp | 1740 bp |
| VP64-dCas9-MSN | 150 bp | 870 bp | 1020 bp |
| MSN-dCas9-VP64 | 870 bp | 150 bp | 1020 bp |
| NMS-dCas9-NMS | 870 bp | 870 bp | 1740 bp |
| MSN-dCas9-MSN | 870 bp | 870 bp | 1740 bp |
| dCas9-VPRmini | | 894 bp | 894 bp |
| MSN-dCas9 | 870 bp | | 870 bp |
| dCas9-VNMS | | 1026 bp | 1026 bp |
| MSN-dCas9-NMS | 870 bp | 870 bp | 1740 bp |
| dCas9-NMS | | 870 bp | 870 bp |
| dCas9-NMSV | | 1026 bp | 1026 bp |
| NMS-dCas9 | 870 bp | | 870 bp |
| dCas9-p65-HSF1 | | 939 bp | 939 bp |
| dCas9-p300-core | | 1851 bp | 1851 bp |
| dCas9-MSN | | 870 bp | 870 bp |
| dCas9-VP64 | | 150 bp | 150 bp |
| VP64-dCas9-VP64 | 150 bp | 150 bp | 300 bp |
| dCas9-VP192 | | 597 bp | 597 bp |

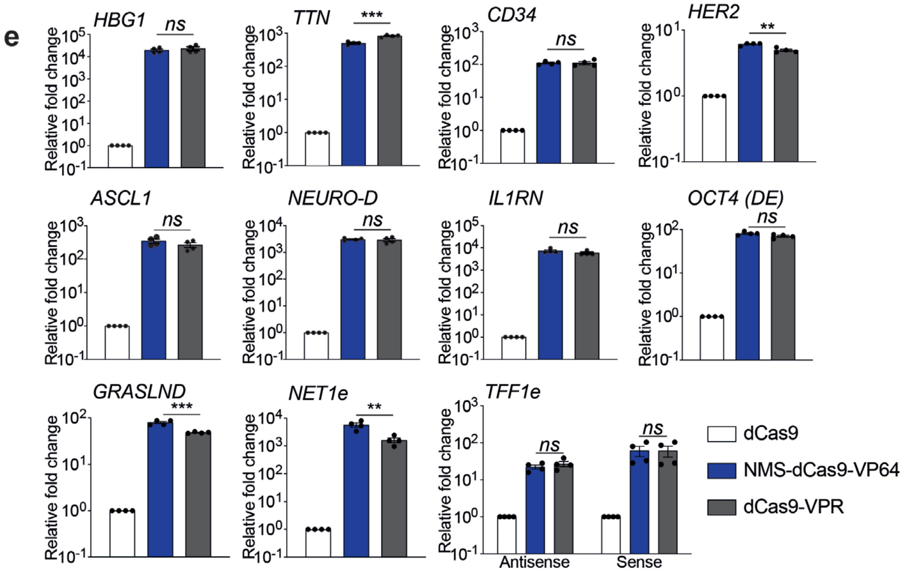


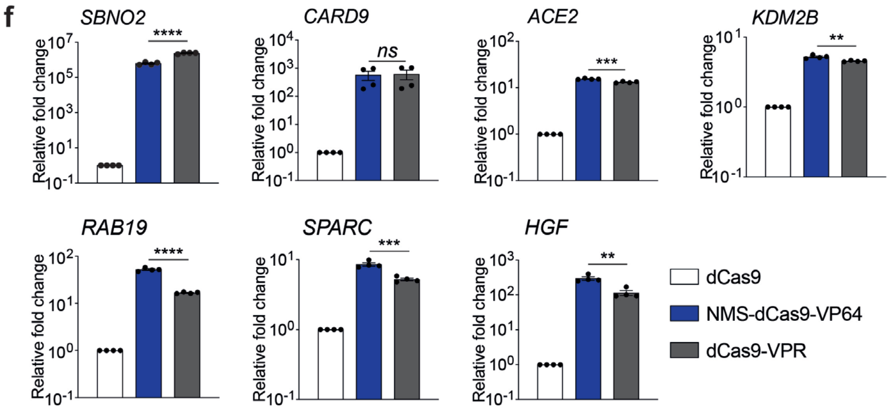


**Supplementary Fig. 24. Generating and validating tripartite TADs in direct fusion architectures. a.** *OCT4* mRNA levels after different dCas9 direct fusions were targeted to the *OCT4* promoter using pooled gRNAs. Indicated direct fusions were generated by linking MSN or NMS domains to either the C-terminus, N-terminus, or both termini of dCas9, along with selected combinations also containing VP64 as indicated. **b.** The expression levels of dCas9, NMS-dCas9-VP64, dCas9-VPR are shown as detected by Western blotting in HEK293T cells 72 hours post-transfection. **c.** *OCT4* mRNA levels after NMS-dCas9-VP4 and other VP64-based dCas9 fusion (VP64-dCas9-VP64 and dCas9-VP192) were targeted to the *OCT4* promoter using pooled gRNAs. **d.** Lengths (in bp) of different fusion proteins and modules are shown. **e.** Relative expression levels of 11 different endogenous human genes after dCas9, NMS-dCas9-VP64, or dCas9-VPR systems were targeted to their respective promoters/enhancers using pooled gRNAs. **f.** Relative expression levels of 7 different endogenous human genes after dCas9, NMS-dCas9-VP64, or dCas9-VPR systems were targeted to their respective promoters/enhancers using single gRNAs. All samples were processed for QPCR analysis 72 – 84 hours post-transfection in HEK293T cells and are the result of at least 4 biological replicates. See source data for more information. Error bars; SEM. *; $P < 0.05$, **; $P < 0.01$, ***; $P < 0.001$. *ns*; not significant.

#### Supplementary Figure 25

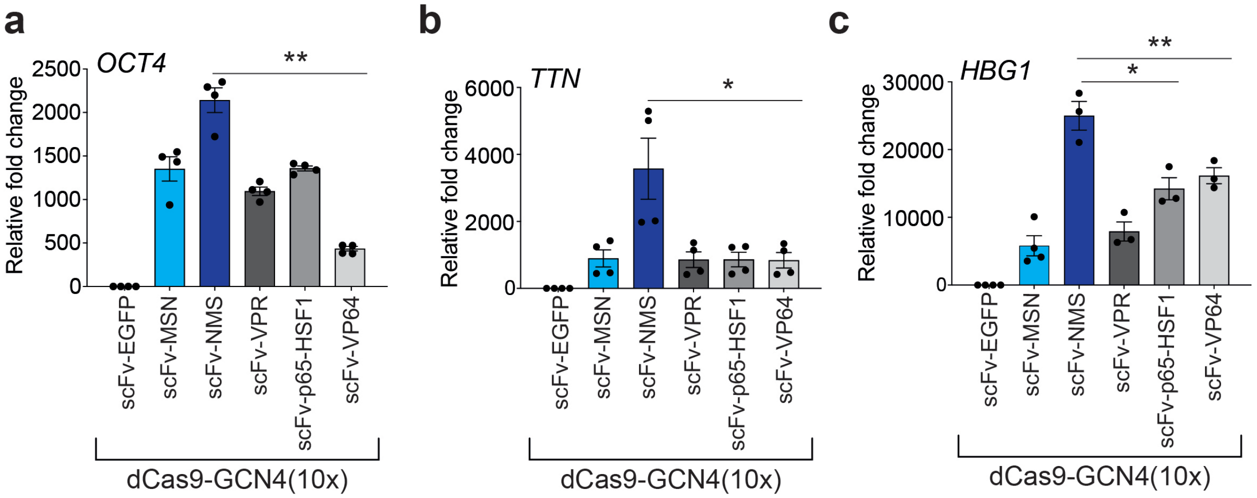


**Supplementary Fig. 25. The NMS effector domain is compatible with, and robust in, the dCas9 SunTag system. a – c.** *OCT4* (**panel a**), *TTN* (**panel b**) and *HBG1* (**panel c**) mRNA levels after the indicated TADs were fused to scFv and recruited via dCas9 harboring a 10xGCN4 C-terminal fusion protein (the SunTag system) along with 4 pooled gRNAs targeting each respective promoter. All samples were processed for QPCR analysis 72 hours post-transfection in HEK293T cells and are the result of at least 3 biological replicates. See source data for more information. Error bars; SEM. *; $P < 0.05$, **; $P < 0.01$. *ns*; not significant.

**Supplementary Figure 26**

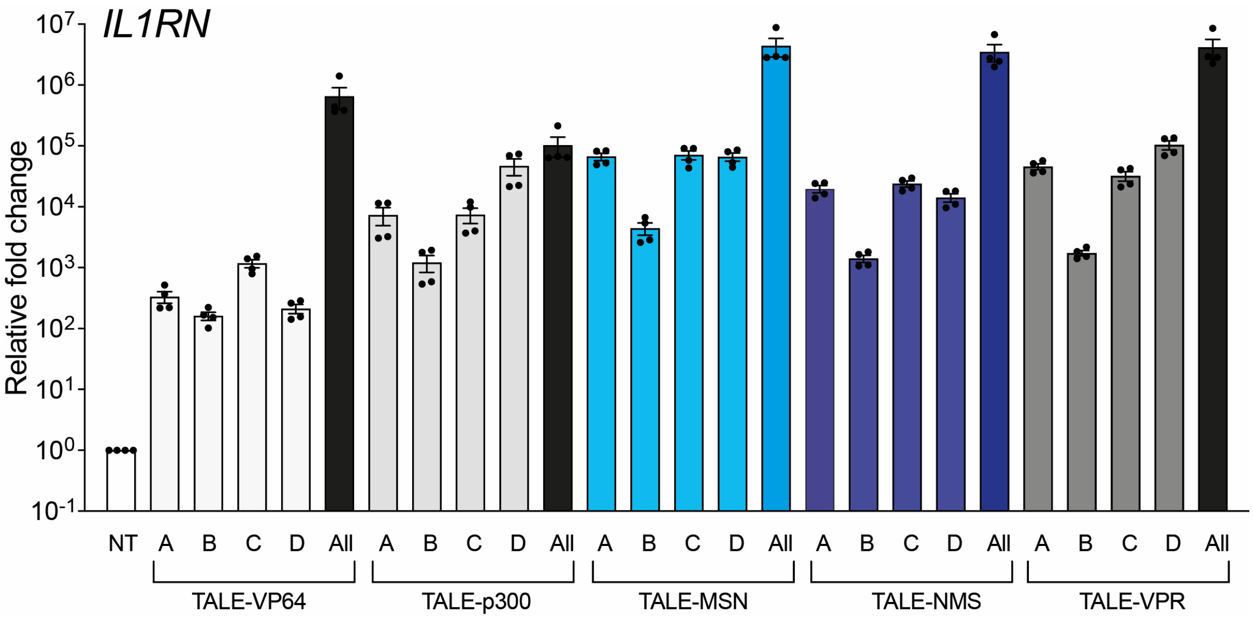


**Supplementary Fig. 26. The tripartite MSN and NMS TADs are portable to synthetic TALE DNA binding systems. a.** Relative *IL1RN* mRNA levels after individual (A – D) or pooled ("all") *IL1RN* promoter-targeting TALEs (TALE-VP64, TALE-p300, TALE-MSN, TALE-NMS and TALE-VPR) were co-transfected into HEK293T cells. All samples were processed for QPCR analysis 72 hours post-transfection in HEK293T cells and are the result of at least 4 biological replicates. Error bars; SEM.

#### Supplementary Figure 27

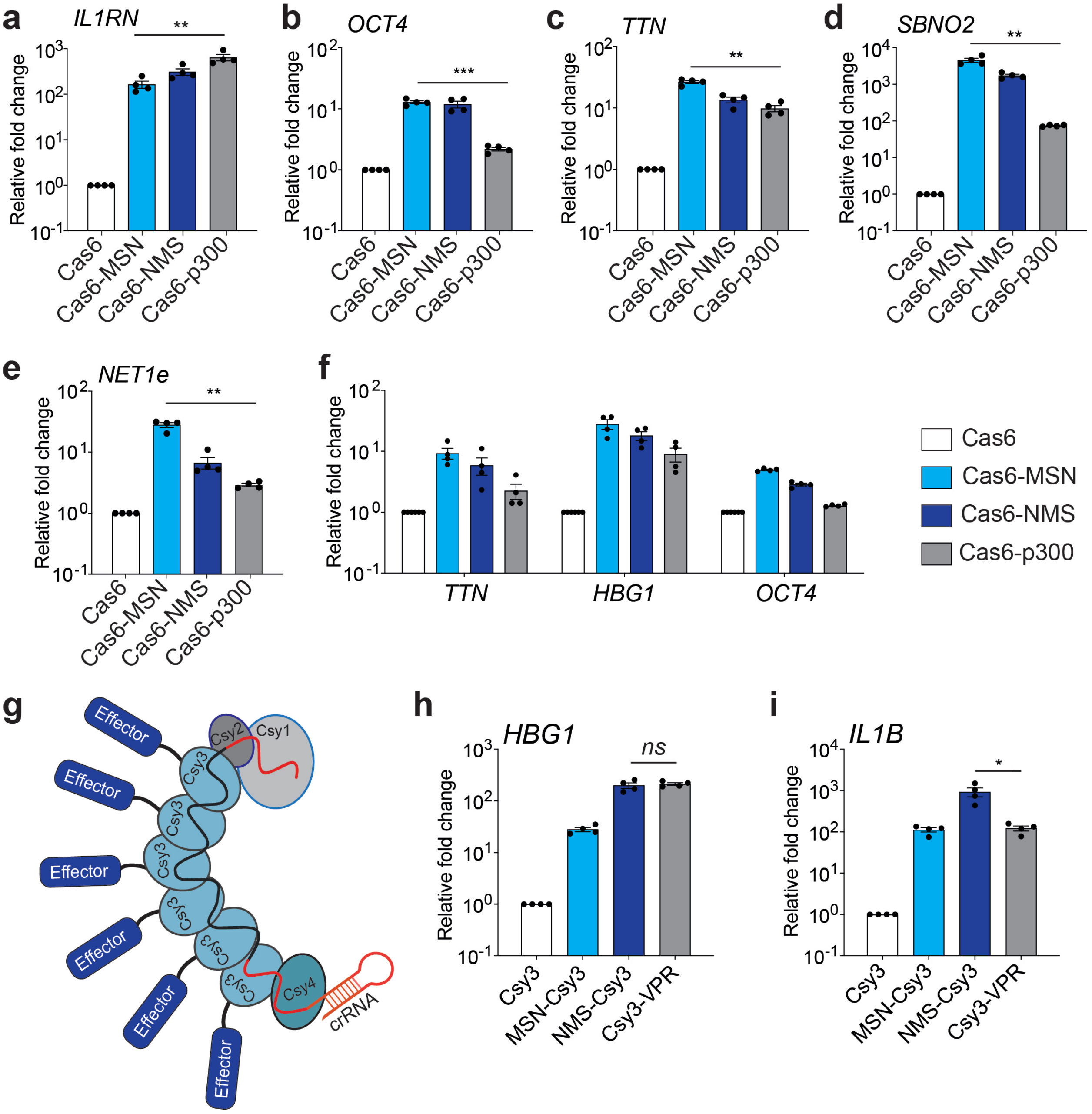


**Supplementary Fig. 27. The tripartite MSN and NMS TADs are portable to different Type I CRISPRa systems. a – d.** *IL1RN* (**panel a**), *OCT4* (**panel b**), *TTN* (**panel c**) or *SBNO2* (**panel d**) mRNA activation when the indicated Cas6 fusion protein encoding plasmids (and plasmids encoding the other components of Eco-cascade complex) were targeted to each corresponding promoter using a single crRNA. **e.** *NET1 eRNA* activation when the indicated Cas6 fusion protein encoding plasmids (and plasmids encoding the other components of Eco-cascade complex) were targeted to respective enhancer using a single crRNA. **f.** Multiplexed activation of 3 protein coding genes (*TTN*, *HBG1* and

*OCT4*) following co-transfection of the indicated Cas6 fusion protein encoding plasmids (and plasmids encoding the other components of Eco-cascade complex) and a multiplexed crRNA expression plasmid encoding 1 crRNA/locus. **g.** The Type I CRISPR system derived from *P. aeruginosa* (*Pae*-Cascade) is schematically depicted along with a representative effector fused to the Csy3 protein subunit. **h – i.** *HBG1* (**panel h**) and *IL1B* (**panel i)** mRNA activation using the indicated Csy3 fusion proteins when targeted to each corresponding promoter using a single crRNA, respectively. All samples were processed for QPCR analysis 72 hours post-transfection in HEK293T cells and are the result of at least 4 biological replicates. See source data for more information. Error bars; SEM. *; $P < 0.05$, **; $P < 0.01$, ***; $P < 0.001$. *ns*; not significant.

#### Supplementary Figure 28

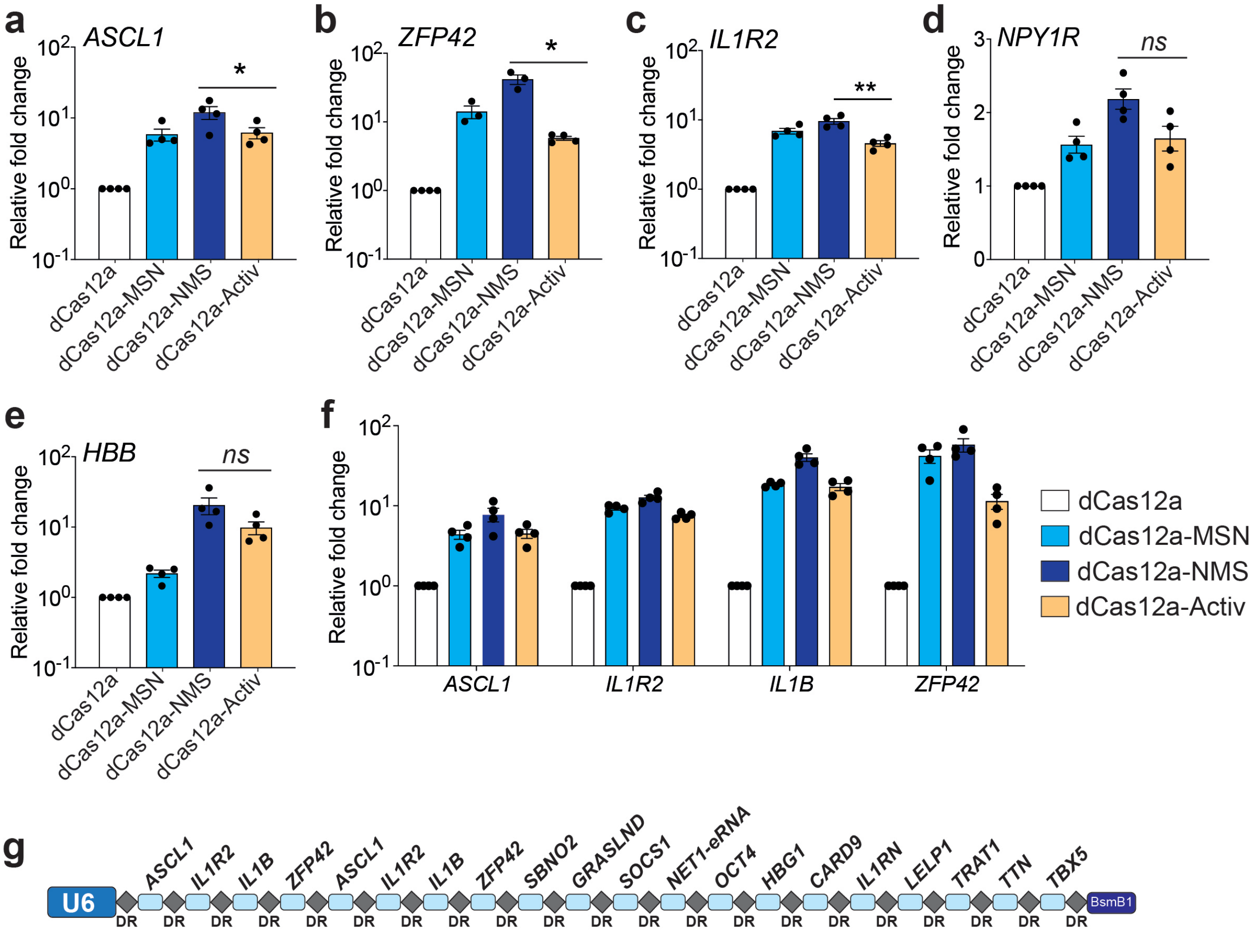


**Supplementary Fig. 28. The tripartite MSN and NMS TADs are portable to the dCas12a-based CRISPRa system. a – c.** *ASCL1* (**panel a**), *ZFP42/REX1* (**panel b**) *and IL1R2* (**panel c**) transactivation after the indicated dCas12a fusion proteins were targeted to each corresponding promoter using 2 crRNAs per respective locus. **d – e.** *NPY1R* (**panel d**) and *HBB* (**panel e**) mRNA activation after the indicated dCas12a fusion proteins were targeted to each corresponding promoter using a single crRNA. **f**. Multiplexed activation of 4 indicated endogenous genes 72 hours after co-transfection of indicated dCas12a fusion protein encoding plasmids and a single plasmid encoding an 8-crRNA expression array (2 crRNAs/gene promoter). **g**. The crRNA expression array encoding 20 crRNA targeting 16 loci used in main text **Figure 4i**, is schematically depicted. All samples were processed for QPCR analysis 72 hours post-transfection in HEK293T cells and are the result of at least 3 biological replicates. See source data for more information. Error bars; SEM. *; $P < 0.05$. *ns*; not significant.

#### Supplementary Figure 29

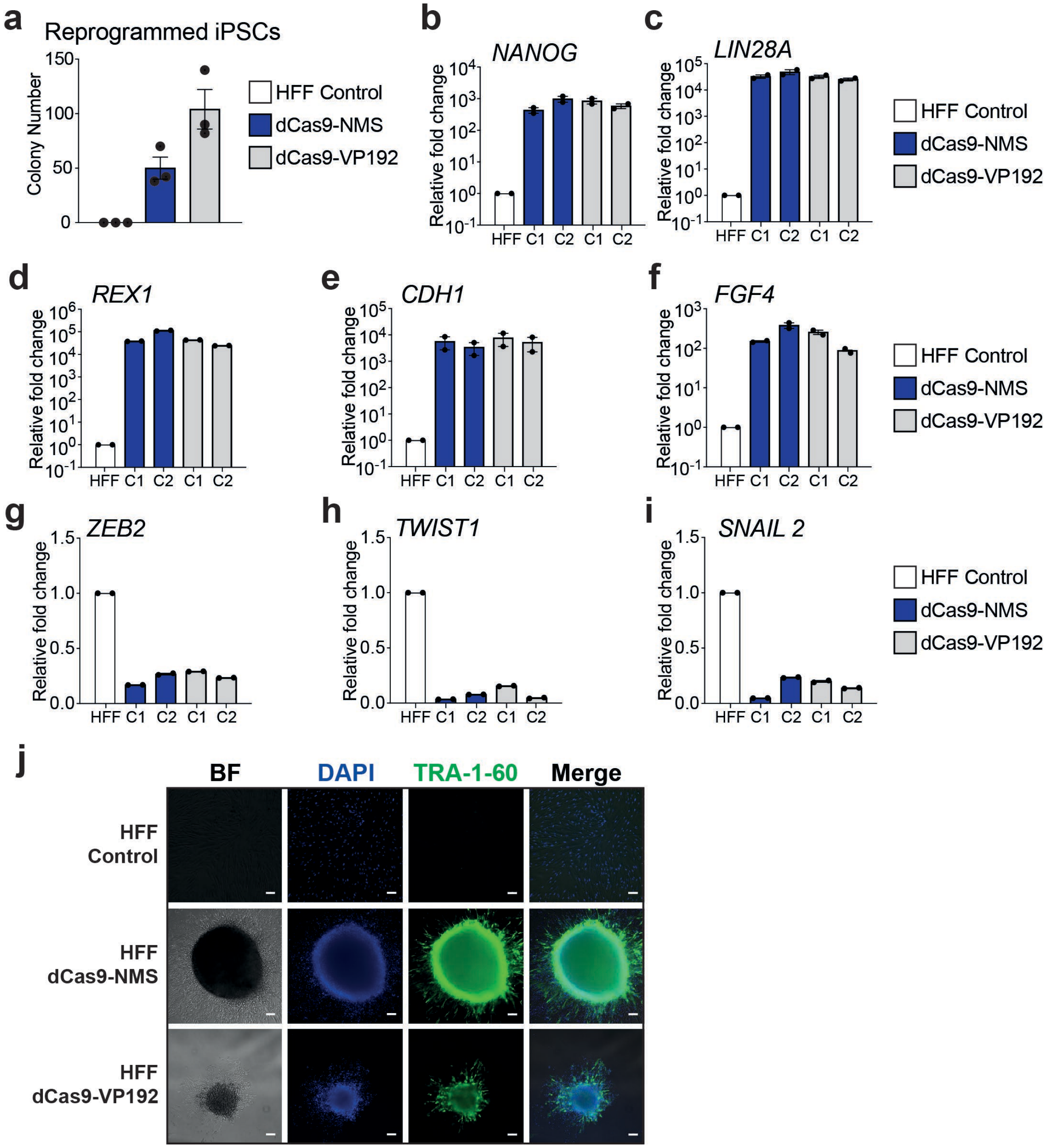


**Supplementary Fig. 29. dCas9-NMS permits efficient *in vitro* reprogramming of human fibroblasts. a.** Bar graph showing iPSC colonies (the total number of colonies per million HFFs) generated from HFFs nucleofected with either dCas9-NMS and dCas9-VP192 and a gRNA cocktail (see main text and methods). **b – f.** Relative expression of pluripotency-associated genes *NANOG* (**panel b**), *LIN28A* (**panel c**), *REX1* (**panel d**), *CDH1* (**panel e**), *FGF4* (**panel f**) in representative colonies (C1 or C2) ~40 days after nucleofection of either dCas9-NMS (blue) or dCas9-VP192 (gray) and multiplexed gRNAs compared to untreated HFF controls. **g – i.** Relative expression of mesenchymalassociated genes *ZEB2* (**panel g**), *TWIST1* (**panel h**) and *SNAIL2* (**panel i**), in representative colonies (C1 or C2) ~40 days after nucleofection of either dCas9-NMS (blue) or dCas9-VP192 (gray) and multiplexed gRNAs compared to untreated HFF controls. **j.** Immunofluorescence microscopy of HFFs ~40 days after nucleofection of either dCas9-NMS or dCas9-VP192 and multiplexed gRNAs compared to untreated HFF controls (white scale bars, 100μm). Cells were immunostained for the expression of pluripotency-associated cell surface marker TRA-1-60 (**panel j**, green). All cells were counterstained with DAPI to visualize the nucleus.

#### Supplementary Figure 30

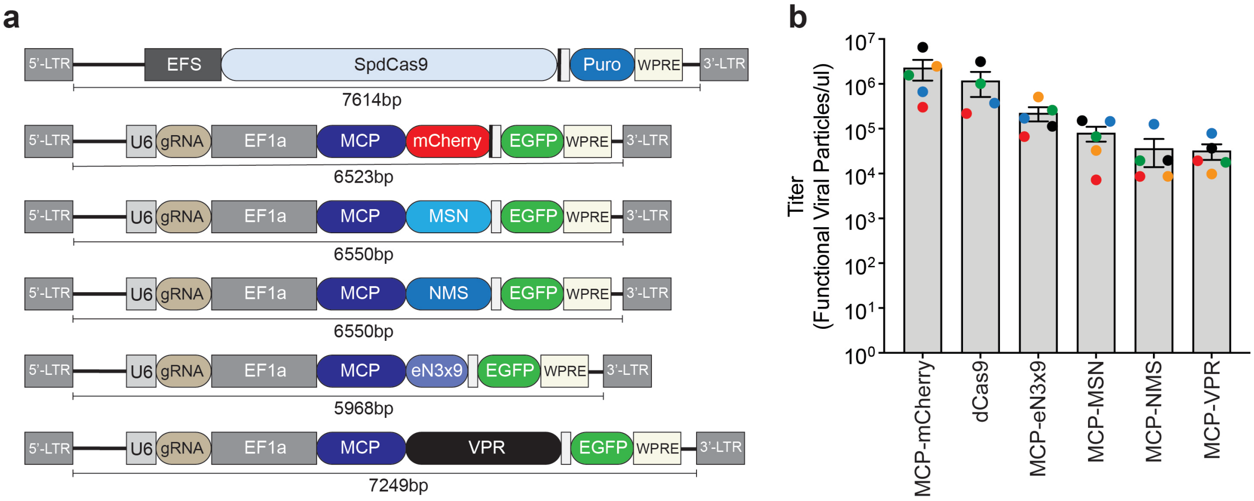


**Supplementary Fig. 30. Lentiviral titers are influenced by the presence of TAD domains. a**. Schematic representation of different lentiviral vectors with their respective sizes shown (in bp). **b**. Titers of different lentiviruses used in this study. Each colored dot indicates a lentiviral titer from an independent preparation.

#### Supplementary Figure 31

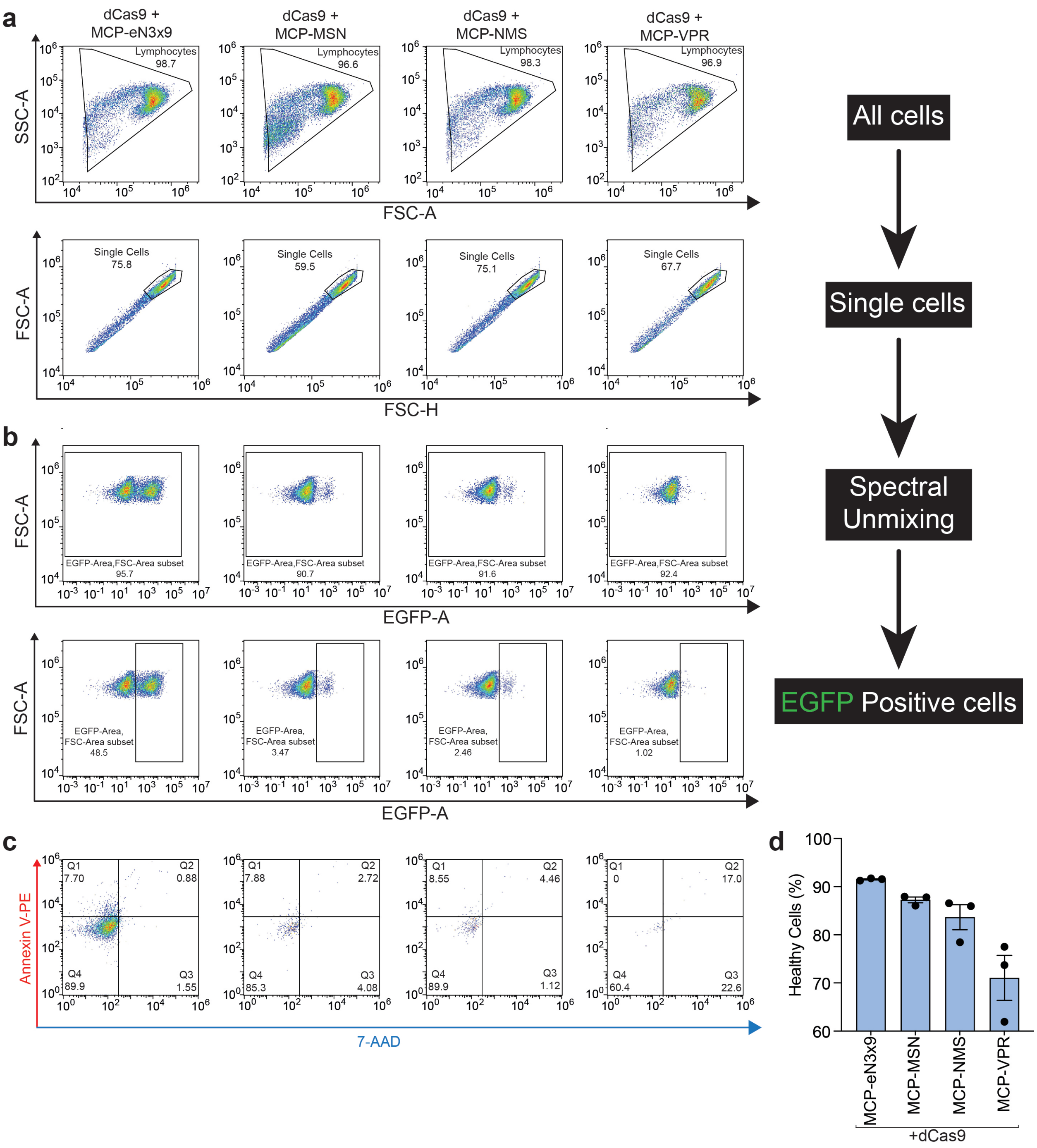


**Supplementary Fig. 31. Engineered TADs from MTFs exhibit lower toxicity compared to VPR in primary T cells. a.** Density plots indicating gating for bulk (top) and single (bottom) primary human T cells after cells were transduced with dCas9 along with MCP-eN3x9 (column 1), MCP-MSN (column 2), MCP-NMS (column 3), or MCP-VPR (column 4). All MCP-fusion encoding vectors also encode an MS2-modifed gRNA targeting human *TTN*. **b.** Density plots indicating the gating strategy to remove events with EGFP signal < $1x10^{-3}$ resulting from spectral unmixing (top) and to select EGFP positive cells (bottom). **c.** Flow cytometry plot showing $7AAD^{+}$/Annexin $V^{+}$ cells (after enriching for EGFP positive cells) co-transduced with indicated lentiviral vectors. **d**. Bar graph showing the percentage of healthy primary T cells among the EGFP positive (from flow cytometry data) transduced with indicated lentiviral particles. All experiments were performed in triplicate and density plots shown are representative of all replicates. See source data for more information.

#### Supplementary Figure 32

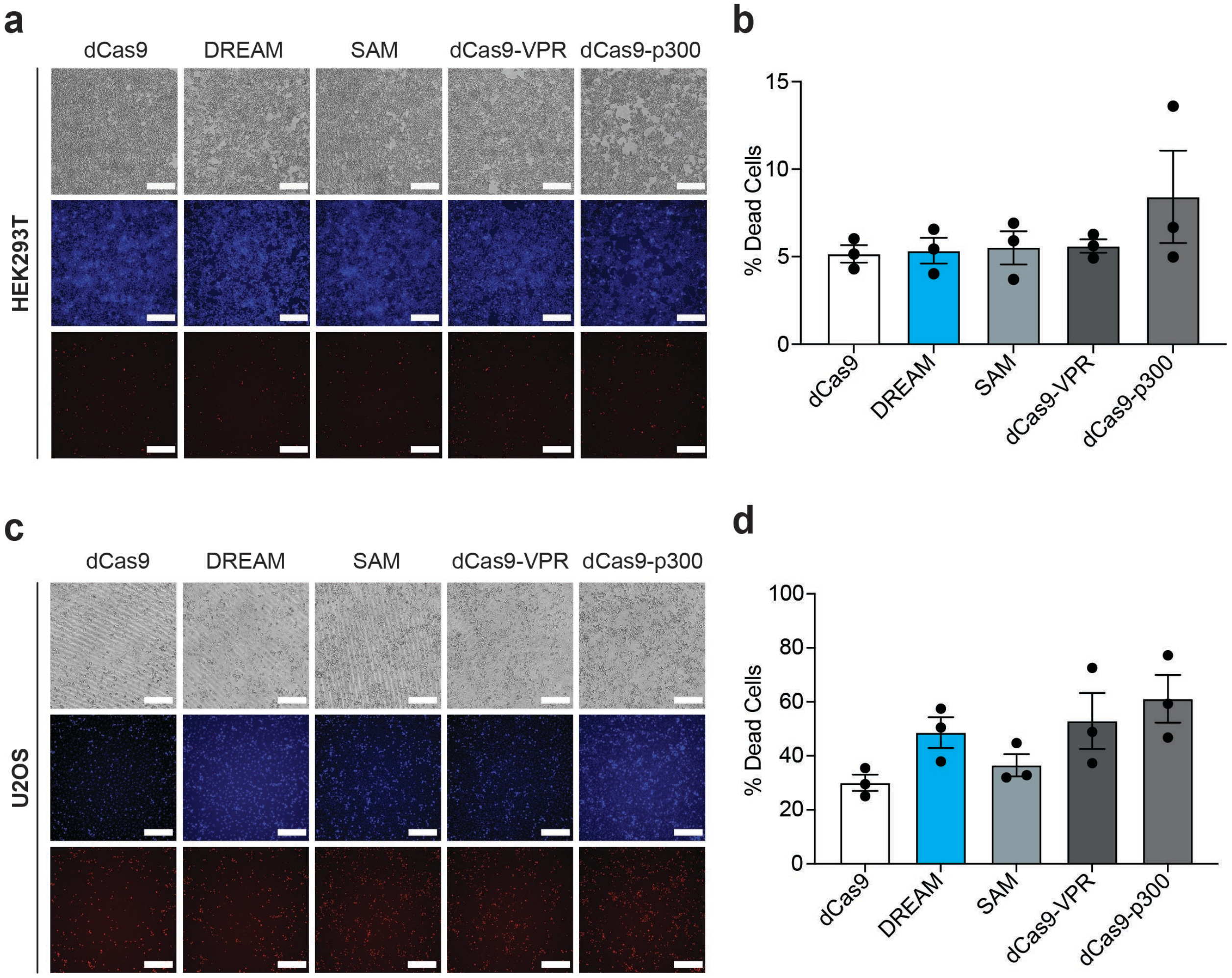


**Supplementary Fig. 32. Comparison of toxicity of different CRISPRa platforms in cultured adherent cell lines. a.** Microscopy images showing brightfield, Hoechst, and 7-AAD staining of HEK293T cells 48 hours post-transfection of indicated CRISPRa tools in presence of 4 pooled gRNAs (MS2 modified for DREAM and SAM systems) targeting *HBG1* (white scale bars, 250μm). **b.** Bar graph showing percentage of dead cells obtained from quantification of microscopic images HEK293T cells. **c.** Microscopy images showing brightfield, Hoechst, and 7-AAD staining of U2OS cells 48 hours post-transfection of indicated CRISPRa tools in presence of 4 pooled gRNAs (MS2 modified for DREAM and SAM systems) targeting *HBG1* (white scale bars, 250μm). **d.** Bar graph showing percentage of dead cells obtained from quantification of microscopic images in U2OS cells.

**Supplementary Figure 33**

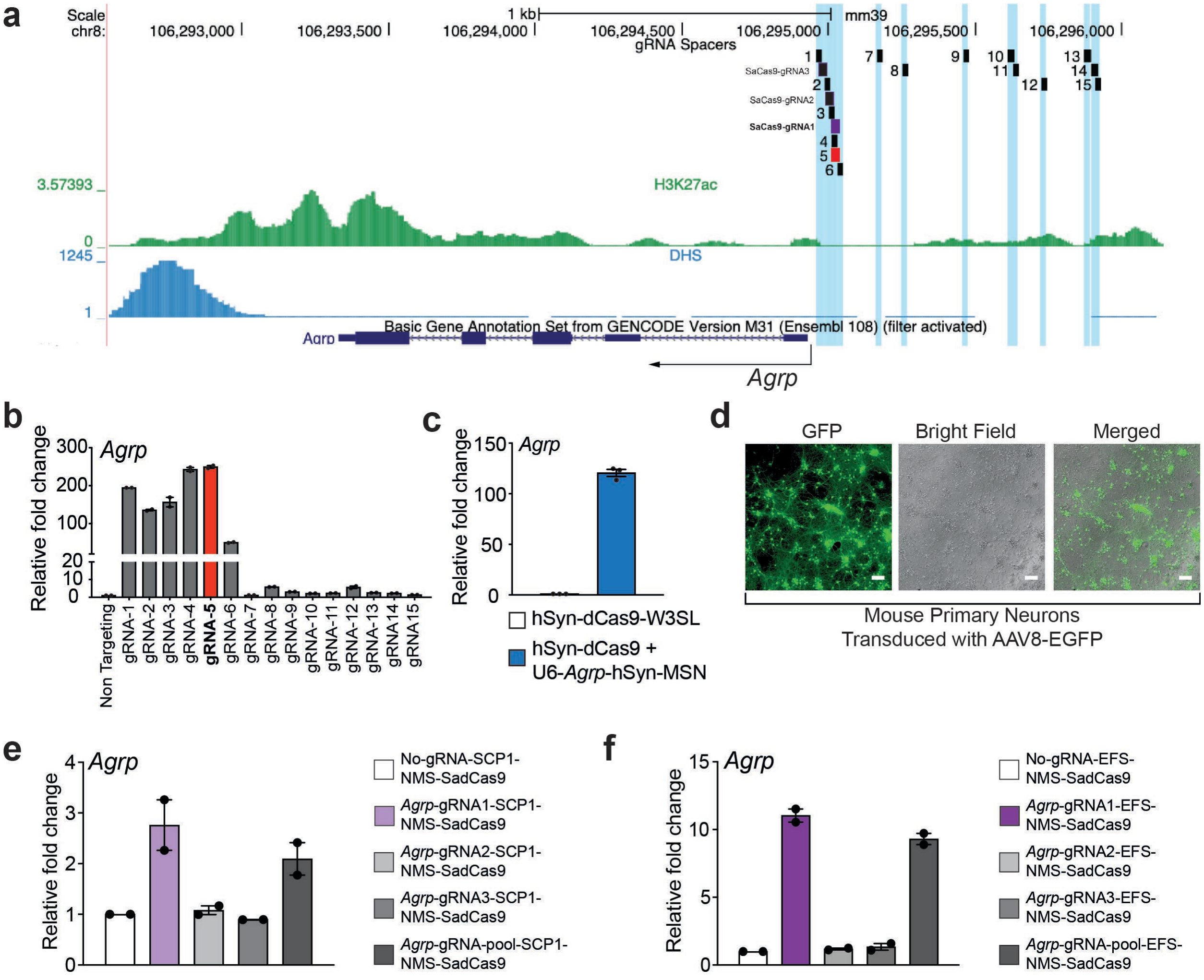


**Supplementary Fig. 33. Selection and validation of optimal *Agrp* gRNA and construct designs for dual and all-in-one (AIO) AAV vectors. a and b.** The genomic region (mm10) encompassing the mouse *Agrp* gene on chromosomes 8 is shown. *Agrp* gene is shown in dark blue; SpdCas9 gRNA (1-15) target regions are indicated by black lines and light blue highlighting except for the most potent gRNA (gRNA 5), which was selected for future experiments and is shown in red. SadCas9 gRNA (1-3) target regions are indicated by black lines and light blue highlighting except most potent gRNA1, which was selected for future experiments was shown in purple. H3K27ac (from GSE10666), and DNase Hypersensitivity Sites (DHSs; from GSE37074) are shown in green, and blue, respectively. Transcription Start Sites (TSSs) for *Agrp* is indicated by black arrows. **b**. Comparison of transactivation potency of different gRNA targeting *Agrp* promoter using the DREAM system. 15 gRNAs were designed to tile across ~1.5kb upstream of the *Agrp* promoter. Non-transfected Neuro-2A cells were used as a control. **c**. Dual AAV plasmid (comprising CRISPR-DREAM and *Agrp* gRNA5) mediated *Agrp* induction in Neuro-2A cells. **d**. Immunofluorescence microscopy showing EGFP expression in AAV8-EGFP transduced (2.5 X $10^4$ vg/ cell) mouse primary cortical neurons 72 hours post-transduction (white scale bars, 100μm). **e**. *Agrp* induction in Neuro-2A cells 72 hours after transfection of the AIO AAV construct consisting of the SCP1 promoter driven NMS-SadCas9. **f**. *Agrp* induction in Neuro-2A cells 72 hours after transfection of the AIO AAV construct consisting of EFS promoter driven NMS-SadCas9.

**Supplementary Figure 34**

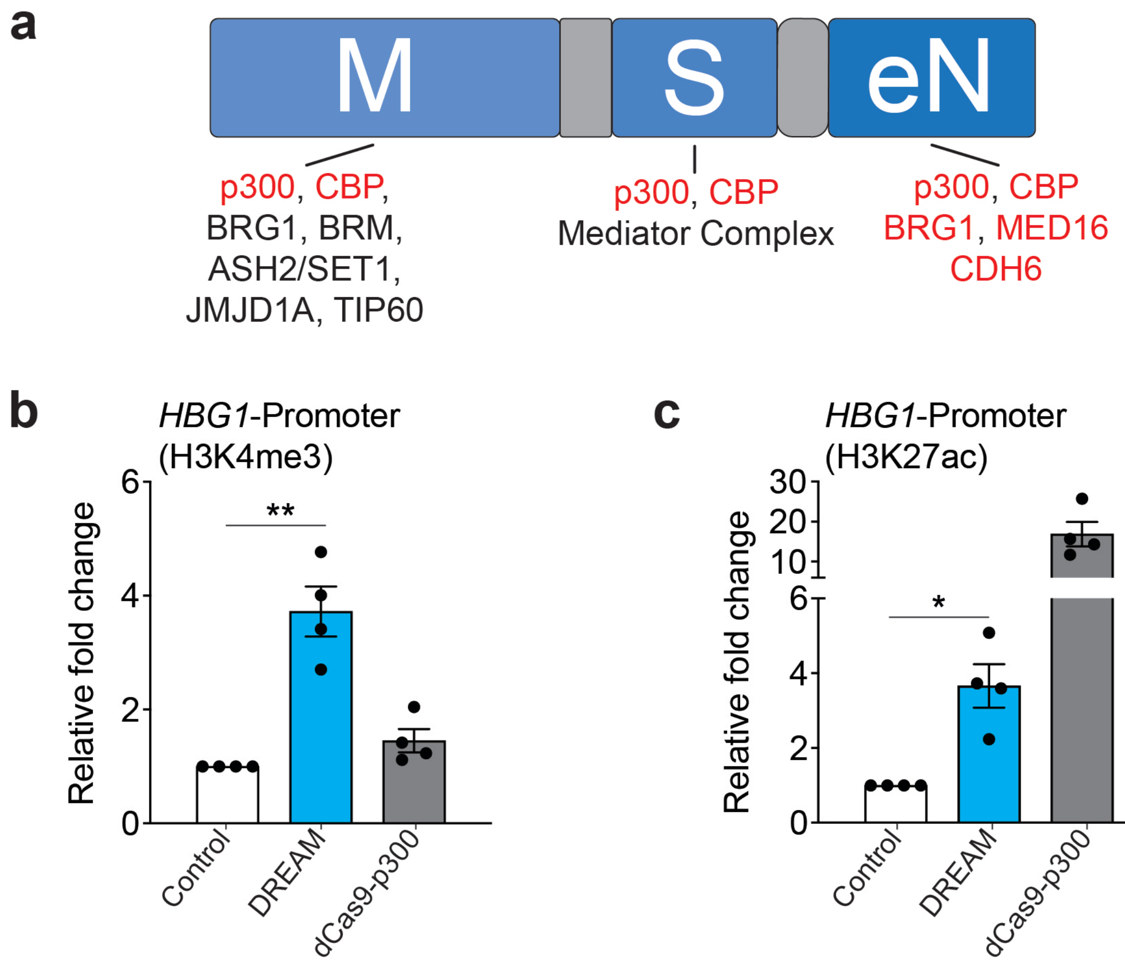


**Supplementary Fig. 34. DREAM deposits epigenetic marks related to transcriptional activation at a targeted promoter. a.** Schematic showing previously described interactions among different epigenetic modifiers/chromatin remodelers and MRTF-A, STAT1, and NRF2. Epigenetic modifiers/chromatin remodelers highlighted in red have been shown to interact directly with the TAD regions of MRTF-A, STAT1 and NRF2 (or eNRF2). Epigenetic modifiers/chromatin remodelers highlighted in black have been observed to interact with MRTF-A, STAT1, and NRF2, but interaction contacts are incompletely characterized to the best of our knowledge. **b and c.** Relative level of H3K4me3 and H3K27ac enrichment after control (dCas9 + MCP-mCherry), DREAM, or dCas9-p300 were targeted to the *HBG1* promoter using pools of 4 MS2-modified gRNA (control or DREAM) or pools of 4 standard gRNAs (dCas9-p300) in HEK293T cells. All samples were processed for CUT&RUN analysis 72 hours post-transfection and are the result of at least 3 biological replicates. See source data for more information. Error bars; SEM. *; $P < 0.05$, **; $P < 0.01$.

**Supplementary Figure 35**

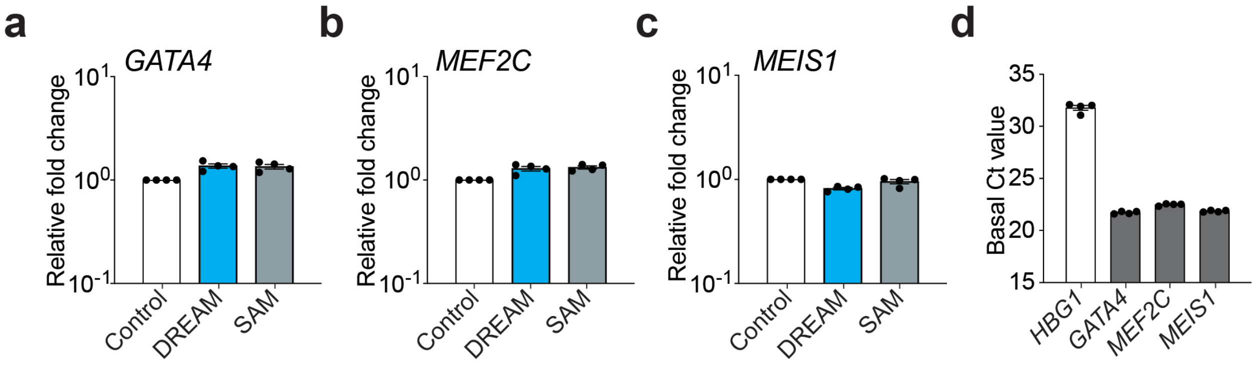


**Supplementary Fig. 35. DREAM and SAM systems fail to activate cardiomyocyte reprogramming genes in HEK293T cells. a – c.** Relative expression of *GATA4, MEF2C and MEIS1* after dCas9 control, DREAM, or SAM systems were targeted to their respective promoters using a single MS2-modifed gRNAs in HEK293T cells. **d.** Relative basal expression of *HBG1* (a relatively lowly expressed gene), *GATA4*, *MEF2C*, and *MEIS1* (relatively highly expressed genes) in HEK293T cells. Raw cycle threshold (Ct) values are shown as a proxy for expression levels. All samples were processed for QPCR analysis 72 hours post-transfection and are the result of at least 4 biological replicates. Error bars; SEM.
